# An Adenine-Based Molecular Rotor as a Universal Fluorescent Nucleobase with High Brightness

**DOI:** 10.64898/2026.01.18.700143

**Authors:** Anna A. Pushkarevskaya, Polina N. Kamzeeva, Evgeny S. Belyaev, Vladimir A. Brylev, Alexander A. Lomzov, Andrey V. Aralov

**Author notes:** (AAL), (AVA).

## Abstract

Chemically modified nucleic acids have become a powerful platform for basic research and applied technologies. Universal nucleobases are used in PCR,sequencing, and the design of nanodevices and aptamers. Fluorescent universal nucleobases have an even wider range of applications, including the development of nucleic acid-based sensors, switches, and relay logic gates. However, few such nucleobases have been proposed to date, and most of them have suboptimal optical properties. Here, we propose an adenine-based molecular rotor, 7,8-dihydro-8-oxo-6-(3-methylbenzo[d]thiazol-2(3H)-ylidene)adenine (**oxo-Ade** ^**BZT**^), as a new, remarkably bright and potent fluorescent universal nucleobase. Its brightness in both oligodeoxyribonucleotides (ODNs) and DNA duplexes (4200 - 10000 M^-1^ × cm^-1^) originates from a high molar extinction coefficient (averaged *ε*_*368*_ 37000 M^-1^ × cm^-1^), provided by the appended 3-methylbenzo[d]thiazolyl moiety, and a relatively high quantum yield (0.11 – 0.27). Melting temperature variations observed upon the incorporation of **oxo-Ade** ^**BZT**^ opposite native nucleobases in a duplex context did not exceed 10%. The basis of these universal hybridizing properties was unveiled using computational methods. According to molecular dynamics simulations, **oxo-Ade** ^**BZT**^ pushes the opposite nucleobase out of the DNA double helix and forms multiple hydrophobic contacts with the flanking base pairs. At the same time, the rotational mobility of the bonds between the **oxo-Ade** ^**BZT**^-constituting heterobicycles decreases, and **oxo-Ade** ^**BZT**^ adopts a planar conformation in both ODNs and their duplexes, resulting in the light-up effect. These properties make **oxo-Ade** ^**BZT**^ a promising molecular tool for analytical, biophysical and biochemical studies.

## Introduction

Fluorescent nucleobases are powerful tools for investigating the structures, dynamics, localization, and binding interactions of nucleic acids^1^. Unlike intercalating dyes and post-synthetically conjugated fluorophores, fluorescent nucleobases can be precisely positioned with high precision within nucleic acid sequences. Fluorescent nucleobases can be divided into two categories: canonical and non-canonical. The former can preserve the standard purine/pyrimidine base pair architecture and regular helix geometry but impose limitations on structural design. Consequently, canonical fluorescent nucleobases usually exhibit poor or suboptimal photophysical properties. Non-canonical fluorescent nucleobases are not designed to exhibit specific base pairing. Thus, they allow wide variability in molecular sizes and shapes, providing strong π-stacking interactions and tunability of spectral properties.

Adenine derivatives are particularly prevalent category among canonical and non-canonical fluorescent nucleobases. The first adenine modifications proposed were its isomer, 2-aminopurine (**2AP**^2^), which has poor discriminating ability towards opposing pyrimidines, and tricyclic 1,*N*^6^-ethenoadenine (**εA**^3^), which is unable to pair with nucleobases (Fig. 1A). Despite their poor spectral properties, both are of great use in nucleic acid research^4,5^. To date, a large number of canonical fluorescent adenine analogs have been proposed. Notable examples include **8vA**^6^, which has a vinyl group at the 8-position, and **A**^**PY** 7^ and **Py⍰A**^8^, which have a pyrene residue appended at the 8-or 2-position, respectively. These analogs possess prospective fluorescent properties but are mostly quenched upon duplex formation. Another subclass includes the tetracyclic derivatives **qA**^9^ and **qAN1**^10^, as well as the brightest representative of this class, **2CNqA**^11^ (Fig. 1A). These analogs have been successfully used to construct FRET pairs for DNA and RNA conformation studies^11,12^ and the preparation of fluorescent gapmers for in-cell imaging^13^. Importantly, **2CNqA** monophosphate and triphosphate demonstrated spontaneous cellular uptake and could be incorporated into *de novo* synthesized cellular RNA. This opened the way for metabolic fluorescent RNA labeling and emphasized the importance of developing fluorescent nucleotides for research and therapeutic purposes^14^ (Fig. 1A). Finally, based on crystal structure analysis indicating the tolerability of modification at the 6-position of adenine by bacterial DNA glycosylase, the adenine analog **A4**, which contains a 5-(methylthio)thiophen-2-yl moiety instead of the exocyclic amino group, was developed for real-time monitoring of enzymatic base excision activity *in vitro*^15^ (Fig. 1A).

**Fig. 1.**
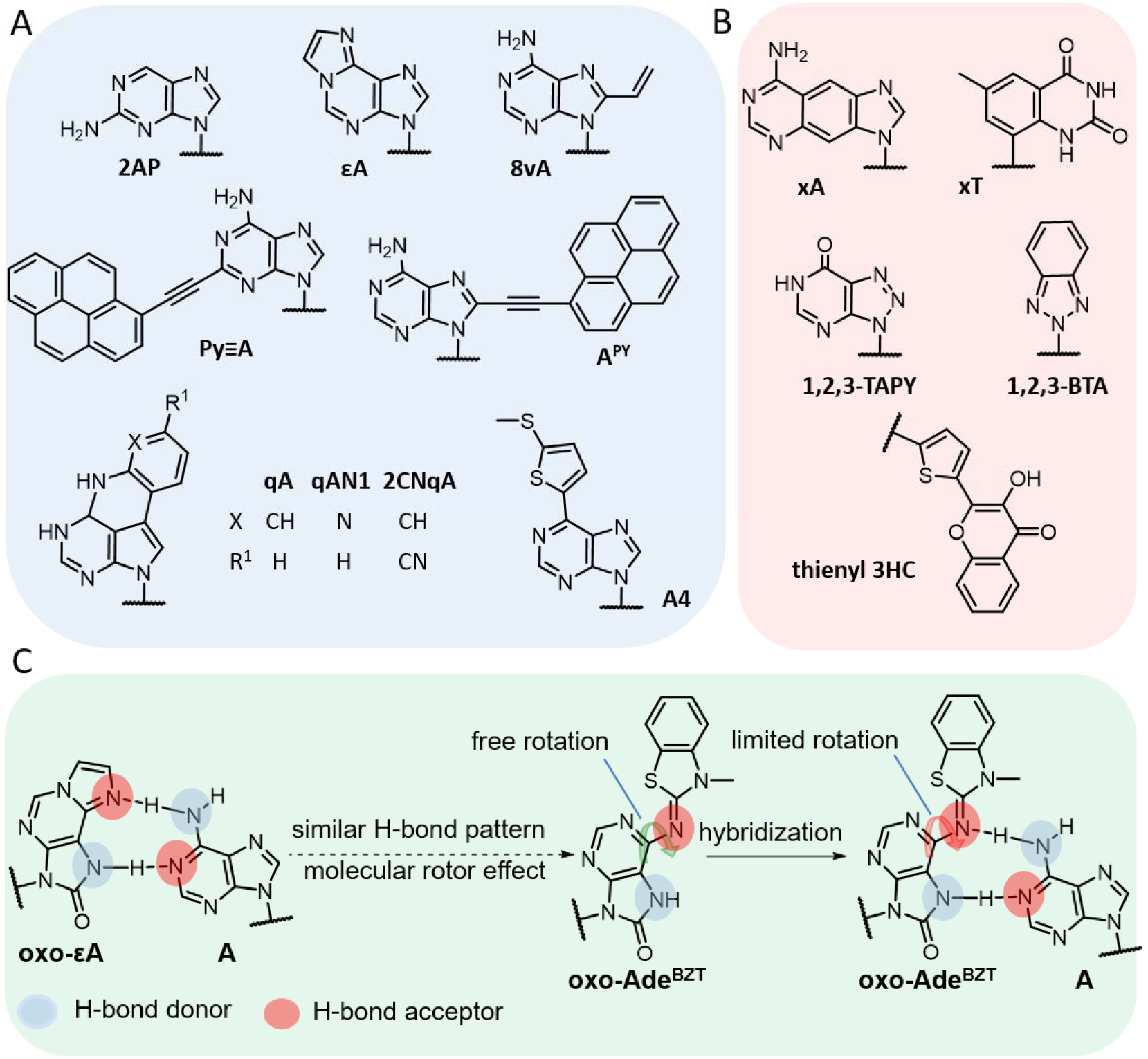
Fluorescent adenine analogs and derivatives (A), fluorescent universal nucleobases (B) and the design of the **oxo-Ade**^**BZT**^ modification based on initial hypothesis, retaining the **oxo-εA** H-bonding pattern and introducing the molecular rotor effect (C).

Universal nucleobase analogs are an attractive class of modifications that could advance DNA hybridization-based techniques such as PCR methods with degenerate primers and randomized sequencing, and DNA-based nanodevices^16^. Recently, they have also found applications in binding site mapping and generating high-affinity aptamers^17–20^. A universal nucleobase must pair equally with natural nucleobases (A, T/U, G, C) without impeding enzymatic DNA/RNA synthesis by polymerase, and the corresponding triphosphate must serve as a polymerase substrate and be recognized by intracellular enzymes^21^. Despite numerous attempts to synthesize a universal nucleobase, none of the reported analogs fulfils all the criteria. Bright and stable fluorescence provides an extra modality for the development of aptamer switches or FRET relay logic gates and is particularly useful in sensor design^17^. To date, only a few fluorescent universal nucleobases have been developed. Prominent examples include **xA** and **xT**; however, they are substantially quenched upon duplex formation^22^. Other examples include 1,2,3-benzotriazole (**1**,**2**,**3-BTA**) and 1,2,3-triazolo[4,5-d]pyrimidine (**1**,**2**,**3-TAPY**), which have low brightness^23^, and thienyl 3-hydroxychromone (**thienyl 3HC**), which has a fluorescence quantum yield (*Φ*_*f*_,) close to zero when flanked by cytosines and guanines in DNA duplexes^24^ (Fig. 1B).

In search of a bright adenine derivative with stable fluorescence in oligodeoxyribonucleotides (ODNs) and their DNA duplexes, we designed a fluorescent **oxo-Ade** ^**BZT**^ nucleoside. We explored the exceptionally high molar extinction coefficient of the appended 3-methylbenzo[d]thiazolyl moiety and studied the ability of the constructed modification to act as a molecular rotor. After confirming the promising photophysical properties of the **oxo-Ade** ^**BZT**^ nucleoside, we further studied its fluorescent properties and its impact on thermal stability in a duplex context. We discovered its universal nature and its stable, bright, and flank-sensitive fluorescence. Finally, molecular modeling helped clarify the basis of the observed increase in fluorescence intensity when moving from the **oxo-Ade** ^**BZT**^ nucleoside to modified ODNs and their complexes and the ability of **oxo-Ade** ^**BZT**^ to serve as a nucleobase surrogate with universal hybridizing properties.

## Results and Discussion

Encouraged by the ability of the naturally occurring damaged nucleobases 7,8-dihydro-8-oxoguanine (**oxoG**), 7,8-dihydro-8-oxoadenine (**oxoA**) and 1,*N*^6^-ethenoadenine (**εA**) to adopt a syn conformation, which enables Hoogsteen H-bonding with non-cognate native nucleobases in the opposite DNA strand, we have recently developed a 7,8-dihydro-8-oxo-1,*N*^6^-ethenoadenine (**oxo-εA**) modification that is a combination of **oxoA** and **εA** (Fig. 1C)^25^. Physicochemical methods and molecular modeling of DNA duplexes demonstrated that **oxo-εA** adopts the non-canonical syn conformation and pairs specifically with an adenine residue in the complementary strand without causing major distortions in local helical architecture. Considered as an adenine derivative, **oxo-εA** was approximately 99% mutagenic in living cells and caused predominantly A→T transversion mutations in *Escherichia coli*^25^. The modification was fluorescent but had relatively low brightness^26^. To improve its spectral properties, we preserved the H-bond pattern of **oxo-εA** and replaced the 1,*N*^6^-etheno bridge with a 3-methylbenzo[d]thiazolyl moiety. This moiety has a relatively high molar extinction coefficient for a heterobicyclic system (ε_222_ 37150 M^-1^cm^-1 27^) and is widely used as a component of asymmetric cyanine dyes that possess prospective photophysical properties^28–30^. At the same time, it might provide a rotary effect due to rotation around the C-N bond between the 7,8-dihydro-8-oxopurinyl and 3-methylbenzo[d]thiazolyl fragments (Fig. 1C). Thus, our initial hypothesis was that preserving the H-bonding pattern of **oxo-εA** would ensure selective pairing of **oxo-Ade** ^**BZT**^ with the adenine residue, while hindering the rotation around the C-N bond would increase in the fluorescence quantum yield and brightness.

The corresponding **oxo-Ade**^**BZT**^ nucleoside **4** was prepared in three steps, starting from 2’-deoxy-7,8-dihydro-8-oxoadenosine **1**^31^ (**Scheme 1**). Treatment of **1** with Ac_2_O in pyridine at 50°C gave a 5’-*O*-, 3’-*O*-, *N*^7^-triacetylated intermediate, which was then reacted with imidazole in CH_3_ OH at room temperature for selective *N*^7^ deblocking to afford **2**. The introduction of the 3-methylbenzo[d]thiazolyl moiety into the nucleobase scaffold was carried out based on an earlier observation that 2-aminopyrimidine derivatives react with a C2-activated N-alkylbenzo[d]thiazolium salt^32^. Briefly, the reaction between **2** and 3-methyl-2-(methylthio)benzo[d]thiazol-3-ium iodide in the presence of TEA at room temperature gave **3**, that was then deacetylated by treatment with aq. NH_3_ in THF at 40°C to afford the **oxo-Ade** ^**BZT**^ nucleoside **4**.

**Scheme 1.**
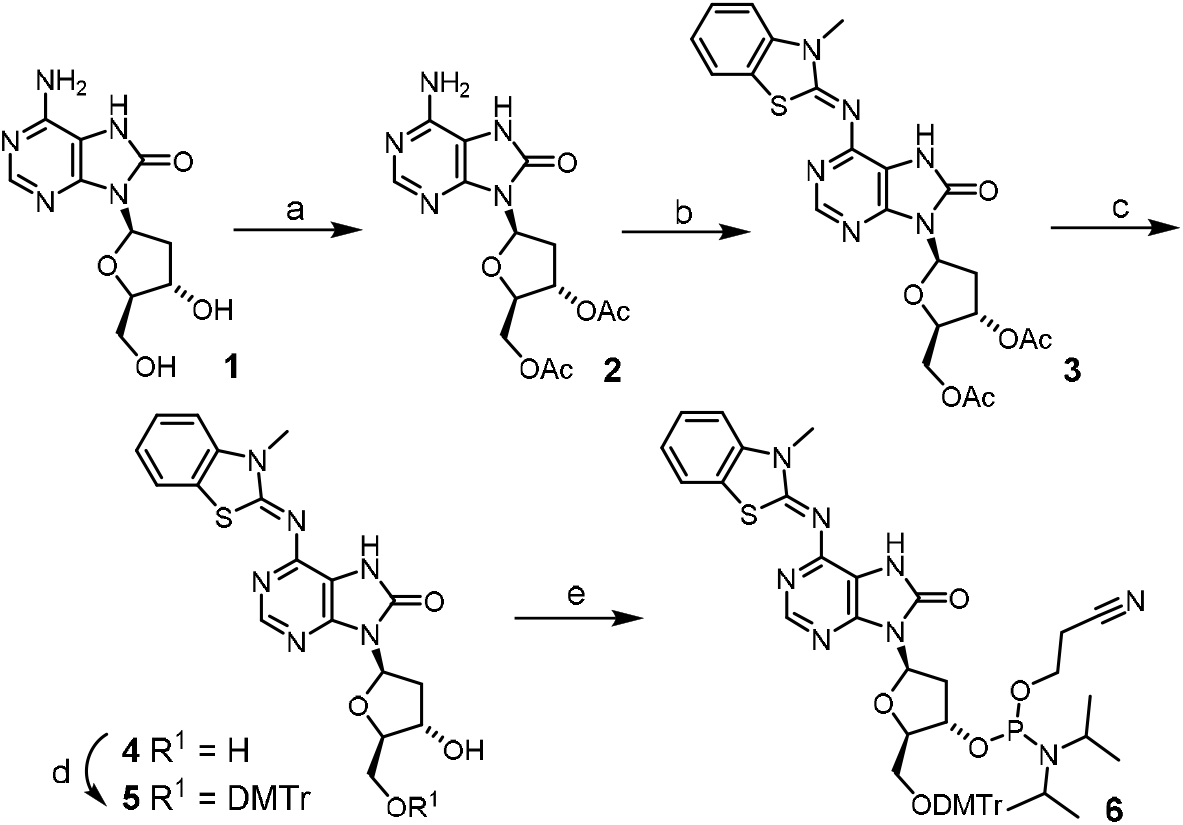
Synthesis of the 7,8-dihydro-8-oxo-6-(3-methylbenzo[d]thiazol-2(3H)-ylidene)adenine (**oxo-Ade**^**BZT**^) nucleoside phosphoramidite. Reagents and conditions: (a) Ac_2_O, pyridine, 50°C, then imidazole, CH_3_OH, rt, 81%; (b) 3-methyl-2-(methylthio)benzo[d]thiazol-3-ium iodide, TEA, CH_2_Cl_2_, 0°C→rt, 37%; (c) aq. NH_3_, THF, 40°C, 82%; (d) DMTr-Cl, pyridine, rt, 78%; (e) NCCH_2_CH_2_OP(Cl)N-*i*Pr_2_, DIPEA, CH_2_Cl_2_, 0°C→rt, 71%.

Photophysical measurements of the **oxo-Ade**^**BZT**^ nucleoside **4** revealed ε_361_ of 62000 ± 2000 M^-1^·cm^-1^ (absorption maximum at 361 nm) and ε_359_ of 47000 ± 1000 M^-1^·cm^-1^ (absorption maximum at 359 nm) in CH_3_OH and H_2_O, respectively. These values are exceptionally high for nucleoside derivatives, thus confirming the prospects of introducing the 3-methylbenzo[d]thiazolyl moiety into the nucleobase scaffold to significantly increase optical absorbance (Fig. S1). The maximum of the excitation spectrum was positioned at 346 nm, with a minor maximum at 322 nm, and the emission spectrum had one maximum at 420 nm, providing a Stokes shift of 74 nm in H_2_O (Fig. S2). The fluorescence quantum yield (*Φ*_f_) and brightness (*B*) of the **oxo-Ade**^**BZT**^ nucleoside were calculated to be 1.2 ± 0.2 % and 740 ± 10 M^-1^·cm^-1^, respectively, in H_2_O (Fig. S3). These values are lower than those of the reference compound quinine sulfate (52 % and 2960 M^-1^·cm^-1^ in 0.05 M H_2_SO_4_ ^33^) and commonly used dyes, such as the cyanine dye Cy5 (27 % and 21470 M^-1^·cm^-1^ in PBS^34^), and might originate from the molecular rotor effect. Free rotation around the C–N bond connecting the two heterobicyclic systems may indeed be responsible for non-emissive relaxation coupled to conformational change at the excited state^35^. To evaluate the potential of **oxo-Ade**^**BZT**^ as a fluorescent molecular rotor, fluorescence emission intensity and *Φ*_f_ were measured in various aqueous mixtures with an increasing content of glycerol and methanol. The latter was added as a control since it has a similar polarity but a much lower viscosity than glycerol. Fluorescence intensity and *Φ*_f_ increased dramatically with increasing viscosity (Figs. S4 and S5A), and the **oxo-Ade**^**BZT**^ nucleoside demonstrated a greater viscosity sensitivity (VS 0.61, Fig. S5B) then the widely used reference compound, 9-(2,2-dicyanovinyl)julolidine (DCVJ, VS 0.47)^36^, and other nucleoside-based molecular rotors, such as 8-DEA-tC (VS 0.11)^36^ and furan/thiophene-modified nucleosides (VS 0.26-0.40)^37^. Inspired by these results, we synthesized the **oxo-Ade**^**BZT**^ nucleoside phosphoramidite **6** for incorporation into ODNs. Derivative **4** was subjected to 5’-*O*-dimethoxytritylation and 3’-*O*-phosphitylation to afford the target phosphoramidite **6** with a yield of 13.6% over five steps.

To test the properties of **oxo-Ade**^**BZT**^ in single- and double-stranded contexts, four decameric modified ODNs were prepared using solid-phase DNA synthesis and named with three-letter codes NXN, where X is the **oxo-Ade**^**BZT**^ nucleotide, and N is the 5’- and 3’-flanking A, C, T, or G (Table S1). Since our initial hypothesis suggested that **oxo-Ade**^**BZT**^ would be a thymine analog, we compared the obtained thermodynamic parameters with those calculated for the duplexes formed by the corresponding unmodified ODNs, with X replaced by T (Table 1). For all the duplexes with any nucleobase opposite the **oxo-Ade**^**BZT**^ nucleotide, we observed well-defined S-shaped denaturation transitions (Fig. S6). The melting curves recorded at 260 nm fit well within the two-state model approximation^38^ and have entropic and enthalpic contributions to the hybridization energy changes (Tables S4 and S5). To our surprise, the introduction of **oxo-Ade**^**BZT**^ opposite adenine resulted in a sequence-dependent decrease in *T*_*m*_ (Δ*T*_*m*_ from - 4.0 to -14.9 °C) when adenine was opposite, and this decrease was particularly pronounced for **oxo-Ade**^**BZT**^ flanked by pyrimidines (Table 1). The presence of any nucleobase opposite the **oxo-Ade**^**BZT**^ nucleotide resulted in close *T*_*m*_ values, with variations usually not exceeding 10% of the mean for each NXN variant. This indicates that **oxo-Ade**^**BZT**^ acts as a universal nucleobase surrogate rather than a thymine mimic. These results are consistent with those obtained for other nucleobase analogs with an extended heteroaromatic system ^24,39^. In addition, a duplex with an AP site opposite **oxo-Ade**^**BZT**^ had much higher thermal stability (*T*_*m*_ 47.8°C) compared to the unmodified one (*T*_*m*_ 22.2 °C), indicating that stacking interactions rather than H-bonding make a major contribution to the stability (Fig. S6, Tables S4 and S5).

**Table 1.**
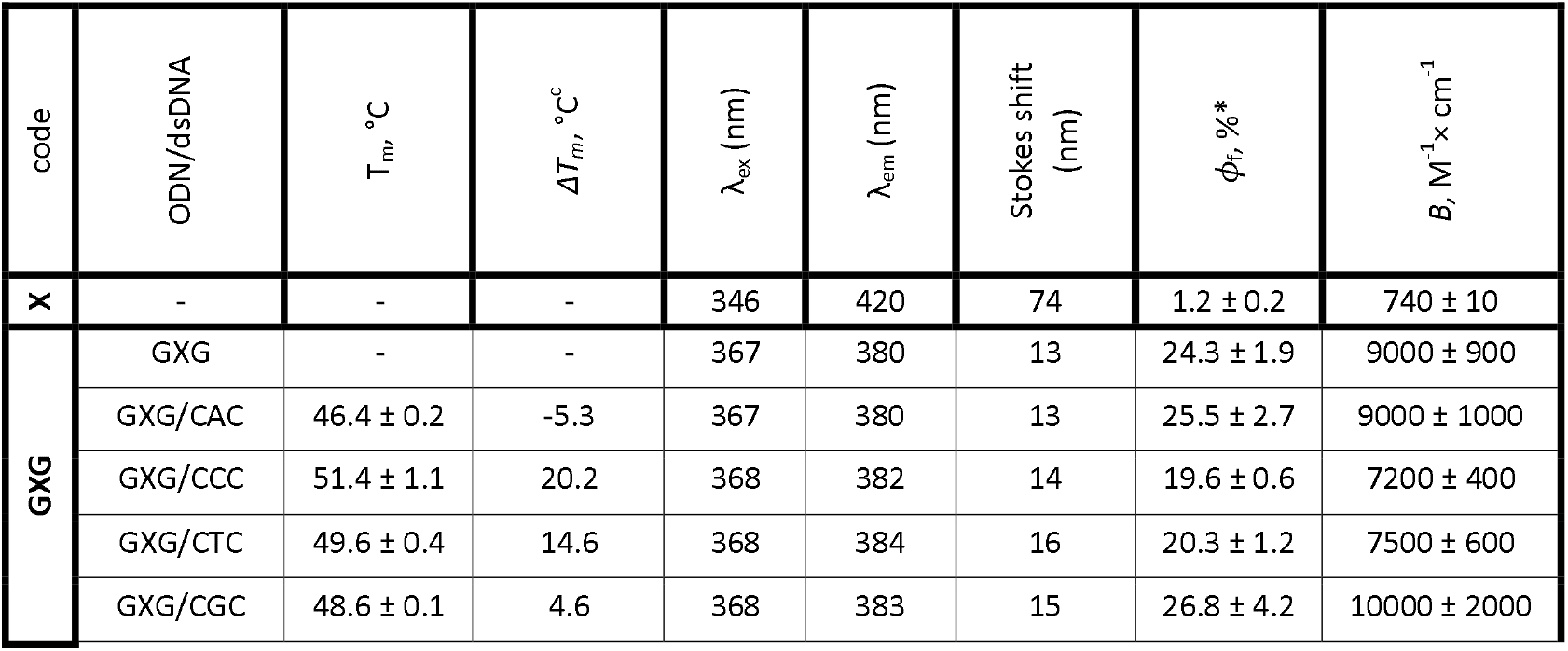
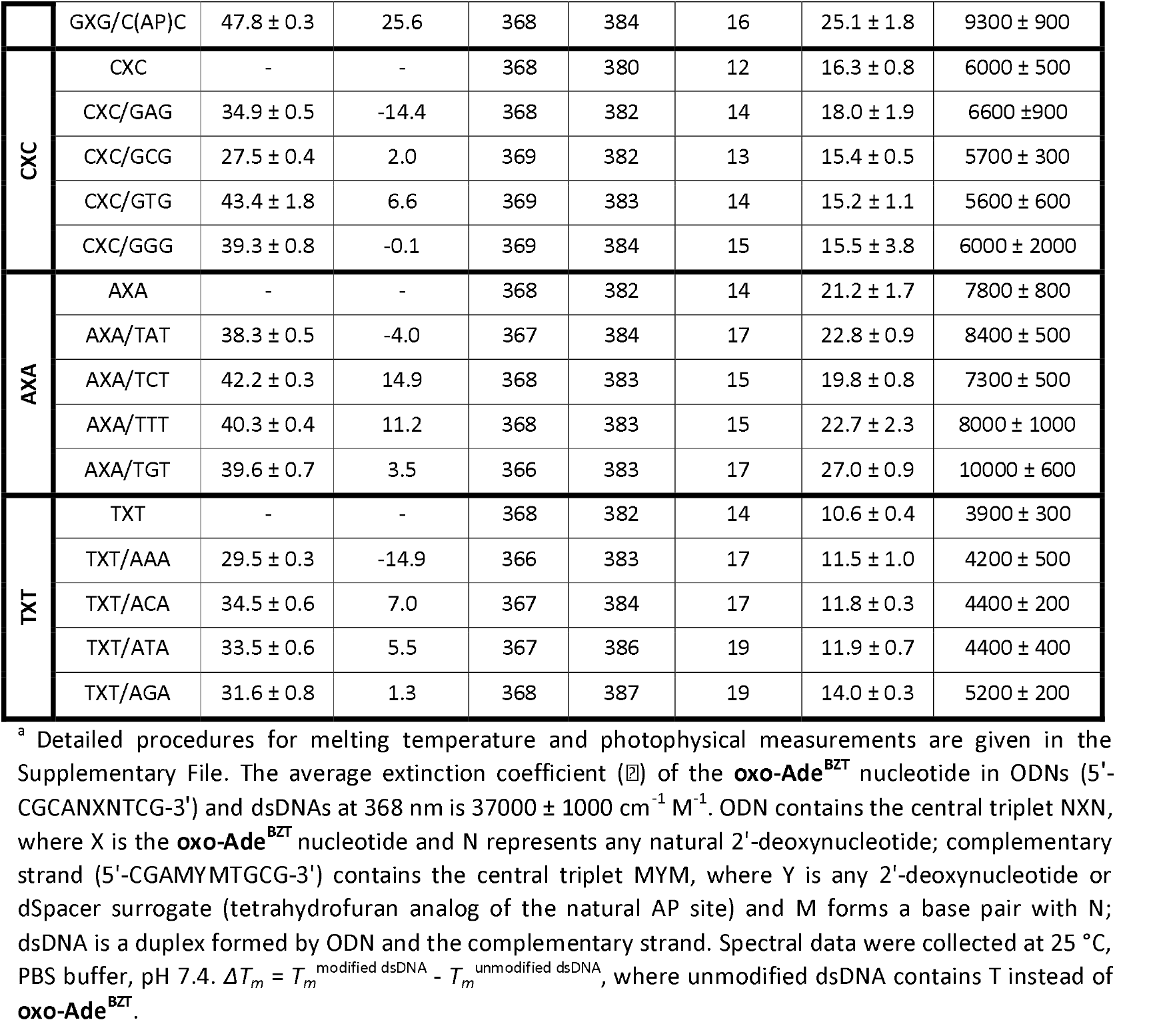
Melting temperatures (*T*_*m*_) and photophysical properties of **oxo-Ade**^**BZT**^-modified ODNs and their DNA duplexes^a^.

To study the influence of **oxo-Ade**^**BZT**^-flanking nucleobases on the thermal stability of duplexes more thoroughly, twelve additional modified ODNs were synthesized and named with three-letter codes NXM (Table S1), where X is the **oxo-Ade**^**BZT**^ nucleotide or T, and N and M are 5’- and 3’-flanking A, C, T or G . These ODNs were hybridized with complementary ODNs bearing A opposite **oxo-Ade**^**BZT**^. Overall, the thermal stability of the modified complexes decreased by Δ*T*_*m*_ from -0.8 to -28.0 °C compared to unmodified controls (Fig. S7, Tables S6 and S7). The destabilizing effect was stronger for 3’- and 5’-flanking pyrimidines than for purines, with the 3’-flank having a greater impact on thermal stability. Flanking purines generally caused less destabilization, and the smallest drop in *T*_*m*_ was observed with 3’-adenine. Overall, the results indicated a strong dependence of *T*_*m*_ on neighboring nucleobases and a major contribution from interactions between the 3’-flanking residue and **oxo-Ade**^**BZT**^.

After ensuring that modified ODNs are mainly in complexes with the complementary strands at 25°C, we evaluated the spectral properties of **oxo-Ade**^**BZT**^ in modified ODNs with the central triplet NXN, where N = A, C, T or G, in single- and double-stranded contexts (Table 1, Figs. S8 and S9). In all cases, the excitation maximum was red-shifted by 20-23 nm, and the emission maximum was blue-shifted by 33-40 nm. Thus, the Stokes shift decreased from 72 nm for the **oxo-Ade**^**BZT**^ nucleoside to 12-19 nm in ODNs and their complexes. At the same time, **oxo-Ade**^**BZT**^ incorporated into ODNs demonstrated a 10- to 27-fold increase in **Φ**_f_ compared to the free nucleoside, as well as an unprecedentedly high brightness of up to 10000 M^-1^ × cm^-1^ in a duplex context (Table 1). Quantum yield values were close in single-stranded and double-stranded contexts, depended mainly on the flanking nucleobases rather than the opposing ones (Fig. S9A), and correlated with **T**_*m*_ of the corresponding duplexes (Fig. S9B). The deviation from the regression line for the duplex formed by the ODN with a central triplet GXG can be explained by the quenching of the fluorescence by guanine residues^40^. To conclude this part, **oxo-Ade**^**BZT**^ is a bright fluorophore with sequence-dependent fluorescence.

To better understand the behavior of the **oxo-Ade**^**BZT**^ modification as a molecular rotor and universal nucleobase surrogate, structure optimization and molecular dynamics (MD) simulation were applied (for details, see the supplementary file). The structure of **oxo-Ade**^**BZT**^ suggests the possibility of rotation around three bonds. The first two bonds connect the 3-methylbenzo[d]thiazolyl and 7,8-dihydro-8-oxopurinyl moieties and can be described by dihedral angles d1 and d2, formed by the C5-N4=C6-N5 and C3-C5-N4=C6 atoms, respectively (Fig. 2A). The last (**N**-glycosidic) bond connects the heterocyclic system to the 2’-deoxyribose residue and can be described by the dihedral angle d3, formed by the C2’-C1’-N6-C1 atoms. Structure optimization using two isomers with a 180° rotation around d1 and two conformers with a 180° rotation around d2 as the starting point revealed isomer 1 with the sulfur atom located above the nitrogen atom with a lone electron pair as energetically most favorable (Fig. S10). In addition, a 180° rotation around the single bond represented by d2 (isomer 1 with minimal energy in Fig. S10), followed by energy minimization, was shown to be energetically unfavorable (Fig. S11). To establish the most energetically favorable conformation with a d1 value of 0° (as in isomer 1), MD simulation was carried out for four conformers with a 180° rotation around d2 and/or d3 of the **oxo-Ade** ^**BZT**^ nucleoside or the **oxo-Ade** ^**BZT**^ nucleotide within ODN with a central triplet GXG and within its duplex (Figs. S12-S34). For the duplex, the 2’-deoxyadenosine nucleotide in the anti or syn conformation was placed opposite **oxo-Ade** ^**BZT**^, in accordance with the experimental model. As a result, the prerequisites for the **oxo-Ade** ^**BZT**^ behavior as a universal nucleobase surrogate with a molecular rotor effect were established. Indeed, MD simulation analysis of the modified duplex shows that the **oxo-Ade** ^**BZT**^ nucleotide displaces the opposite adenine from the DNA double helix and participates in stacking interactions with flanking base pairs (Figs. S15 and S16). Thus, stacking interactions with flanking base pairs, rather than H-bonding with the opposite nucleobase, primarily determine thermal stability of the duplex. In addition, the results corroborate the observation that duplexes with flanking purines possess higher thermal stability, presumably due to their stronger stacking interactions with **oxo-Ade** ^**BZT**^ - constituting heterocycles. The standard deviation of the d1 and d2 angles representing the bonds between 3-methylbenzo[d]thiazolyl and 7,8-dihydro-8-oxopurinyl moieties decreased upon switching from the **oxo-Ade** ^**BZT**^ nucleoside to the ODN and the duplex (Fig. 2B, S23 and S28, Tables S8 and S9). This was in line with the increase in the observed fluorescence quantum yield (Table 1), further confirming that the **oxo-Ade** ^**BZT**^ modification acts as a molecular rotor.

**Figure 2.**
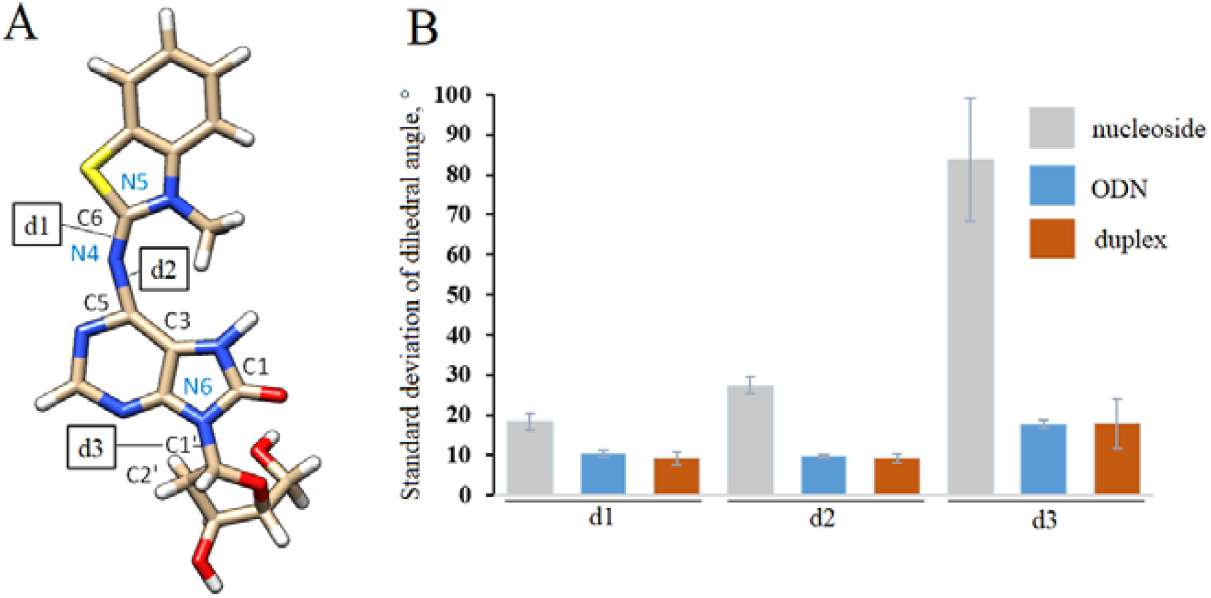
Structure of the **oxo-Ade**^**BZT**^ nucleoside with dihedral angles under study (A) and a histogram showing the flexibility of these dihedral angles in the **oxo-Ade**^**BZT**^ nucleoside, ODN with the central triplet G**oxo-Ade**^**BZT**^G and its duplex with 2’-deoxyadenosine nucleotide opposite the modification. Atom coloring: carbon - beige, oxygen - red, nitrogen-blue, sulfur - yellow and hydrogen - white.

As a result, the molecular rotor with a very high molar extinction coefficient (averaged **ε** 37000 M^-1^ × cm^-1^), a pronounced and stable light-up effect with exceptional brightness reaching up to **9950** M^-1^ × cm^-1^ depending on the flanking nucleobases, and the ability to maintain high opposite-nucleobase-insensitive thermal stability of the modified duplexes was developed. Since fluorescence quantum yield (and hence the brightness) of **oxo-Ade**^**BZT**^ decreases by an average of 20-fold when switching from ODNs and their duplexes to the nucleoside, the nucleobase surrogate can be used to monitor nucleic acid-protein interactions and the activity of enzymes involved in nucleic acid degradation^41,42^, DNA maintenance^15,43^ and RNA editing^44^. In addition, **oxo-Ade**^**BZT**^ can find application as a fluorescent reporter in biosensors based on a bulged-type enzyme-free recognition strategy, where interaction with the analyte triggers a switch from a conformationally mobile state within the bulge to a plane state with restricted librations ^45^. The universal behavior of **oxo-Ade**^**BZT**^ also makes it suitable for designing aptamers with improved characteristics^19,20,46^ and for structural studies^17^.

## Conclusion

In conclusion, based on our previous observation that simultaneous modification of the adenine residue with small substituents (namely, the 8-oxo group and the 1,**N**^6^-etheno bridge) shifted the equilibrium toward the syn conformation while creating an H-bond pattern that ensured exclusive pairing with the opposite adenine, we designed the **oxo-Ade** ^**BZT**^ modification. The introduction of a 3-methylbenzo[d]thiazolyl moiety at the **N**^6^ atom of the 7,8-dihydro-8-oxoadenine residue instead of the 1,**N**^6^-etheno bridge produced a molecular rotor with prospective photophysical properties. However, **oxo-Ade** ^**BZT**^ lost the ability to discriminate adenine from other natural nucleobases in the complementary strand, as has been observed for **oxo-εA**, highlighting the importance of fine-tuning the balance between H-bonding and stacking interactions. Instead, **oxo-Ade** ^**BZT**^ demonstrated a universal ability to maintain thermal stability of the modified duplexes, independent of the opposite nucleobase, with variation in melting temperatures generally not exceeding 10% for each pair of flanking nucleobases. According to MD simulation results, this property may be due to the extrusion of the opposite nucleobase from the DNA double helix and the involvement of **oxo-Ade** ^**BZT**^ in multiple stacking interactions with flanking base pairs. The latter limits the conformational mobility of **oxo-Ade** ^**BZT**^ within both ODN and its duplex and may explain an observed increase in fluorescence intensity.

## Supporting information

a summary of materials, instrumentations and general methods, synthesis, photophysical and thermodynamic measurements, and molecular modeling

## Author contributions

Conceptualization: A. A. L., A. V. A., investigation: A. A. P., P. N. K., E. S. B., V. A. B., data analysis and interpretation: A. A. P., A. A. L., A. V. A., visualization: A. A. P., A. A. L., A. V. A., methodology: A. A. L., A. V. A., supervision: A. A. L., A. V. A., writing – original draft: A. A. P., A. V. A., writing – review & editing: A. A. L., funding acquisition: A. A. L.

## Data availability

Supplementary information (SI): a summary of materials, instrumentations and general methods, synthesis, photophysical and thermodynamic measurements, and molecular modeling. See DOI:

## Acknowledgements

This work was supported by a Russian state-funded project for ICBFM SB RAS (grant number 123021600208-7). We thank Alexander Korshun and Anna Varizhuk for the fruitful discussion.

