## Supplementary material for "An Adenine-Based Molecular Rotor as a Universal Fluorescent Nucleobase with High Brightness": a summary of materials, instrumentations and general methods, synthesis, photophysical and thermodynamic measurements, and molecular modeling

### 1. Materials, instrumentations and general methods

All reagents were commercially available unless otherwise mentioned and used without further purification. All solvents were purchased from commercial sources. $^{1}H$ and $^{13}C$ NMR spectra of compounds **2**, **3**, and **4** were recorded on a Bruker Fourier 300 spectrometer (Bruker, Germany) at 300 and 75 MHz, respectively. $^{1}H$ and $^{13}C$ NMR spectra of compounds **5** and **6** were recorded on a Bruker Avance III 600 spectrometer (Bruker, Germany) at 600 and 150 MHz, respectively. $^{31}P$ NMR spectrum of compound **6** was recorded on a Bruker Avance III 600 spectrometer (Bruker, Germany) at 243 MHz. Chemical shifts are reported in δ (ppm) units using residual $^{1}H$ signals from deuterated solvents as references. The coupling constant (J) is given in Hz. Column chromatography (CC) was performed on silica gel (0.040–0.063 mm, Merck, Germany). Thin layer chromatography (TLC) was performed on plates (Merck, Germany) precoated with silica gel (60 µm, F254) and visualized using UV light (254 and 365 nm). ESI HR mass spectra were acquired on a LTQ FT Ultra (Thermo Electron Corp., Bremen, Germany) mass spectrometer in a positive ion mode. 2'-Deoxy-7,8-dihydro-8-oxoadenosine **1** was prepared according to the reported procedure[1].

Absorption spectra were recorded in 200 µL quartz cuvettes with optical path length of 0.2 cm on a Cary 3500 spectrophotometer (Agilent Technology, USA) in the wavelength range of 200 - 800 nm at 25 °C. The experiment was repeated at least twice.

UV-melting analysis was carried out in 200 µL quartz cuvettes with an optical path length of 0.2 cm on a Cary 300-Bio spectrophotometer (Varian, Australia). Registration of optical density was performed at 260 and 270 nm using a slit width of 2 nm and signal averaging time of 0.5 s in the temperature range of 5 - 95 °C with heating/cooling rate of 0.5 °C/min[2]. Experiments were performed at an oligodeoxyribonucleotide (ODN) concentration of $8 \times 10^{-6}$ M in phosphate-buffered saline (PBS, pH 7.4).

The molar extinction coefficients of unmodified ODNs (umODNs) were calculated from molar extinction coefficients for mono- and dinucleotides[3]. The molar extinction coefficients of modified ODNs (mODNs) were calculated using the following equation:

$$\varepsilon_{260}(mODN) = \varepsilon_{260}(umODN) + \left(\varepsilon_{260}(oxo - Ade^{BZT}) - \varepsilon_{260}(dA)\right) \quad (1).$$

Fluorescence spectra were recorded in 100 µL quartz cuvettes with an optical path length of 1.0 cm on a Cary Eclipse Fluorescence Spectrometer (Agilent Technology, USA). The experiment was repeated at least twice. The molar extinction coefficient ($\varepsilon$) of the **oxo-Ade$^{BZT}$** nucleoside was determined based on three concentration points.

To determine a relative fluorescence quantum yield ($\Phi_f$) of the **oxo-Ade$^{BZT}$** nucleoside and the corresponding **oxo-Ade$^{BZT}$** nucleotide within mODNs and their complexes, the reported method with quinine bisulfate in 0.05 M $H_2SO_4$ as a fluorescence reference standard with known quantum yield was used[4,5]. $\Phi_f$ was calculated using the following equation:

$$\phi_{f,x} = \phi_{f,st} \frac{F_X}{F_{st}} \times \frac{f_{st}}{f_x} \times \frac{n_x^2(\lambda em)}{n_{st}^2(\lambda em)} \quad (2),$$

where $F$ is the integral photon flux, $f$ is the optical absorption coefficient, $n$ is the refractive index of the solvent used. The index $X$ denotes the sample, and the index $st$ denotes the standard. The optical absorption coefficient $f$ was calculated using the following equation:

$$f = 1 - 10^{-D(\lambda_{ex})} \quad (3),$$

where $D(\lambda_{ex})$ is the optical density at $\lambda_{ex}$.

Brightness ($B$) of the fluorophore was determined using the following equation:

$$B(X) = \varepsilon_{\lambda(max)}(X) \times \phi(X) \quad (4),$$

where $\varepsilon_{\lambda(max)}(X)$ is the molar absorption coefficient at the excitation wavelength.

To determine $\Phi_f$ of the **oxo-Ade^BZT** nucleotide within mODNs and their complexes, their solutions at concentrations of $1.5\times10^{-6}$ and $2\times10^{-7}$ M, respectively, in 0.01 M PBS buffer (pH 7.4) were used.

The values of thermodynamic parameters ($\Delta H°, \Delta S°, \Delta G°_{37}$) of the complex formation were determined by fitting of heating and cooling curves obtained at wavelengths of 260 and 270 nm using two-state model approximation[6]. The values obtained were averaged. The melting temperature ($T_m$ in °C) of the DNA duplexes was calculated using the following equation:

$$T_m = \frac{\Delta H°}{\Delta S° + R \times ln\left(\frac{C_t}{4}\right)} - 273.15 \quad (5),$$

where $C_t = [A]_0 + [B]_0$ is a total ODN concentration ($[A]_0 = [B]_0$).

The Gibbs energy change was calculated using the following equation at 37 °C (310.15 K):

$$\Delta G°_{37} = \Delta H° - T\Delta S° \quad (6).$$

### 2. Synthesis and characterization of novel compounds

5',3'-*O*-acetyl-2'-deoxy-7,8-dihydro-8-oxoadenosine **2**

To a solution of 2'-deoxy-7,8-dihydro-8-oxoadenosine **1** (1.39 g, 5.20 mmol) in dry pyridine (25 mL) acetic anhydride (1.5 mL, 15.9 mmol) was added at room temperature. The solution was kept at 50°C overnight and then poured into 5% aqueous $NaHCO_3$ solution (100 mL). The triacetylated intermediate was extracted with $CH_2Cl_2$ (2 × 50 mL), concentrated *in vacuo* followed by co-evaporation with toluene (2 × 20 mL). The residue was dissolved in $CH_3OH$ (50 mL) and to the resulting solution imidazole (0.49 g, 7.2 mmol) was added. After 5 hours at room temperature, the reaction mixture was concentrated, and the residue was partitioned between a mixture of *n*-$C_4H_9OH/CH_2Cl_2$ (100 mL, 1:9, v/v) and $H_2O$ (100 mL). The organic layer was separated, concentrated *in vacuo* and co-evaporated with toluene (2 × 20 mL). The crude product was purified by column chromatography on silica gel (0-4% $CH_3OH$ in $CH_2Cl_2$) affording **2** (1.47 g, 4.18 mmol, 81%) as white foam. $^1$H NMR (300 MHz, DMSO-*d₆*): δ 10.34 (s, 1H), 8.04 (s, 1H), 6.51 (s, 2H), 6.15 (t, *J* = 7.1 Hz, 1H), 5.47-5.39 (m, 1H), 4.40-4.29 (m, 1H), 4.17-4.07 (m, 2H), 3.43-3.28 (m, 1H), 2.36-2.25 (m, 1H), 2.07 (s, 3H), 1.98 (s, 3H). $^{13}$C NMR (75 MHz, DMSO-$d_6$): δ 170.0, 169.9, 151.1, 150.7, 147.0, 146.4, 103.3, 80.8 (2C), 74.3, 63.5, 32.4, 20.7, 20.4. HRMS (ESI) m/z: calcd for $C_{14}H_{18}N_5O_6^+$ $[M+H]^+$: 352.1252; found 352.1257.

5',3'-*O*-acetyl-2'-deoxy-7,8-dihydro-8-oxo-6-(3-methylbenzothiazol-2(3H)-ylidene)adenosine **3**

To a stirred solution of **2** (1.41 g, 4.01 mmol) in $CH_2Cl_2$ (40 mL) 3-methyl-2-(methylthio)benzo[d]thiazol-3-ium iodide (1.62 g, 5.01 mmol) and triethylamine (700 µL, 5.00 mmol) were added at 0°C. The reaction mixture was stirred at room temperature overnight, washed with $H_2O$ (2 × 20 mL) and concentrated *in vacuo*. The crude product was purified by column chromatography on silica gel (0-1% $CH_3OH$ in $CH_2Cl_2$) affording **3** (0.74 g, 1.48 mmol, 37%) as an off-white amorphous solid. $^1$H NMR (300 MHz, DMSO-$d_6$): δ 11.63 (s, 1H), 8.46 (s, 1H), 7.79 (d, J = 7.7 Hz, 1H), 7.49-7.40 (m, 2H), 7.29-7.20 (m, 1H), 6.24 (t, *J* = 7.1 Hz, 1H), 5.53-5.44 (m, 1H), 4.43-4.33 (m, 1H), 4.21-4.12 (m, 2H), 3.87 (s, 3H), 3.45-3.32 (m, 1H), 2.43-2.32 (m, 1H), 2.09 (s, 3H), 1.99 (s, 3H). $^{13}$C NMR (75 MHz, DMSO-$d_6$): δ 170.0, 170.0, 159.1, 152.0, 148.7, 148.0, 144.9, 137.7, 126.7, 125.3, 122.9, 122.2, 112.4, 111.2, 80.9, 80.9, 74.2, 63.5, 32.5, 31.2, 20.7, 20.4. HRMS (ESI) m/z: calcd for $C_{22}H_{23}N_6O_6S^+$ $[M+H]^+$: 499.1394; found 499.1394.

2'-deoxy-7,8-dihydro-8-oxo-6-(3-methylbenzothiazol-2(3H)-ylidene)adenosine **4**

To a solution of **3** (0.70 g, 1.41 mmol) in THF (15 mL) aqueous $NH_3$ solution (33%, 3 mL) was added and the resulting mixture was kept at 40°C overnight. After concentration, the product was triturated with $CH_3OH$ and filtered affording **4** (0.48 g, 1.16 mmol, 82%) as a white amorphous solid. $^1$H NMR (300 MHz, DMSO-$d_6$): δ 11.61 (s, 1H), 8.44 (s, 1H), 7.79 (d, J = 7.7 Hz, 1H), 7.49-7.40 (m, 2H), 7.29-7.20 (m, 1H), 6.22 (t, *J* = 7.3 Hz, 1H), 5.24 (d, J = 4.2 Hz, 1H), 5.00 (dd, J = 7.4 Hz, J = 4.6 Hz, 1H), 4.48-4.39 (m, 1H), 3.87 (s, 3H), 3.87-3.79 (m, 1H), 3.70-3.60 (m, 1H), 3.54-3.44 (m, 1H), 3.13-3.01 (m, 1H), 2.12-2.01 (m, 1H). $^{13}$C NMR (75 MHz, DMSO-$d_6$): δ 159.1, 152.1, 148.5, 148.0, 144.9, 137.7, 126.7, 125.3, 122.9, 122.2, 112.5,

111.2, 87.4, 81.3, 71.2, 62.2, 35.9, 31.2. HRMS (ESI) m/z: calcd for $C_{18}H_{19}N_6O_4S^+$: $[M+H]^+$: 415.1183; found 415.1187.

5'-*O*-(4,4'-Dimethoxytrityl)-2'-deoxy-7,8-dihydro-8-oxo-6-(3-methylbenzothiazol-2(3H)-ylidene)adenosine **5**

Compound **4** (0.46 g, 1.11 mmol) was co-evaporated with anhydrous pyridine (15 mL), dissolved in anhydrous pyridine (20 mL), and to the resulting solution 4,4'-dimethoxytrityl chloride (0.48 g, 1.42 mmol) was added. After 4 h at room temperature, the reaction mixture was diluted with $CH_2Cl_2$ (50 mL) and washed with 5% aqueous $NaHCO_3$ solution (50 mL) and saline (50 mL). The organic layer was concentrated *in vacuo* and co-evaporated with toluene (2 × 30 mL). Purification was performed by column chromatography on silica gel (0–2% $CH_3OH$ in $CH_2Cl_2$ with 0.1% triethylamine), yielding **5** as yellowish foam (0.62 g, 0.87 mmol, 78%). $^1H$ NMR (600 MHz, DMSO-$d_6$): δ 11.44 (s, 1H), 8.29 (s, 1H), 7.78 (d, J = 7.6 Hz, 1H), 7.47-7.43 (m, 2H), 7.37-7.34 (m, 2H), 7.27-7.19 (m, 7H), 7.18-7.14 (m, 1H), 6.82-6.78 (m, 2H), 6.77-6.74 (m, 2H), 6.24 (dd, *J* = 7.2 Hz, J = 6.3 Hz, 1H), 5.15 (d, J = 4.8 Hz, 1H), 4.59-4.54 (m, 1H), 3.97-3.92 (m, 1H), 3.87 (s, 3H), 3.71 (s, 3H), 3.68 (s, 3H), 3.23-3.11 (m, 3H), 2.18-2.12 (m, 1H). $^{13}C$ NMR (150 MHz, DMSO-$d_6$): δ 158.8, 157.7, 157.6, 152.0, 148.2, 148.1, 144.8, 144.6, 137.6, 135.6, 135.5, 129.4 (2C), 129.2 (2C), 127.5 (2C), 127.2 (2C), 126.5, 126.1, 125.1, 122.6, 122.0, 112.7 (2C), 112.6 (2C), 112.3, 110.9, 85.1, 85.0, 80.6, 70.8, 64.0, 54.7, 54.6, 35.5, 31.0. HRMS (ESI) m/z: calcd for $C_{39}H_{37}N_6O_6S^+$: $[M+H]^+$: 717.2490; found 717.2492.

5'-*O*-(4,4'-Dimethoxytrityl)-3'-*O*-(*N*,*N*-diisopropylamino-2-cyanoethoxyphosphinyl)-2'-deoxy-7,8-dihydro-8-oxo-6-(3-methylbenzothiazol-2(3H)-ylidene)adenosine **6**

Compound **5** (0.60 g, 0.83 mmol) was co-evaporated with anhydrous $CH_2Cl_2$ (2 × 10 mL), dissolved in anhydrous $CH_2Cl_2$ (15 mL) and then DIPEA (0.63 mL, 3.40 mmol) was added. To the resulting solution, 2-cyanoethyl diisopropylchlorophosphoramidite (0.40 g, 1.66 mmol) was added at 0 °C under argon atmosphere. After 1 hour at room temperature, the reaction was quenched by the addition of $CH_3OH$ (1.0 mL), the organics were diluted with $CH_2Cl_2$ (40 mL), washed with 5% aqueous $NaHCO_3$ solution (40 mL) and brine (40 mL). The organic layer was dried over $Na_2SO_4$ and concentrated *in vacuo*. The crude product was purified by column chromatography on silica gel (0→5% acetone in $CH_2Cl_2$ with 1.0 % triethylamine, yielding **6** as yellowish foam (0.54 g; 0.59 mmol; 71%). $^1H$ NMR (600 MHz, $CD_2Cl_2$): δ 8.37 & 8.36 (s, 1H, diastereomers), 7.62 (d, J = 7.7 Hz, 1H), 7.46-7.40 (m, 4H), 7.32-7.14 (m, 8H), 6.77-6.71 (m, 4H), 6.37 & 6.36 (t, *J* = 7.1 Hz, 1H, diastereomers), 4.98-4.92 & 4.90-4.85 (m, 1H, diastereomers), 3.88-3.77 (m, 1H), 3.82 (s, 3H), 3.73 & 3.73 (s, 3H, diastereomers), 3.72 & 3.72 (s, 3H, diastereomers), 3.67-3.56 (m, 2H), 3.44-3.31 (m, 2H), 2.76-2.71 (m, 3H), 2.64-2.61 (m, 1H), 2.51-2.48 (m 1H), 2.38-2.32 (m, 1H, diastereomers), 1.22-1.18 (m, 12H).$^{31}P$ NMR (243 MHz, $CD_2Cl_2$): δ 148.3, 148.3. HRMS (ESI) m/z: calcd for $C_{48}H_{54}N_8O_7PS^+$: $[M+H]^+$: 917.3568; found 917.3565.

### 3. Oligonucleotide synthesis

ODNs were obtained using ASM-2000 DNA automatic synthesizer (Biosset, Novosibirsk, Russia) by the phosphoramidite method according to the manufacturer's protocols. Protected 2'-deoxyribonucleoside 3'-phosphoramidites were purchased from Hongene Biotech (Union City, CA, USA). Unylinker-CPG (500 Å) and S-ethylthio-*1H*-tetrazole were purchased from ChemGenes (Wilmington, MA, USA). Condensation time for the phosphoramidite **6** was 4 minutes. ODNs on CPG were deprotected using AMA – 1:1 (v/v) conc. aq. ammonia and 40% aq. methylamine for 30 min at 65°C. Purification of 5'-*O*-DMTr-protected ODNs was carried on Glen-Pak (Glen Research, Sterling, VA, USA) reverse-phased cartridges according to standard protocol. HPLC/MS spectra of ODNs were registered using Agilent 1100 (Agilent Technologies, USA) with quadrupole MSD G1956B (ESI) in negative mode (buffer A – 100 mM (hexafluoroisopropanol with diisopropylamine), pH 8.9, buffer B – acetonitrile.

Table S1. Synthesized modified ODNs with calculated and observed masses.

| Sequence name | Sequence 5'→3' | Expected Molecular Weight (g/mol) | Mass Spec Analysis (amu) |
|---|---|---|---|
| TXC | CGCAT**X**CTCG | 3151 | 3150 |

| | | | |
|---|---|---|---|
| TXA | CGCAT**X**ATCG | 3175 | 3174 |
| TXG | CGCAT**X**GTCG | 3191 | 3190 |
| TXT | CGCAT**X**TTCG | 3166 | 3166 |
| AXC | CGCAA**X**CTCG | 3160 | 3158 |
| AXA | CGCAA**X**ATCG | 3184 | 3184 |
| AXG | CGCAA**X**GTCG | 3200 | 3198 |
| AXT | CGCAA**X**TTCG | 3175 | 3174 |
| CXC | CGCAC**X**CTCG | 3136 | 3136 |
| CXA | CGCAC**X**ATCG | 3160 | 3158 |
| CXG | CGCAC**X**GTCG | 3176 | 3176 |
| CXT | CGCAC**X**TTCG | 3151 | 3148 |
| GXC | CGCAG**X**CTCG | 3176 | 3176 |
| GXA | CGCAG**X**ATCG | 3200 | 3200 |
| GXG | CGCAG**X**GTCG | 3216 | 3214 |
| GXT | CGCAG**X**TTCG | 3191 | 3188 |

**X** = **oxo-Ade$^{BZT}$** nucleotide

### 4. Determination of the molar extinction coefficient (*ε*) of oxo-Ade$^{BZT}$ nucleoside

The molar extinction coefficient of the **oxo-Ade$^{BZT}$** nucleoside was calculated using the Beer–Lambert law:

$$\varepsilon_\lambda(X) = \frac{D_\lambda}{C(X)\cdot 0.2} \quad (7),$$

where $\varepsilon_\lambda$ - molar extinction coefficient at specified wavelength, $D_\lambda$ – absorbance at specified wavelength and C – concentration of the **oxo-Ade$^{BZT}$** nucleoside (X).

The UV spectra of the **oxo-Ade$^{BZT}$** nucleoside at a concentration of 2.5 µM were recorded in $CH_3OH$ and $H_2O$ (Figure S1A). The absorption maxima were at 361 and 359 nm in $CH_3OH$ and $H_2O$, respectively, with additional maxima at 347 and 222 nm in $CH_3OH$.

To determine the molar extinction coefficient of the **oxo-Ade$^{BZT}$** nucleoside, the stock solution at a concentration of 0.2 mM in $CH_3OH$ was prepared followed by the registration of UV-Vis absorbance spectra for 13.8x, 26.6x and 55.1x dilutions in triplicate (Figure S1B, Table S2). The calculated $\varepsilon_{260}$ and $\varepsilon_{361}$ were 8300±500 and 62000±2000 $M^{-1}cm^{-1}$, respectively, in $CH_3OH$. By the comparison of UV spectra in $CH_3OH$ and $H_2O$ (Figure S1A), $\varepsilon_{260}$ and $\varepsilon_{359}$ were calculated to be equal to 10000±580 and 47000±1000 $M^{-1}cm^{-1}$, respectively, for the **oxo-Ade$^{BZT}$** nucleoside in $H_2O$.

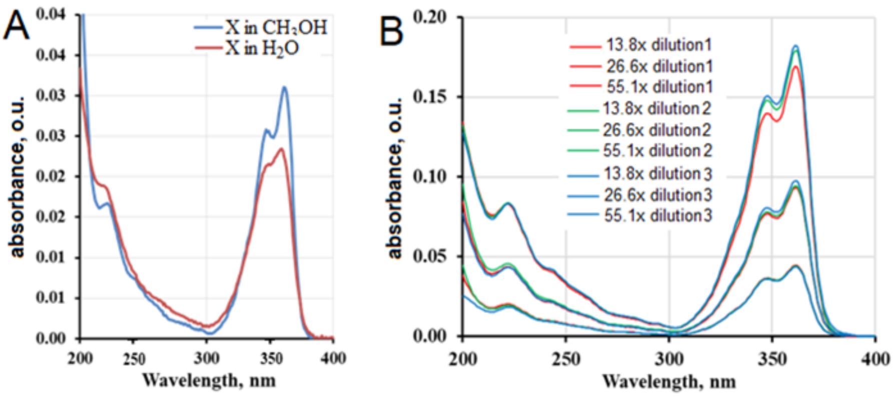

Figure S1. Absorption spectrum of the **oxo-Ade$^{BZT}$** nucleoside in $CH_3OH$ and $H_2O$ (A) and in $CH_3OH$ at different dilutions (B).

Table S2. Absorbance and molar extinction coefficient (*ε*) of the **oxo-Ade$^{BZT}$** nucleoside at different concentrations in $CH_3OH$.

| sample | Dilution | $D_{260}$, o.u. | $D_{361}$, o.u. | $\varepsilon_{260}$ | $\varepsilon_{361}$ |
|---|---|---|---|---|---|
| 1 | 55.1 | 0.006 | 0.044 | 7887 | 61192 |
| | 26.6 | 0.013 | 0.093 | 8374 | 61948 |
| | 13.8 | 0.024 | 0.169 | 8321 | 58347 |
| 2 | 55.1 | 0.006 | 0.044 | 7744 | 60899 |
| | 26.6 | 0.013 | 0.094 | 8878 | 62795 |
| | 13.81 | 0.025 | 0.179 | 8779 | 61800 |
| 3 | 55.1 | 0.005 | 0.044 | 7541 | 60059 |
| | 26.6 | 0.013 | 0.098 | 8572 | 64919 |
| | 13.8 | 0.025 | 0.182 | 8697 | 62916 |

### 5. Determination of relative fluorescence quantum yield of the oxo-Ade$^{BZT}$ nucleoside

To determine the fluorescence quantum yield ($\Phi_f$), quinine sulfate with known quantum yield of 52% in 0.05 M $H_2SO_4$ was used as a standard[4]. The solutions of the **oxo-Ade$^{BZT}$** nucleoside in deionized $H_2O$ (MQ) with concentrations ranging from $2\times10^{-5}$ to $2.2\times10^{-7}$ M were used. The fluorescent spectrum of the **oxo-Ade$^{BZT}$** nucleoside has one maximum at 420 with excitation at 346 nm (Figure S2).

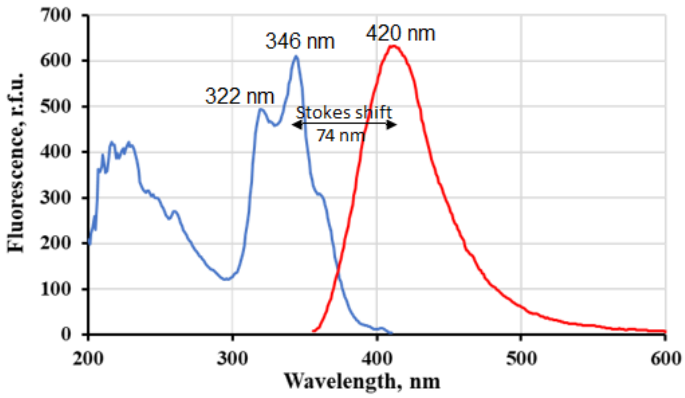

Figure S2. Excitation and emission spectra of the **oxo-Ade$^{BZT}$** nucleoside in $H_2O$ at a concentration of $7.3\times10^{-6}$ M.

The intersection wavelength of absorption spectra of the **oxo-Ade$^{BZT}$** nucleoside and quinine sulfate was determined to be 350 nm (Figure S3A). Then, the optical density of the **oxo-Ade$^{BZT}$** nucleoside and quinine sulfate at 350 nm and the emission spectra were recorded (Figure S3B) and the data are summarized in Table S3. The quantum yield for each concentration was determined, and the data were averaged. Using the equations (2) and (3), and the data from Table S3, $\Phi_f$ of the **oxo-Ade$^{BZT}$** nucleoside in $H_2O$ was calculated to be 1.2 ± 0.2 %.

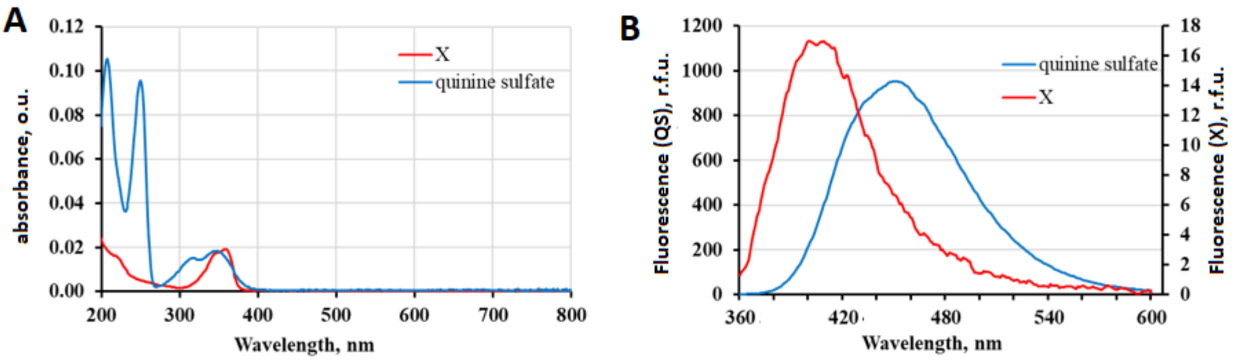

Figure S3. UV-Vis absorption spectra of the **oxo-Ade$^{BZT}$** nucleoside (X) in $H_2O$ at a concentration of $1.6\times10^{-5}$ M and quinine sulfate in 0.05 M $H_2SO_4$ at a concentration of $2.5\times10^{-6}$ M (A). Emission spectra of the **oxo-**

**Ade**$^{BZT}$ nucleoside (X) at a concentration of 2.0×10$^{-5}$M and quinine sulfate at a concentration of 5.7×10$^{-4}$ M at an excitation wavelength of 350 nm.

Table S3. Dependence of the area under the emission spectrum and optical density on the concentration of quinine sulfate and the **oxo-Ade**$^{BZT}$ nucleoside (X) with excitation at 350 nm

| Quinine sulfate (standard) | | | **oxo-Ade**$^{BZT}$ nucleoside (X) | | |
|---|---|---|---|---|---|
| C, M | $Dst(350nm)$ | $F_{st}$ | C, M | $Dx(350nm)$ | $F_X$ |
| 5.7×10$^{-4}$ | 0.641 | 81739 | 2.0×10$^{-5}$ | 0.353 | 1343 |
| 1.1×10$^{-4}$ | 0.124 | 38151 | 3.8×10$^{-5}$ | 0.141 | 1180 |
| 3.3×10$^{-5}$ | 0.038 | 11851 | 1.1×10$^{-6}$ | 0.034 | 195 |
| 1.5×10$^{-5}$ | 0.017 | 5307 | 5.7×10$^{-7}$ | 0.009 | 68 |

Using the equation (4), the fluorophore brightness (*B*) at the wavelength of the absorption maximum (361 nm) in $H_2O$ was calculated to be 740 ± 11 M$^{-1}$ × cm$^{-1}$.

### 6. Effect of viscosity on fluorescence quantum yield

Molecular rotors are known to possess viscosity-dependent fluorescent properties. The responsiveness of the **oxo-Ade**$^{BZT}$ nucleoside to viscosity changes using mixtures of $H_2O$ and glycerol was determined. Mixtures of $H_2O$ and $CH_3OH$ were used as controls, since the latter has very similar polarity but drastically different (lower) viscosity compared to glycerol.

The fluorescence spectra of the **oxo-Ade**$^{BZT}$ nucleoside were examined in aqueous mixtures with an alcohol content ranging from 0 to 93% at 25 °C (Figure S4). The increase in an alcohol content led to the increase in fluorescence intensity and changes in the fluorescence spectra shape, with the Stokes shift value decreased from 74 nm in $H_2O$ to 29-31 nm at 93% alcohol content.

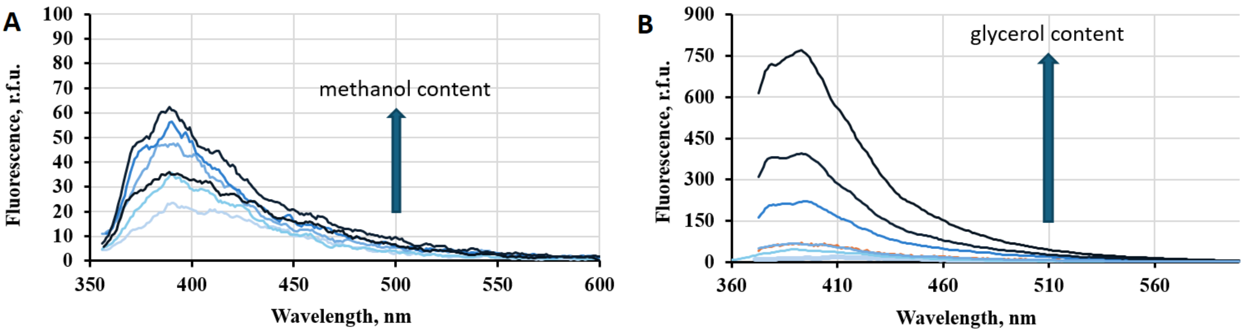

Figure S4. Emission of the **oxo-Ade**$^{BZT}$ nucleoside as a function of viscosity in different mixtures of $H_2O$-$CH_3OH$ (A) and $H_2O$-glycerol (B). Excitation at 350 nm, 10 nm excitation slit widths, 0.5 nm emission slit widths.

Then, $\Phi_f$ at an excitation wavelength of 350 nm was calculated and plotted as a function of alcohol content (Figure S5A). A slight increase in $\Phi_f$ from 1.2 ± 0.2% in $H_2O$ to 5.8 ± 0.4% at 50-75% $CH_3OH$ content was observed. In contrast, an increase in the glycerol content and, therefore, viscosity, caused a dramatic increase in $\Phi_f$ up to 78.1 ± 5.5% at 93% glycerol content.

The relationship between the fluorescence intensity of the molecular rotor and the viscosity can be determined by the following Förster–Hoffmann equation:

$$logI = C + xlog(\eta) \quad (8),$$

where $\eta$ represents the viscosity, *I* represents the integrated fluorescence intensity of the molecular rotor, *C* is a constant, and *x* represents the sensitivity of the molecular rotor towards viscosity.

According to equation (8), the **oxo-Ade**$^{BZT}$ nucleoside has a slope representing viscosity sensitivity equal to 0.61 (Figure S5B).

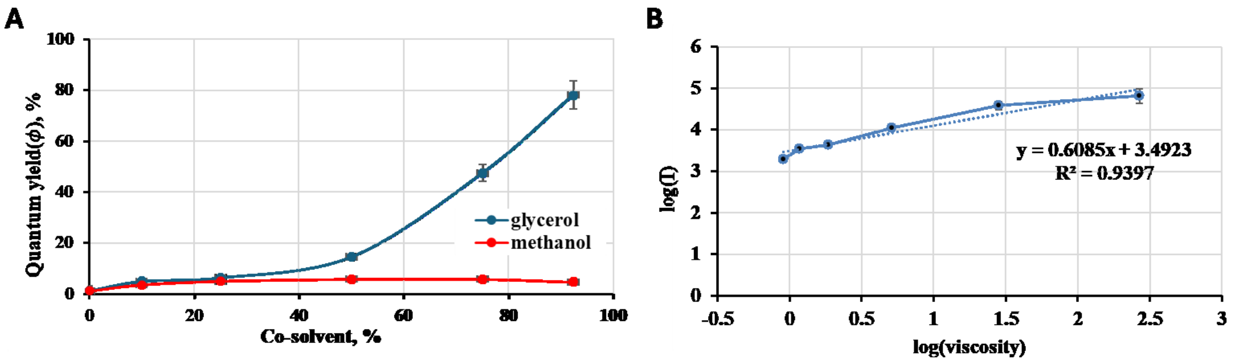

Figure S5. Dependence of fluorescence quantum yield on the alcohol content (A) and the integrated fluorescence intensity on the viscosity of glycerol according to the Foster-Hoffman relation (B).

### 7. Melting Temperature Analysis ($T_m$)

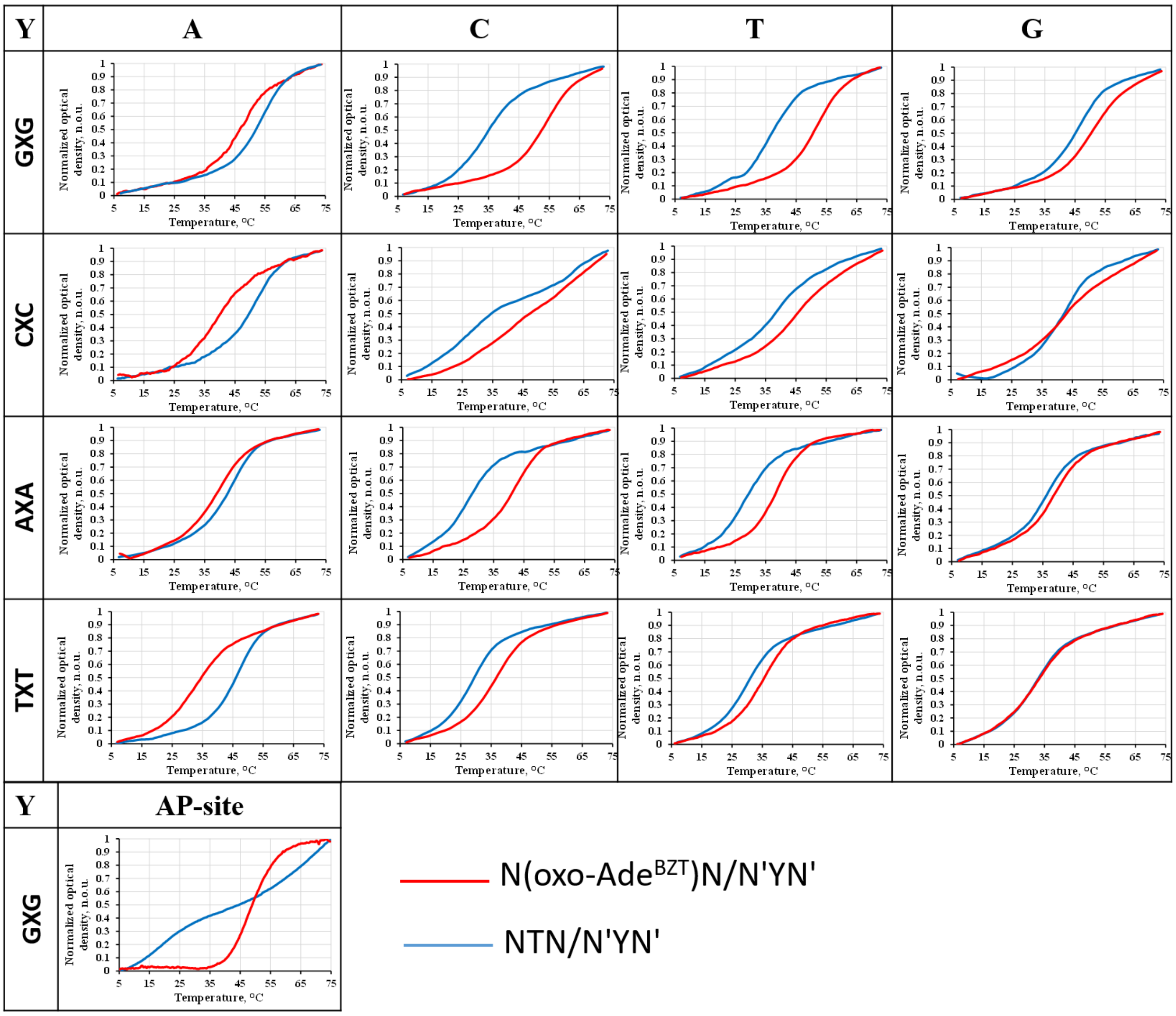

Figure S6. UV melting curves for native and modified duplexes with the central triplet NXN/N'YN' (X = **oxo-Ade**$^{BZT}$ or T; Y = A, C, T, G or a tetrahydrofuran analog of the natural AP site; N and N' = A, C, T or G; N and N’ form Watson-Crick base pair) registered at 260 nm.

Table S4. Thermodynamic parameters of DNA duplexes with the central triplet NTN/N'YN', where Y = A, C, T, G or a tetrahydrofuran analog of the natural AP site; N and N' = A, C, T or G; N and N’ form Watson-Crick base pair)

| Duplex | $\Delta S°$, cal/mol/K | $\Delta H°$, kcal/mol | $\Delta G°_{37}$, kcal/mol | $T_m$, °C |
|---|---|---|---|---|
| GTG/CAC | -221 ± 7 | -80.42 ± 2.35 | -8.47 ± 0.06 | 51.7 ± 0.2 |
| GTG/CCC | -135 ± 3.0 | -49.13 ± 0.80 | -7.94 ± 0.06 | 31.3 ± 0.4 |
| GTG/CTC | -167 ± 16 | -59.49 ± 4.8 | -8.03 ± 0.05 | 35 ± 0.3 |
| GTG/CGC | -180 ± 7.0 | -65.38 ± 2.3 | -8.27 ± 0.06 | 44 ± 0.2 |
| GTG/C(AP)C | -112 ± 11 | -40.7 ± 3.14 | -7.7 ± 0.17 | 22.2 ± 1 |
| CTC/GAG | -188 ± 3.0 | -69.14 ± 1.10 | -8.41 ± 0.05 | 49.2 ± 0.2 |
| CTC/GCG | -230 ± 4.0 | -76.59 ± 0.81 | -7.79 ± 0.31 | 25.5 ± 1.1 |
| CTC/GTG | -135 ± 2.0 | -49.83 ± 0.75 | -8.08 ± 0.14 | 36.7 ± 0.9 |
| CTC/GGG | -156 ± 0.1 | -56.76 ± 0.10 | -8.15 ± 0.04 | 39.4 ± 0.2 |
| ATA/TAT | -211 ± 4.0 | -74.68 ± 1.29 | -8.22 ± 0.06 | 42.2 ± 0.3 |
| ATA/TCT | -179 ± 8.0 | -61.53 ± 2.43 | -7.83 ± 0.12 | 27.3 ± 0.7 |
| ATA/TTT | -160 ± 7.0 | -56.2 ± 2.20 | -7.88 ± 0.11 | 29.1 ± 0.3 |
| ATA/TGT | -175 ± 24 | -62.29 ± 7.43 | -8.06 ± 0.10 | 36.1 ± 0.4 |
| TTT/AAA | -225 ± 12 | -79.85 ± 3.84 | -8.28 ± 0.07 | 44.4 ± 0.1 |
| TTT/ACA | -198 ± 18 | -67.35 ± 5.44 | -7.84 ± 0.12 | 27.5 ± 0.3 |
| TTT/ATA | -184 ± 5.0 | -63.12 ± 1.35 | -7.85 ± 0.06 | 27.9 ± 0.1 |
| TTT/AGA | -174 ± 16 | -60.68 ± 4.96 | -7.91 ± 0.10 | 30.3 ± 0.2 |

Table S5. Thermodynamic parameters of DNA duplexes with the central triplet N(**oxo-Ade$^{BZT}$**)N/N'YN', where Y = A, C, T, G or a tetrahydrofuran analog of the natural AP site; N and N' = A, C, T or G; N and N' form Watson-Crick base pair)

| Duplex | $\Delta S°$, cal/mol/K | $\Delta H°$, kcal/mol | $\Delta G°_{37}$, kcal/mol | $T_m$, °C |
|---|---|---|---|---|
| GXG/CAC | -185 ± 8 | -67.48 ± 2.52 | -8.33 ± 0.1 | 46.4 ± 0.2 |
| GXG/CCC | -226 ± 13 | -81.93 ± 4.04 | -8.46 ± 0.18 | 51.4 ± 1.1 |
| GXG/CTC | -217 ± 17 | -78.52 ± 5.47 | -8.41 ± 0.17 | 49.6 ± 0.4 |
| GXG/CGC | -192 ± 10 | -70.04 ± 3.40 | -8.39 ± 0.15 | 48.6 ± 0.1 |
| GXG/C(AP)C | -194 ± 17 | -70.54 ± 5.53 | -8.37 ± 0.19 | 47.8 ± 0.3 |
| CXC/GAG | -122 ± 2.0 | -45.75 ± 0.70 | -8.03 ± 0.08 | 34.9 ± 0.5 |
| CXC/GCG | -76 ± 6.0 | -30.61 ± 1.79 | -7.84 ± 0.38 | 27.5 ± 0.4 |
| CXC/GTG | -120 ± 0.1 | -46.32 ± 0.28 | -8.25 ± 0.26 | 43.4 ± 1.8 |
| CXC/GGG | -125 ± 1.0 | -47.35 ± 0.33 | -8.15 ± 0.12 | 39.3 ± 0.8 |
| AXA/TAT | -189 ± 23 | -66.97 ± 7.24 | -8.12 ± 0.13 | 38.3 ± 0.5 |
| AXA/TCT | -215 ± 17 | -76.14 ± 5.30 | -8.22 ± 0.05 | 42.2 ± 0.3 |
| AXA/TTT | -211 ± 11 | -74.17 ± 3.27 | -8.17 ± 0.08 | 40.3 ± 0.4 |
| AXA/TGT | -218 ± 15 | -76.37 ± 4.84 | -8.15 ± 0.18 | 39.6 ± 0.7 |
| TXT/AAA | -138 ± 6.0 | -49.51 ± 1.86 | -7.89 ± 0.03 | 29.5 ± 0.3 |
| TXT/ACA | -182 ± 25 | -63.97 ± 7.67 | -8.02 ± 0.07 | 34.5 ± 0.6 |
| TXT/ATA | -179 ± 14 | -63 ± 4.23 | -8 ± 0.16 | 33.5 ± 0.6 |
| TXT/AGA | -145 ± 20 | -52.06 ± 6.32 | -7.95 ± 0.02 | 31.6 ± 0.8 |

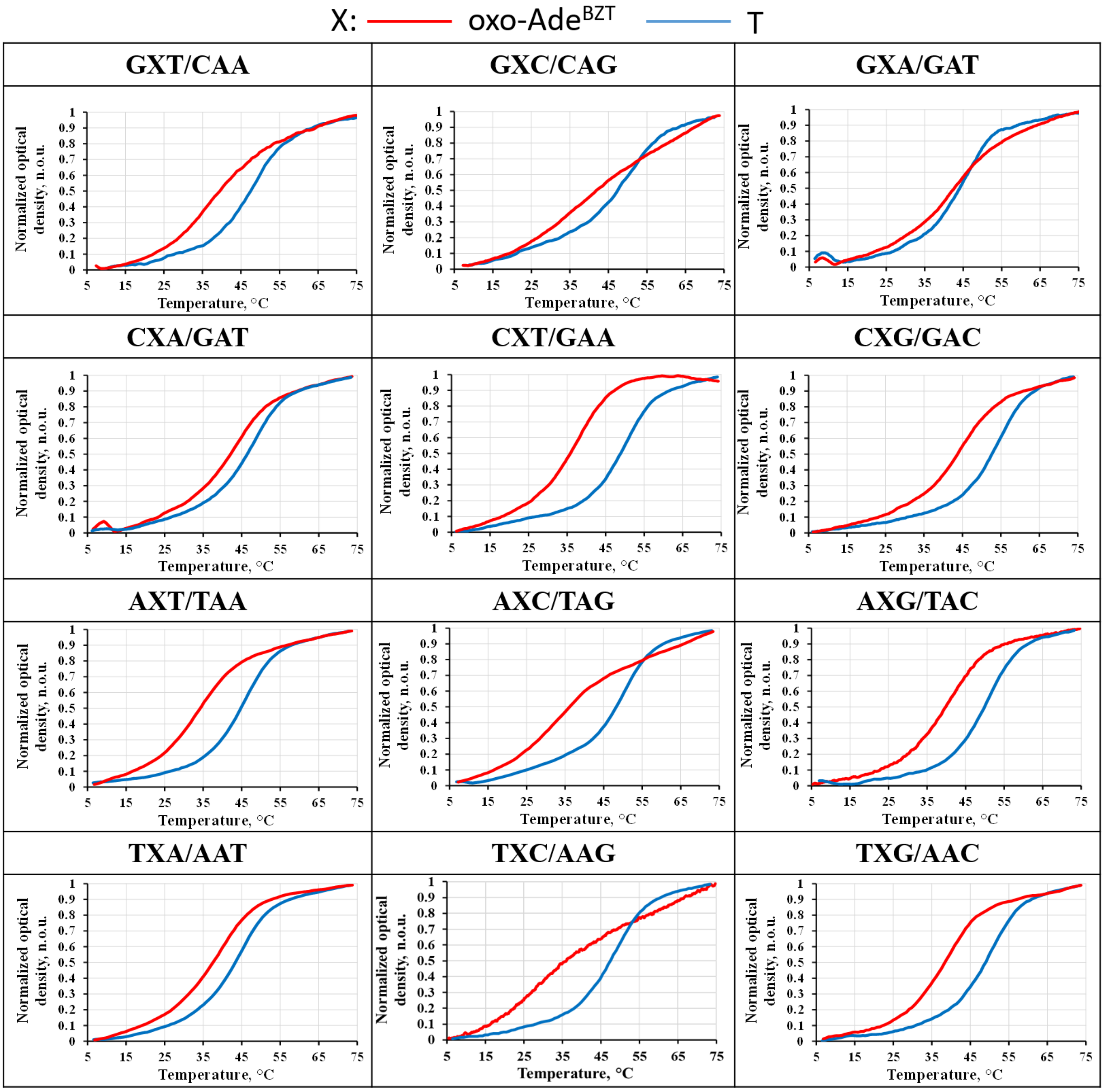

Figure S7. UV melting curves for modified and unmodified duplexes with the central triplet NXM/N'AM' (X = **oxo-Ade$^{BZT}$** or T; N, N', M, and M' = A, C, T or G; N /N' and M/M' form Watson-Crick base pair) registered at 260 nm.

Table S6. Thermodynamic parameters of DNA duplexes with the central triplet NTM/N'AM', where N, M, N' and M' = A, C, T or G; N /N' and M/M' form Watson-Crick base pair)

| Duplex | $\Delta S°$, cal/mol/K | $\Delta H°$, kcal/mol | $\Delta G°_{37}$, kcal/mol | $T_m$, °C |
|---|---|---|---|---|
| GTA/CAT | -183 ± 9 | -66.4 ± 2.77 | -8.29 ± 0.05 | 44.8 ± 0.2 |
| GTC/CAG | -124 ± 0 | -48.52 ± 0 | -8.42 ± 0.01 | 49.7 ± 0 |
| GTG/CAC | -221 ± 7 | -80.42 ± 2.35 | -8.47 ± 0.06 | 51.7 ± 0.2 |
| GTT/CAA | -268 ± 8 | -94.09 ± 2.51 | -8.34 ± 0.01 | 46.6 ± 0.2 |
| CTA/GAT | -200 ± 6 | -71.94 ± 1.83 | -8.31 ± 0.09 | 45.4 ± 0.4 |
| CTC/GAG | -188 ± 3 | -69.14 ± 1.1 | -8.41 ± 0.05 | 49.2 ± 0.2 |
| CTG/GAC | -211 ± 4 | -77.09 ± 1.1 | -8.47 ± 0.02 | 51.7 ± 0.2 |
| CTT/GAG | -220 ± 11 | -78.85 ± 3.59 | -8.37 ± 0.11 | 47.9 ± 0.1 |
| ATA/TAT | -211 ± 4 | -74.68 ± 1.29 | -8.22 ± 0.06 | 42.2 ± 0.3 |
| ATC/TAG | -221 ± 10 | -79.42 ± 3.36 | -8.37 ± 0.18 | 47.8 ± 0.3 |

| | | | | |
|---|---|---|---|---|
| ATG/TAC | -206 ± 8 | -74.62 ± 2.66 | -8.38 ± 0.15 | 48.4 ± 0.3 |
| ATT/TAA | -214 ± 9 | -76.44 ± 2.97 | -8.29 ± 0.07 | 44.6 ± 0.1 |
| TTA/AAT | -207 ± 8 | -73.56 ± 2.44 | -8.22 ± 0.02 | 42.1 ± 0.1 |
| TTC/AAG | -199 ± 6 | -71.74 ± 1.77 | -8.33 ± 0.06 | 46.2 ± 0.2 |
| TTG/AAC | -201 ± 8 | -72.76 ± 2.43 | -8.37 ± 0.08 | 47.8 ± 0.2 |
| TTT/AAA | -225 ± 12 | -79.85 ± 3.84 | -8.28 ± 0.07 | 44.4 ± 0.1 |

Table S7. Thermodynamic parameters of DNA duplexes with the central triplet N(**oxo-Ade$^{BZT}$**)M/N'AM', where N, M, N' and M' = A, C, T or G; N /N' and M/M' form Watson-Crick base pair)

| Duplex | $\Delta S°$, cal/mol/K | $\Delta H°$, kcal/mol | $\Delta G°_{37}$, kcal/mol | $T_m$, °C | $\Delta T_m$, °C[a] |
|---|---|---|---|---|---|
| GXA/CAT | -172 ± 23 | -62.94 ± 7.3 | -8.27 ± 0.15 | 44.0 ± 0.3 | -0.8 |
| GXC/CAG | -58 ± 13 | -25.72 ± 3.99 | -7.94 ± 0.38 | 31.2 ± 5.3 | -18.5 |
| GXG/CAC | -185 ± 8 | -67.48 ± 2.52 | -8.33 ± 0.1 | 46.4 ± 0.2 | -5.3 |
| GXT/CAA | -157 ± 7 | -56.84 ± 2.17 | -8.1 ± 0.06 | 37.6 ± 0.3 | -9.0 |
| CXA/GAT | -184 ± 10 | -66.32 ± 3.33 | -8.22 ± 0.2 | 42.0 ± 0.7 | -3.4 |
| CXC/GAG | -122 ± 2 | -45.75 ± 0.7 | -8.03 ± 0.08 | 34.9 ± 0.5 | -14.3 |
| CXG/GAC | -177 ± 6 | -64.22 ± 1.98 | -8.23 ± 0.11 | 42.5 ± 0.5 | -9.2 |
| CXT/GAA | -161 ± 14 | -57.74 ± 4.48 | -8.05 ± 0.07 | 35.7 ± 0.4 | -12.2 |
| AXA/TAT | -189 ± 23 | -66.97 ± 7.24 | -8.12 ± 0.13 | 38.3 ± 0.5 | -3.9 |
| AXC/TAG | -126 ± 12 | -46.12 ± 3.67 | -7.9 ± 0.09 | 29.7 ± 1 | -18.1 |
| AXG/TAC | -158 ± 6 | -57.51 ± 1.91 | -8.13 ± 0.02 | 38.7 ± 0.1 | -9.7 |
| AXT/TAA | -171 ± 10 | -60.64 ± 3.07 | -8.02 ± 0.09 | 34.5 ± 0.4 | -10.1 |
| TXA/AAT | -167 ± 14 | -60.07 ± 4.45 | -8.1 ± 0.14 | 37.6 ± 0.7 | -4.5 |
| TXC/AAG | -70 ± 5 | -27.91 ± 1.68 | -7.6 ± 0.09 | 18.2 ± 1.7 | -28.0 |
| TXG/AAC | -164 ± 9 | -58.74 ± 2.93 | -8.07 ± 0.01 | 36.4 ± 0 | -11.4 |
| TXT/AAA | -138 ± 6 | -49.51 ± 1.86 | -7.89 ± 0.03 | 29.5 ± 0.3 | -14.9 |

[a]$\Delta T_m = T_m^{modified\ dsDNA} - T_m^{unmodified\ dsDNA}$, where modified and unmodified dsDNA contains **oxo-Ade$^{BZT}$** or T, respectively ($T_m^{unmodified\ dsDNA}$ was taken from Table S6).

### 8. Determination of relative fluorescence quantum yield of the oxo-Ade$^{BZT}$ nucleotide in mODNs and their complexes

Quinine sulfate standard in 0.05 M $H_2SO_4$ was used to determine $\Phi_f$ of the **oxo-Ade$^{BZT}$** nucleotide in mODNs with the central triplet N(**oxo-Ade$^{BZT}$**)N, where N = A, C, T or G, and their complexes formed by mODNs with complementary strand bearing any natural nucleobase opposite **oxo-Ade$^{BZT}$**.

For example, Figure S8A shows the absorption spectra of quinine sulfate and ODN with the central triplet A(**oxo-Ade$^{BZT}$**)A. Absorption spectrum has pronounced well-spaced maxima associated with absorption of nucleotides and **oxo-Ade$^{BZT}$**. The absorption maximum of the **oxo-Ade$^{BZT}$** nucleotide is red-shifted by 7 nm compared to the **oxo-Ade$^{BZT}$** nucleoside (Figure S3). The intersection of the spectra is observed at a wavelength of 350 nm.

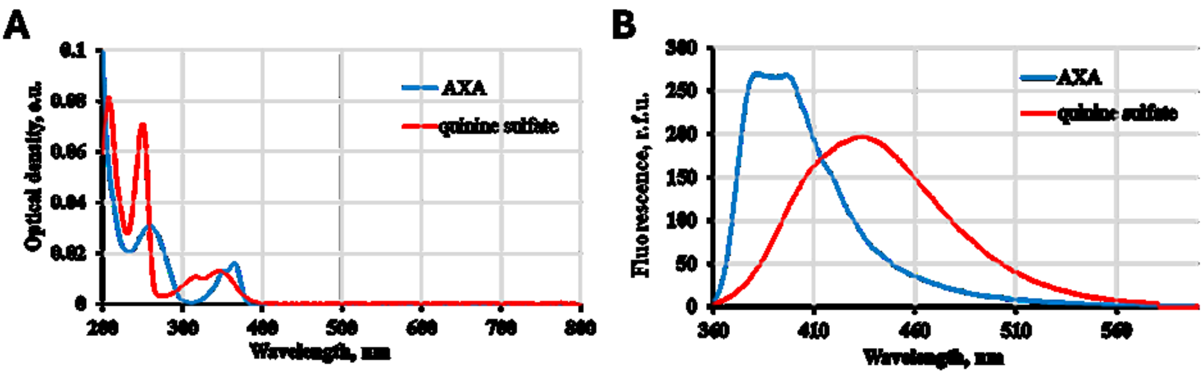

Figure S8. UV-Vis absorption spectra of ODN with the central triplet A(**oxo-Ade$^{BZT}$**)A at a concentration of $1.0\times10^{-6}$ M in PBS (pH 7.4) and quinine sulfate at a concentration $1.1\times10^{-5}$ M in 0.05M $H_2SO_4$ (A). Fluorescence emission spectra of mODN and quinine sulfate at a concentration of $1.2\times10^{-5}$ and $9.0\times10^{-7}$ M, respectively, at an excitation wavelength of 350 nm (B).

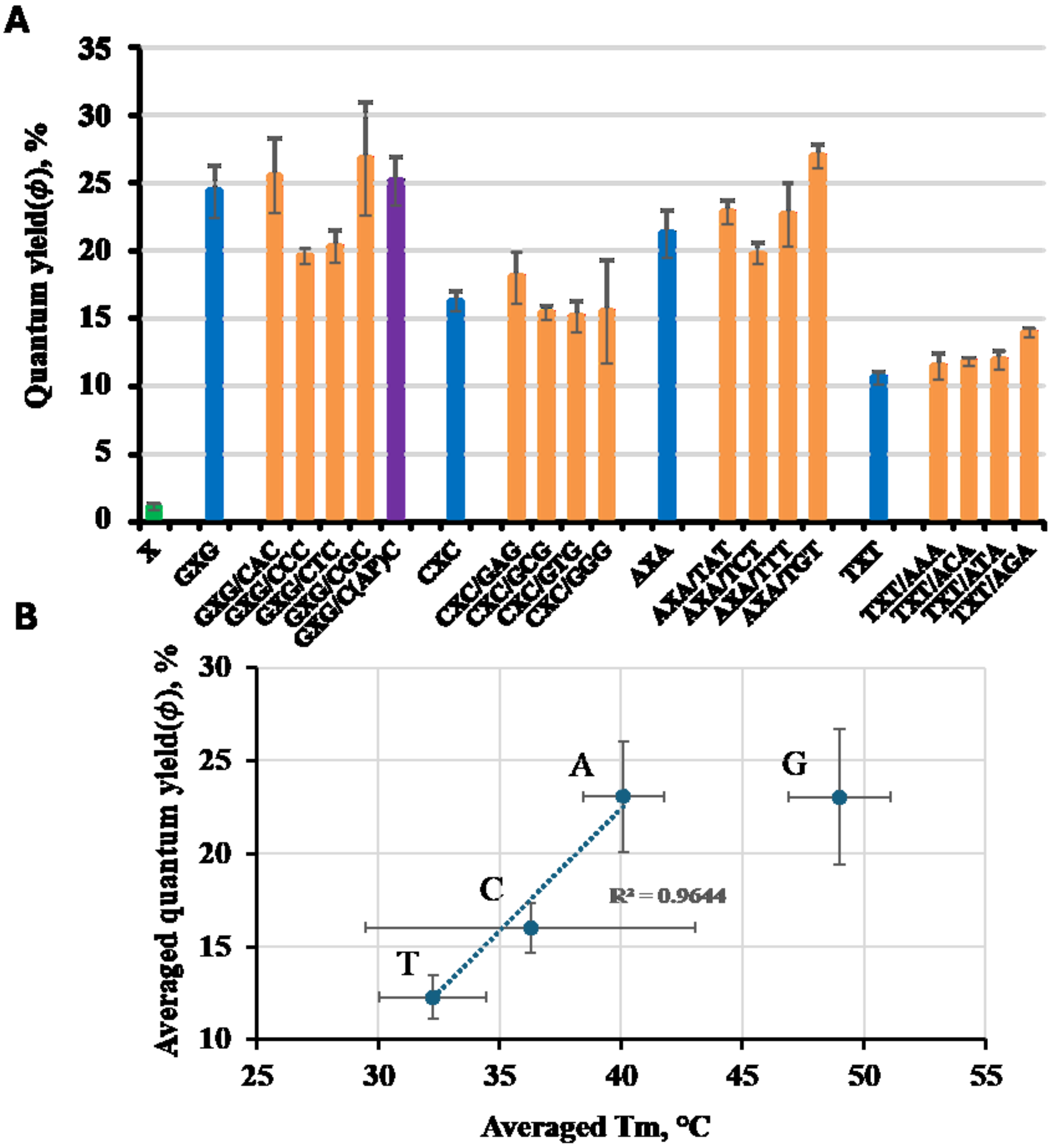

Figure S9. Fluorescence quantum yield ($\Phi_f$) values of the **oxo-Ade$^{BZT}$** nucleoside (green), mODNs (blue) and their complexes with complementary strands bearing any natural nucleobase (orange) or a tetrahydrofuran analog of the natural AP site (purple) opposite **oxo-Ade$^{BZT}$** (A). Concentrations of modified and complementary ODNs were 1.2 and 1.5·10$^{-6}$ M, respectively, in 0.01 M PBS buffer (pH 7.4). The dependence of the averaged fluorescence quantum yield on the averaged melting temperature ($T_m$) of the duplex for each pair of flanking bases (data from Table 1) (B).

### 9. Molecular modeling

Molecular dynamics (MD) simulation was performed in AMBER20 and AMBER24 software packages, with the use of parallel calculations on central (CPU) and graphical (GPU) processors. The OL21 force field was used for natural nucleotides within ODNs[7], and the gaff2 force field was used for the **oxo-Ade$^{BZT}$** [8]. MD simulation was performed in an explicit water shell (OPC model)[7]. Sodium ions (ionsjc_opc force field) were added to periodic simulation box to neutralize the negative charges of the duplexes. The duplexes were placed in a cubic cell with a distance from the simulated molecule to the cell boundaries of at least 14 Å.

The simulation was performed following reported procedure[9] that included five stages: (1) minimization of the system with a fixed **oxo-Ade$^{BZT}$** nucleoside/mODN/duplex (10,000 minimization steps, constant restraint force of 10 kcal/mol/Å$^2$) (PMEMD.MPI), (2) heating of the system from 0 to 300 K with a fixed **oxo-Ade$^{BZT}$** nucleoside/mODN/duplex for 2 ns with time step of 0.1 fs (constant restraint force of 10 kcal/mol/Å$^2$) (PMEMD.CUDA), (3) equilibration of the system density at the constant pressure of 1 bar for 400 ps (SANDER.MPI), (4) equilibration of the system density at the constant pressure of 1 bar and a temperature of 300 K for 2 ns (PMEMD.CUDA); (5) MD simulations for 300 ns at a constant pressure of 1 bar and a temperature of 300 K (PMEMD.CUDA).

The cpptraj[10] module of the AmberTools package was used for trajectory analysis: for cluster analysis of trajectories, calculation root mean square deviations (RMSD) and root mean square fluctuations of atoms positions (RMSF) of the mODN or its duplex and to monitor the librations of the dihedral angels. The hierarchical cluster analysis methods were used to find the most represented structures in the MD trajectory by analyzing the coordinates of the heavy atoms of the system, which was used to obtain the ten most representative structures in the MD trajectory. Molecular graphics were prepared in the UCSF Chimera package[11].

The 2'-deoxyribose residue was replaced by an isopropyl group and conducted the structure optimization using two isomers with a 180° rotation around d1 and two conformers with a 180° rotation around d2 as the starting points for quantum mechanical calculations (QM) (Figure S10). The QM calculations were performed using density functional theory (DFT) at the B3LYP-D3BJ/def2-TZVP theory level. After optimization, the isomer 1 with the sulfur atom located above the nitrogen atom with a lone electron pair proven to be energetically most favorable.

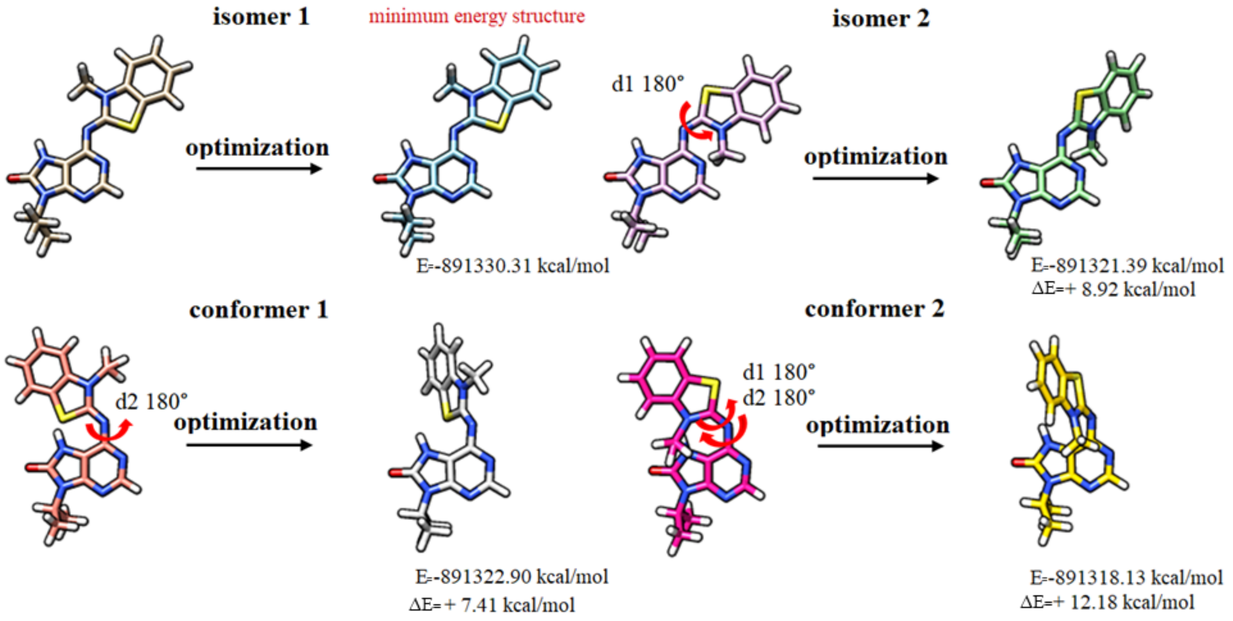

Figure S10. The results of the structure optimization of isomers 1,2 and conformers 1,2 and the energy of the optimized structures. Atom coloring: carbon - varying color, oxygen - red, nitrogen - blue, sulfur - yellow, hydrogen – white. The red arrow shows a 180° rotation around the bonds described by d1 and/or d2, respectively, relative to isomer 1. ΔE was calculated as the difference between the structure with the lowest energy and the other structures.

To analyze energy profile, we evaluated the potential energy profile of the molecule during rotation around a single bond, represented by d2 (Figure S11A,B). Rotation of d2 from 0 to 360° in 10° increments reveals steric hindrance: the methyl group of the *N*-methylbenzo[d]thiazolyl moiety cannot be positioned near the N-H bond of 7,8-dihydro-8-oxopurine. This demonstrates that isomer 2 with a 360° rotation around d2 is energetically unfavorable (Figure S11A,B). Two states were found in which the N-methylbenzo[d]thiazolyl moiety was perpendicular to the N-H bond or the nitrogen atom with the lone electron pair of 7,8-dihydro-8-oxopurine, in which the molecule could not overcome the energetic barrier (Figure S11C,D).

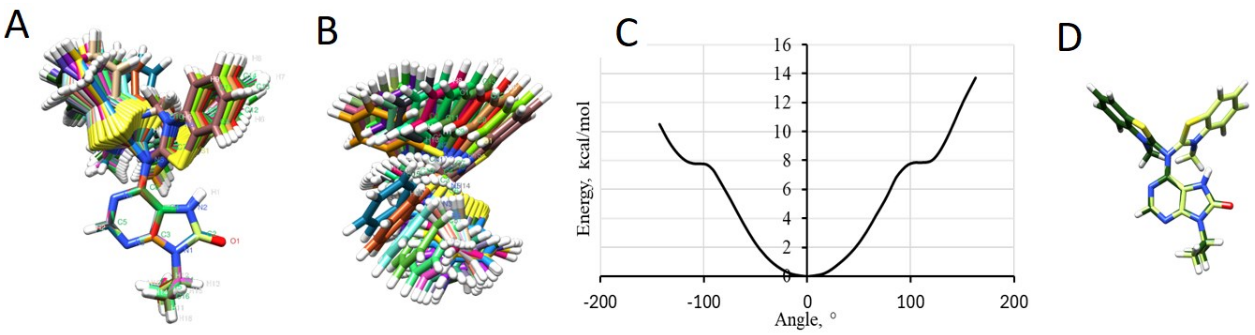

Figure S11. Superposition of the structures obtained by rotation around a single bond represented by d2: side view (A) and top view (B). Dependence of energy of the molecule on rotation around a single bond represented by d2 (C) and two structures with -100 and +100° rotations corresponding to the minima ("shelves") in the energy profile.

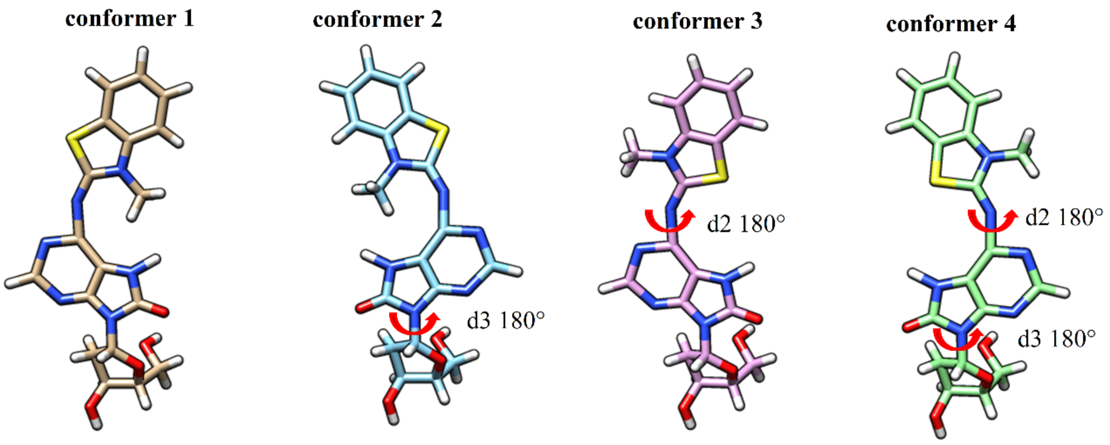

Figure S12. The structure of possible four conformers of the **oxo-Ade$^{BZT}$** nucleos(t)ide (the phosphate group is not shown). Atom coloring: carbon - varying color, oxygen - red, nitrogen - blue, sulfur - yellow, hydrogen – white. The red arrow shows a 180° rotation around the bonds described by d2 and/or d3, respectively, relative to conformer 1.

Due to the restricted rotation around d2, we employed MD simulations to analyze the conformational space of the modified nucleobase. Regardless of the initial conformer (Figure S12), the most represented structures of the **oxo-Ade$^{BZT}$** nucleoside observed in MD simulations are practically flat (maximum rotation of 10° along the bond described by d2) and contains the sulfur atom located above the nitrogen atom with a lone electron pair (Figure S13). The results are consistent with those of the structure optimization (Figure S10).

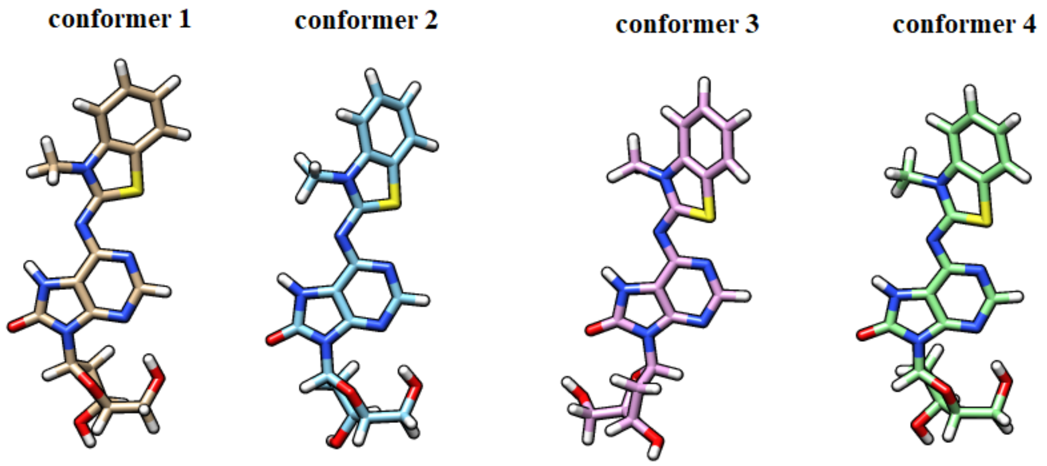

Figure S13. The most represented structures of the **oxo-Ade$^{BZT}$** nucleoside in MD trajectories. Atom coloring: carbon - varying color, oxygen - red, nitrogen - blue, sulfur - yellow, hydrogen – white.

According to MD simulation results for all four conformers of ODN with the central triplet G**oxo-Ade$^{BZT}$**G, heterocyclic system of the **oxo-Ade$^{BZT}$** nucleotide remains planar, with the orientation *N*-methylbenzo[d]thiazole relative to 7,8-dihydro-8-oxopurine for all initial conformers, as shown in Figure S12, maintained throughout most of the MD simulation (Figure S14). No significant rotations of the heterocyclic system of the **oxo-Ade$^{BZT}$** nucleotide around d1 and d2 were observed.

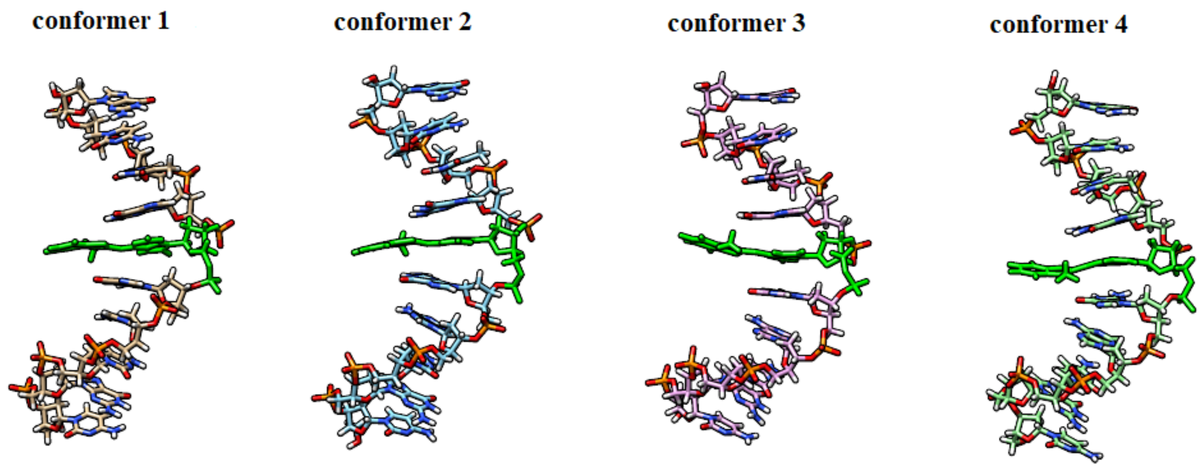

Figure S14. The most represented structures of ODNs with the central triplet G**oxo-Ade$^{BZT}$**G in MD trajectories. Atom coloring: carbon - varying color, oxygen - red, nitrogen - blue, hydrogen – white. The **oxo-Ade$^{BZT}$** nucleotide is indicated in green.

For the duplexes with the central base pair triplet G**oxo-Ade$^{BZT}$**G/CAC, where 2'-deoxyadenosine nucleotide is in both the anti and syn conformation, **oxo-Ade$^{BZT}$** pushes the opposite adenine out of the DNA double helix and occupies all the space between the flanking base pairs during the MD simulation. For conformer 1 for the both conformations of the opposite 2'-deoxyadenosine nucleotide, the **oxo-Ade$^{BZT}$** nucleotide is not a planar. In this way, 3-methylbenzo[d]thiazole gets involved in stacking interactions with the 2'-deoxycytidine nucleotide (nucleotide 14) that 5'-flanks the opposite 2'-deoxyadenosine, whereas the 2'-deoxyadenosine nucleotide itself flipped out of the major groove of the DNA double helix. For conformer 4 for the both conformations of the opposite 2'-deoxyadenosine nucleotide, 3-methylbenzo[d]thiazole is in stacking with the opposite adenine, which is accompanied by a deviation from planarity of the heterocyclic system of the **oxo-Ade$^{BZT}$** nucleotide (Figures S15 and S16). The conformational stability and flexibility analysis of the structures was performed based on the root mean square deviation (RMSD) and root mean square fluctuation (RMSF) of atoms from the initial structures along the MD trajectory. The RMSD and RMSF values of the **oxo-Ade$^{BZT}$** nucleoside, ODNs and duplexes are shown in Figures S17 and S18, respectively. The RMSD ranges from 0.5 to 2.75 Å, from 1 to 8 Å and from 1 to 7.5 Å for the **oxo-Ade**$^{BZT}$ nucleoside, ODNs and duplexes, respectively (Figure S17). Analysis of atomic fluctuations relative to the equilibrium position averaged along the MD trajectory for ODNs and duplexes shows that the RMSF values are higher at the ODN and duplex termini (nucleotides 1, 10, 11 and 20) (Figure S18). Numbering (5'→3') from 1 to 10 for NXN strand (**oxo-Ade**$^{BZT}$ is the nucleotide 6) and from 11 to 20 for N'YN' strand (the opposite 2'-deoxyadenosine is the nucleotide 15) was used. For nucleotide 6, the RMSF value in ODNs and duplexes is 3 Å and from 1 to 2.5 Å, respectively. This value is 0.5 Å higher than that of the stable duplex regions. The significant increase in the RMSF value for nucleotide 15 in the duplexes is due to the pushing adenine out of the DNA major groove by **oxo-Ade**$^{BZT}$ and its high conformational mobility due to the absence of steric hindrances associated with interactions with adjacent nucleotides on the same and opposite strands.

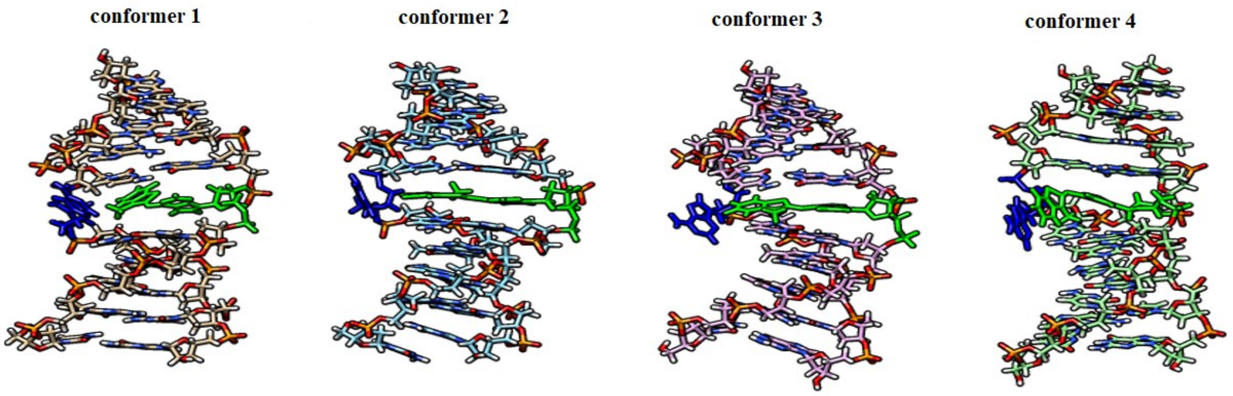

Figure S15. The most represented structures of duplexes with base pair triplet G**oxo-Ade**$^{BZT}$G/CAC obtained by hierarchical analysis of MD trajectory, where the 2'-deoxyadenosine nucleotide is in the anti conformation, in MD trajectories. Atom coloring: carbon - varying color, oxygen - red, nitrogen - cyan, hydrogen – white. The **oxo-Ade**$^{BZT}$ nucleotide and the opposite 2'-deoxyadenosine nucleotide are indicated in green and blue, respectively.

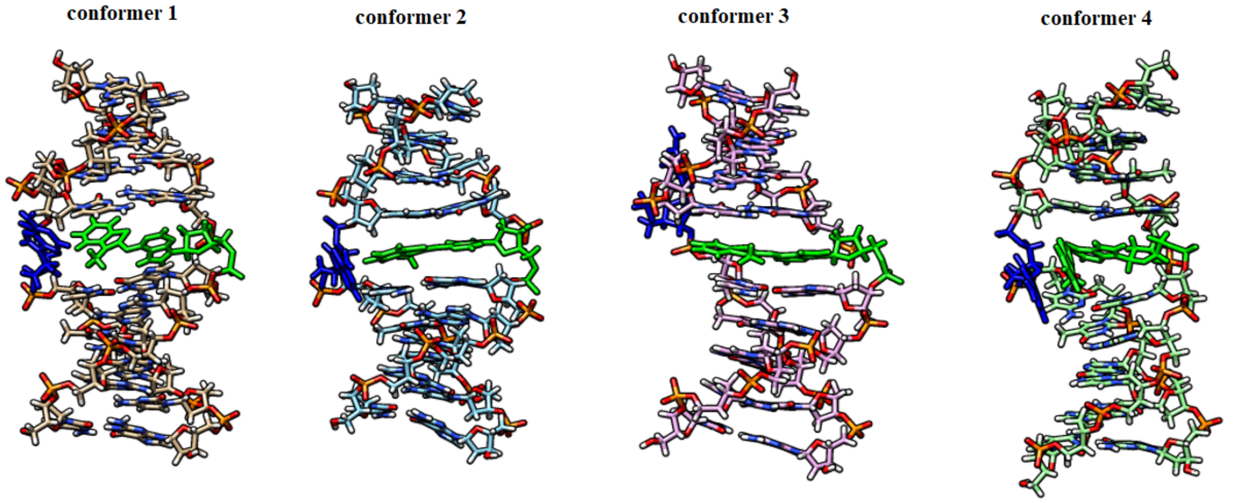

Figure S16. The most represented structures of duplexes with base pair triplet G**oxo-Ade**$^{BZT}$G/CAC obtained by hierarchical analysis of MD trajectory, where the 2'-deoxyadenosine nucleotide is in the syn conformation, in MD trajectories. Atom coloring: carbon - varying color, oxygen - red, nitrogen - cyan, hydrogen – white. The **oxo-Ade**$^{BZT}$ nucleotide and the opposite 2'-deoxyadenosine nucleotide are indicated in green and blue, respectively.

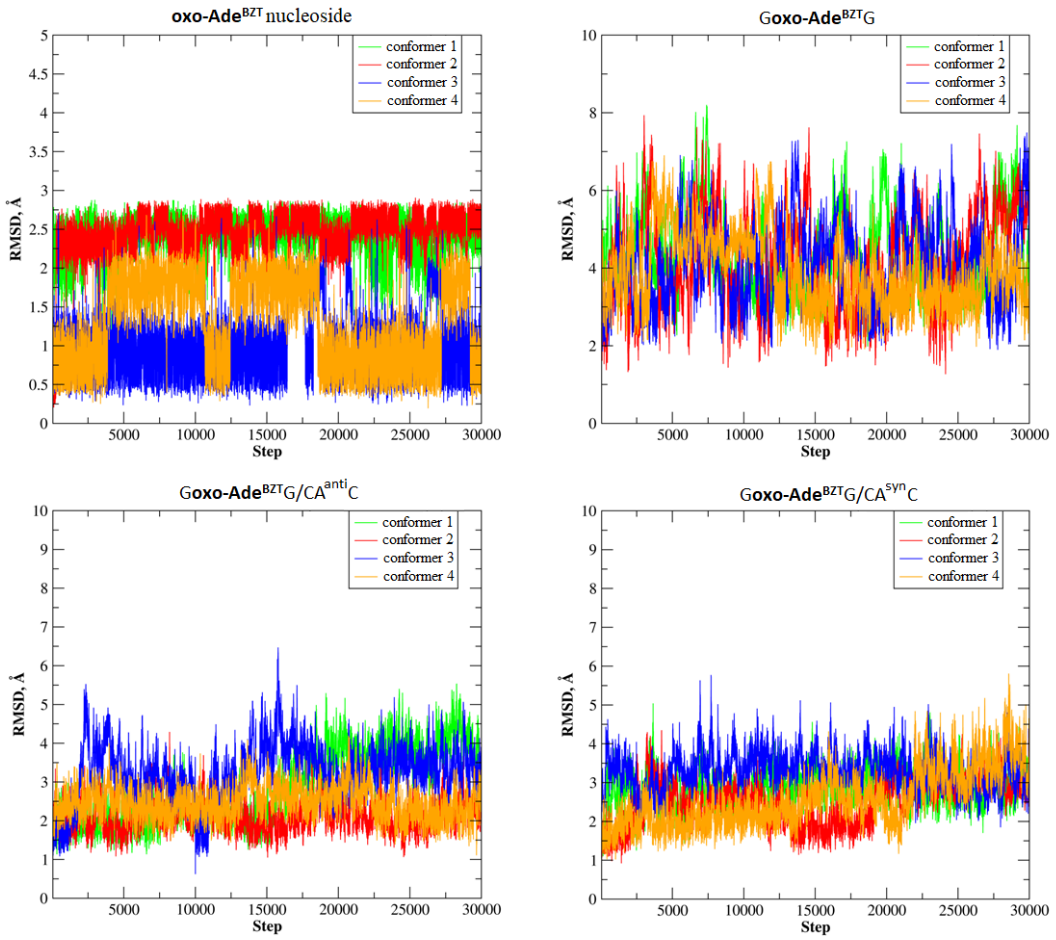

Figure S17. RMSD values for the **oxo-Ade**$^{BZT}$ nucleoside and nucleotide within ODN and duplexes calculated for all heavy atoms along the MD trajectories.

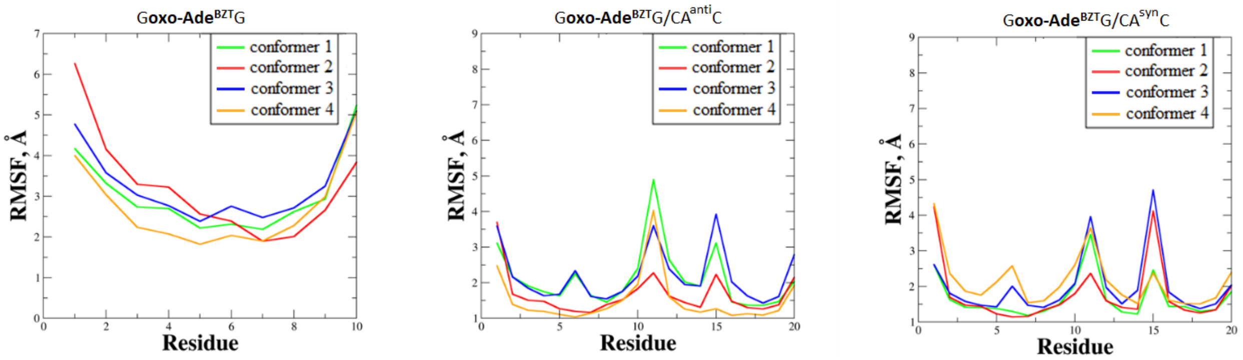

Figure S18. RMSF values for residues within ODN and duplexes.

To evaluate the flexibility of the **oxo-Ade**$^{BZT}$ nucleoside as well as the **oxo-Ade**$^{BZT}$ nucleotide within ODN and duplexes, the dihedral angles d1, d2, and d3 were analyzed. The standard deviations of each dihedral angle are presented in Figures S23 (for d1), S28 (for d2), and S33 (for d3) and are summarized in Tables S8 and S9.

**Dihedral angle d1**

Two peaks are observed for all four conformers of the **oxo-Ade**$^{BZT}$ nucleoside, with maxima at 160° and 185° (Figure S19). The distance between maxima is no more than 25° for all four conformers (Figure S19A), which in agreement with the most representative structures founded in MD trajectories of the **oxo-Ade**$^{BZT}$ nucleoside (Figure S13). For ODN with the central triplet G**oxo-Ade**$^{BZT}$G, one peak is observed for all four conformers (Figure S20), with a maximum of 180°. The peak width and amplitude are the same for all four conformers and the distribution width decreases by 1.8 times compared to the **oxo-Ade**$^{BZT}$ nucleoside (Figure S20B). For the duplex with the central base pair triplet G**oxo-Ade**$^{BZT}$G/CA$^{anti}$C, where 2'-deoxyadenosine nucleotide is in the anti conformation, the peak maxima and their amplitudes are identical for conformers 1 and 2 (Figure S21). For conformer 4, the peak maximum is 325°. For conformer 3, a 25° rotation is observed at 100 ns of the MD simulation, with a total duration of no more than 30 ns. Significant changes in the d1 value for conformer 3 (Figure S21A) are associated with structural changes (Figure S34A). For most of the MD simulation, the **oxo-Ade$^{BZT}$** nucleotide adopts a planar conformation and occupies the entire space between the flanking base pairs (Figure S34A). The 25° rotation with a total duration of no more than 35 ns is associated with stacking interactions between 3-methylbenzo[d]thiazole and the opposite adenine (Figure S34A). For the duplex with the central base pair triplet G**oxo-Ade$^{BZT}$**G/CA$^{syn}$C, where 2'-deoxyadenosine nucleotide is in the syn conformation, a single peak is observed with a maximum at 175° for conformer 1, 177° for conformers 2 and 3, and 325° for conformer 4 (Figure S22A). The peak width and amplitude are the same for conformers 1 and 2. For conformer 4, a 60° rotation is observed. Significant changes in the d1 value, as in the case of conformer 3 with the opposite 2'-deoxyadenosine nucleotide in anti conformation (Figure S21A), are associated with structural changes (Figure S34C). For most of the MD simulation, the **oxo-Ade$^{BZT}$** nucleotide adopts a planar conformation and occupies the entire space between the flanking base pairs (Figure S34C). The 70° rotation with a total duration of no more than 15 ns is associated with stacking interactions between 3-methylbenzo[d]thiazole and the opposite adenine (Figure S34C). In summary, the mobility of d1 in the**oxo-Ade$^{BZT}$** nucleoside is 1.8 times higher than that in ODN and duplexes (Figure S23).

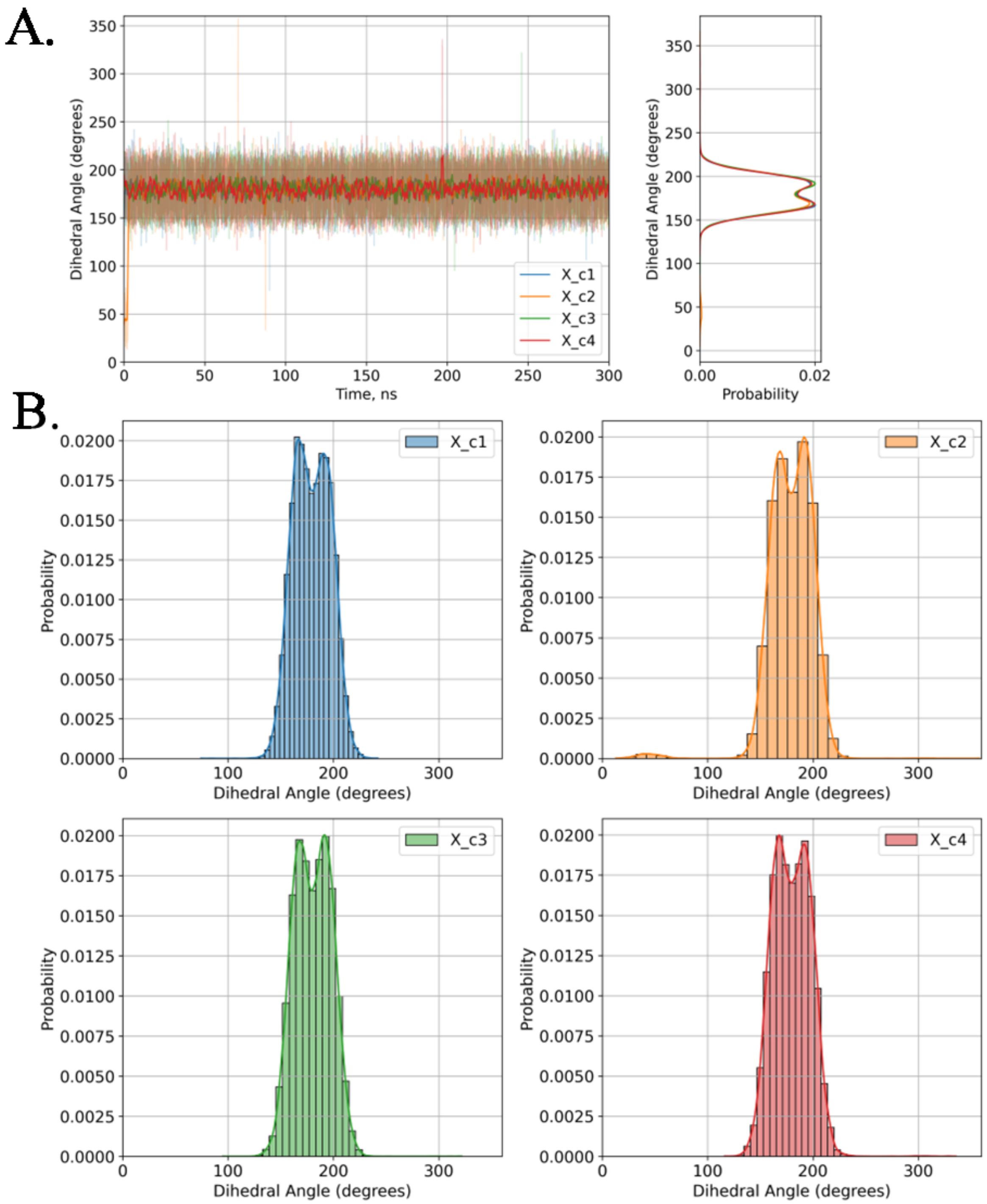

Figure S19. Evolution (A) and distribution (B) of dihedral angle d1 for four conformers of the **oxo-Ade**$^{BZT}$ nucleoside: conformer 1 (blue), conformer 2 (orange), conformer 3 (green), conformer 4 (red).

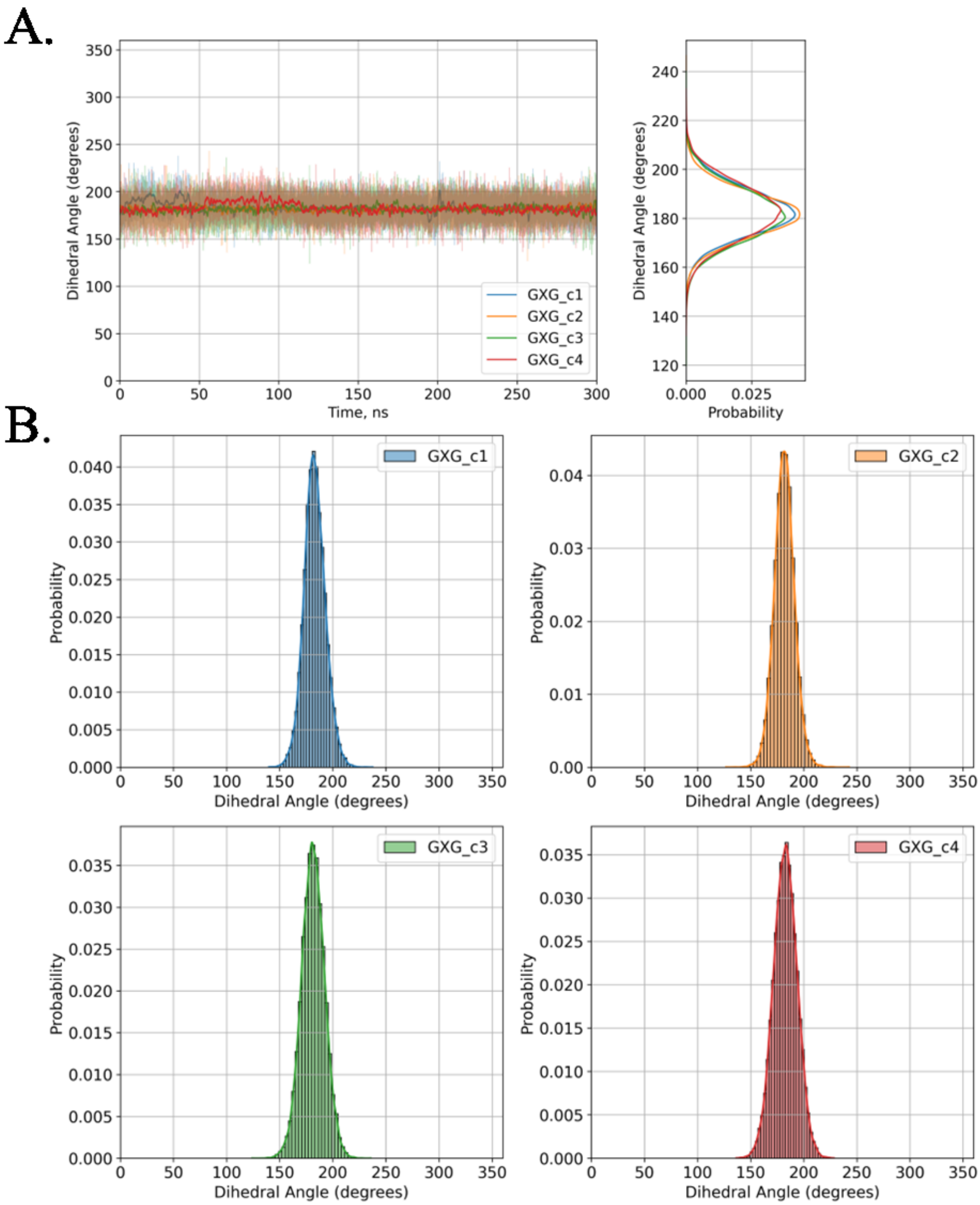

Figure S20. Evolution (A) and distribution (B) of dihedral angle d1 for four conformers of the **oxo-Ade**$^{BZT}$ nucleotide within ODN with the central triplet G**oxo-Ade**$^{BZT}$G: conformer 1 (blue), conformer 2 (orange), conformer 3 (green), conformer 4 (red).

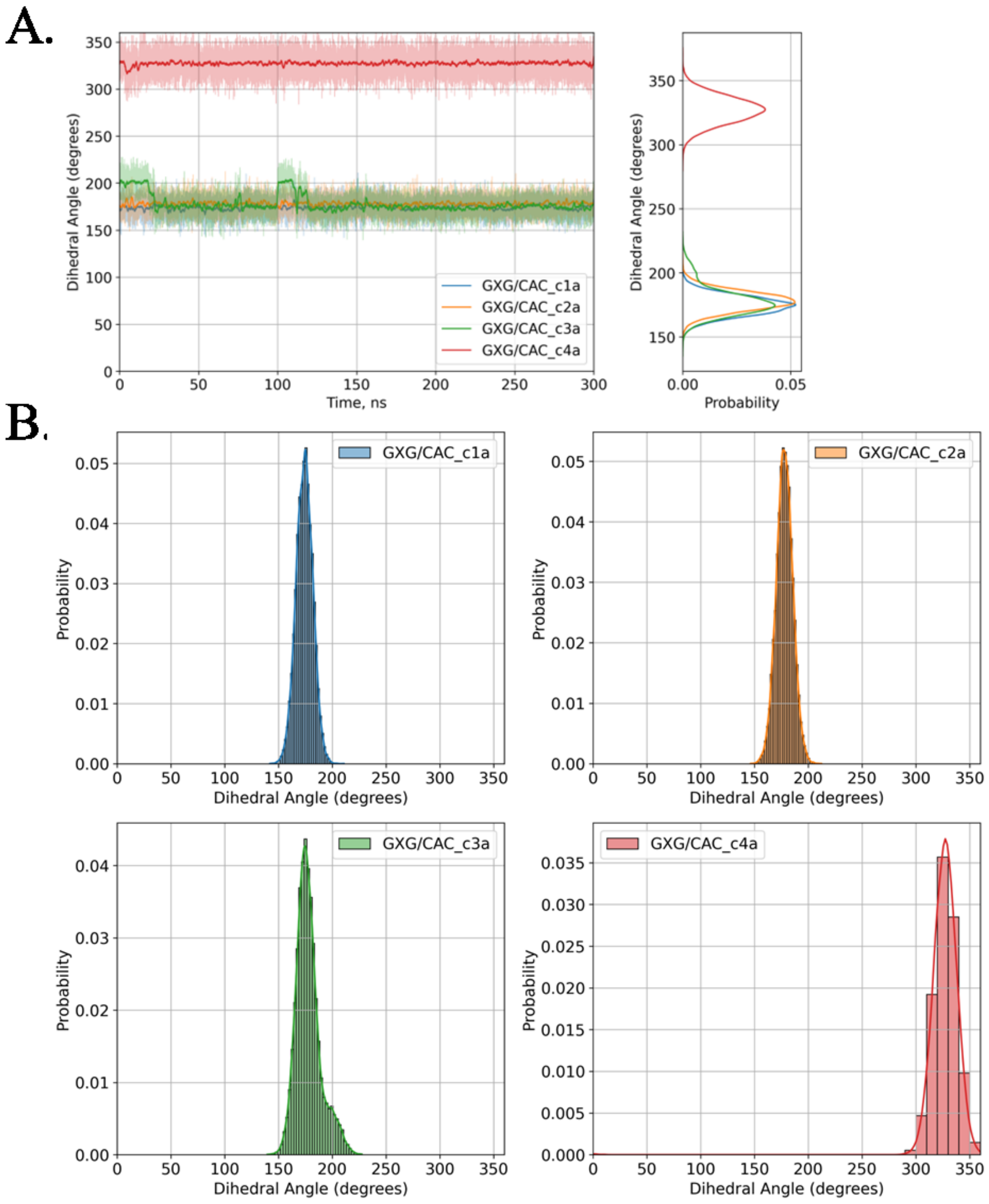

Figure S21. Evolution (A) and distribution (B) of dihedral angle d1 for four conformers of the **oxo-Ade**$^{BZT}$ nucleotide within the duplex with the central base pair triplet G**oxo-Ade**$^{BZT}$G/CA$^{anti}$C, where 2'-deoxyadenosine nucleotide is in the anti conformation: conformer 1 (blue), conformer 2 (orange), conformer 3 (green), conformer 4 (red).

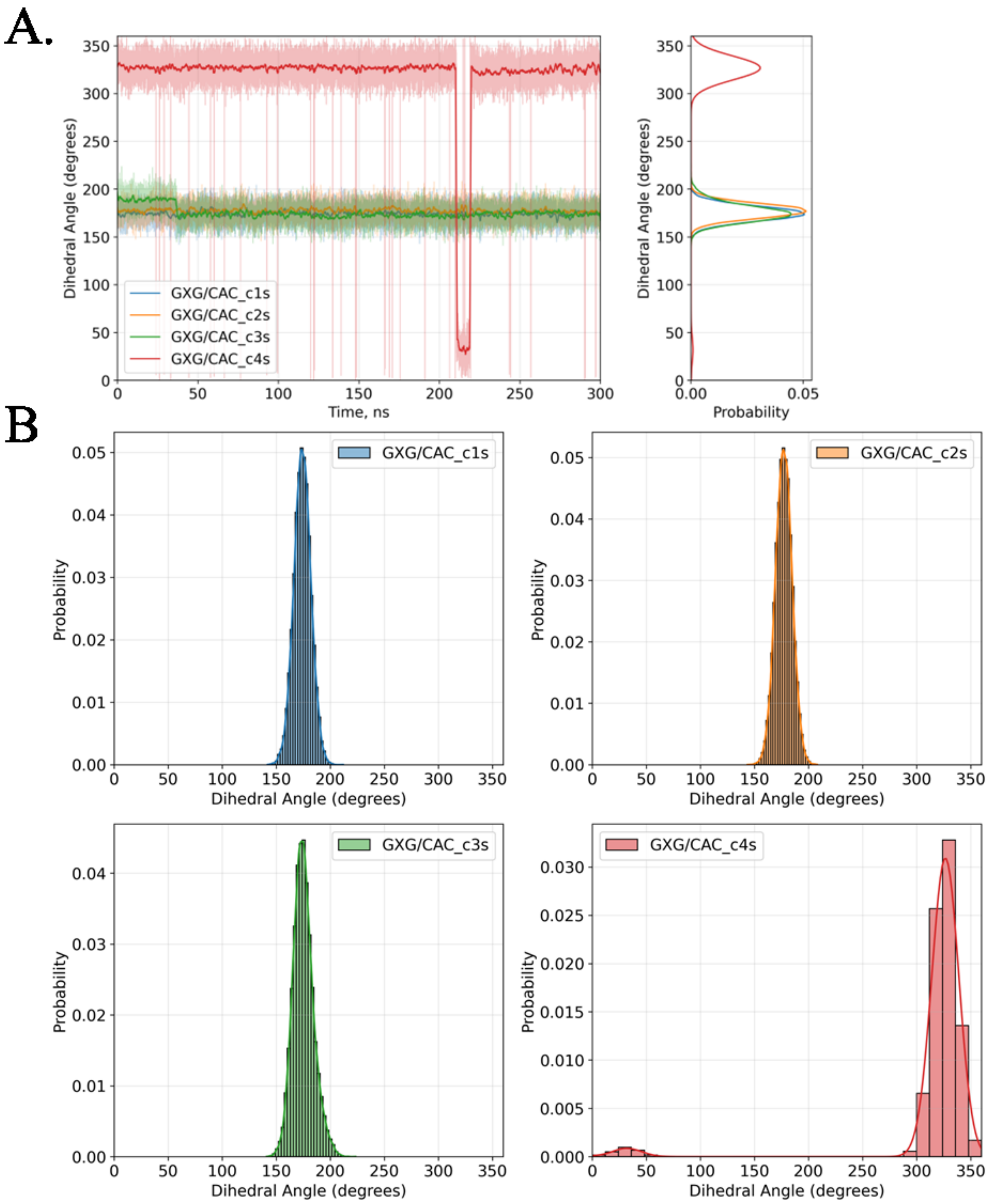

Figure S22. Evolution (A) and distribution (B) of dihedral angle d1 for four conformers of the **oxo-Ade**$^{BZT}$ nucleotide within the duplex with the central base pair triplet G**oxo-Ade**$^{BZT}$G/CA$^{syn}$C, where 2'-deoxyadenosine nucleotide is in the syn conformation: conformer 1 (blue), conformer 2 (orange), conformer 3 (green), conformer 4 (red).

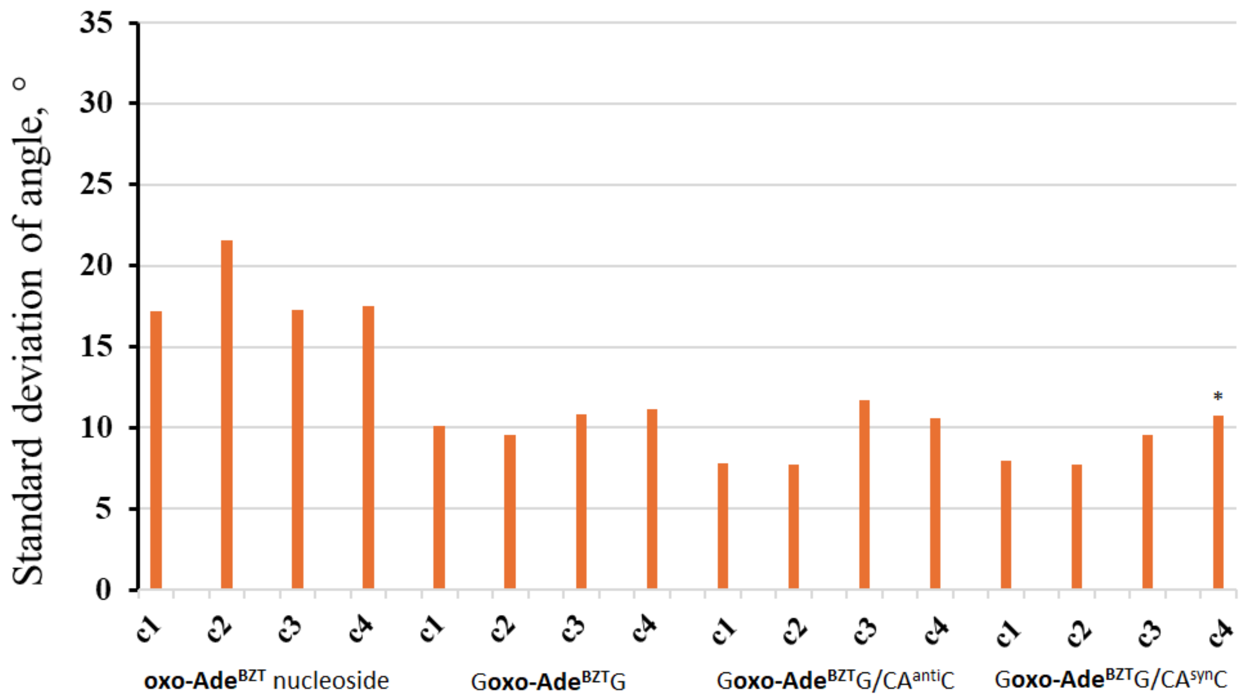

Figure S23. Standard deviation of d1 within the **oxo-Ade**$^{\mathbf{BZT}}$ nucleoside, ODN and duplexes for four conformers. * since two stable states with different angles (Figure S22) were observed during the MD simulation, the state with the longest occupation time in the MD simulation was taken for the data; both values are presented in Table S8.

**Dihedral angle d2**

Two peaks are observed for all four conformers of the o**xo-Ade**$^{BZT}$ nucleoside with maxima at 160° and 210° (Figure S24A). The distance between the maxima depending on the conformer is the same and is 50° for all four conformers (Figure S24A). For ODN with the central triplet G**oxo-Ade**$^{BZT}$G, one peak is observed with a maximum of 182° for conformers 1, 2, and 3, and 185° for conformer 4 (Figure S25). The peak width and amplitude are the same for all four conformers. For the duplex with the central base pair triplet G**oxo-Ade**$^{BZT}$G/CA$^{anti}$C, where 2'-deoxyadenosine nucleotide is in the anti conformation, one peak is observed with a maximum of 310° for conformers 1 and 4, and 178° for conformers 2 and 3 (Figure S26). Conformer 3 exhibits a peak at 50°, which corresponds to the transition of the **oxo-Ade**$^{BZT}$ nucleotide at 0-20 and 100-115 ns in the MD simulation. Similar to d1, significant changes in the d2 value for conformer 3 (Figure S26) are associated with structural changes (Figure S34A). For most of the MD simulation, the **oxo-Ade**$^{BZT}$ nucleotide adopts a planar conformation and occupies the entire space between the flanking base pairs (Figure S34A). The 120° rotation with a total duration of no more than 35 ns is associated with stacking interactions between 3-methylbenzo[d]thiazole and the opposite adenine (Figure S34A). For the duplex with the central base pair triplet G**oxo-Ade**$^{BZT}$G/CA$^{syn}$C, where 2'-deoxyadenosine nucleotide is in the syn conformation, one peak is observed with a maximum of 303° and 180° for conformer 1 and 2, respectively (Figure S27). For conformer 3, two peaks are observed with a maximum of 175° (the second peak at 45°), which corresponds to the rotation of heterocyclic system of the **oxo-Ade**$^{BZT}$ nucleotide at 0-40 ns in the MD simulation, and with a maximum of 320° (the second peak at 175°), which corresponds to the rotation of heterocyclic system of the **oxo-Ade**$^{BZT}$ nucleotide at 215-300 ns in the MD simulation (Figure S27A). Significant changes in the d2 value are associated with structural changes. For conformer 3, **oxo-Ade**$^{BZT}$ nucleotide retains a planar structure, but rotates toward the center of the hydrophobic pocket between flanking base pairs (Figure S34B). For conformer 4, the **oxo-Ade**$^{BZT}$ nucleotide adopts a planar conformation and pushes adenine out of the DNA double helix (Figure S34C). Unlike conformer 4 with the **oxo-Ade**$^{BZT}$ nucleotide in the anti conformation, the conformer maintains stacking interactions with the opposite adenine throughout the MD simulation and does not transition to a planar structure. To summarize this part, the mobility of d2 in the **oxo-Ade**$^{BZT}$ nucleoside is 1.8 times higher than that in ODN and duplexes depending on the conformer, but no significant changes between ODN and duplexes are observed (Figure S28).

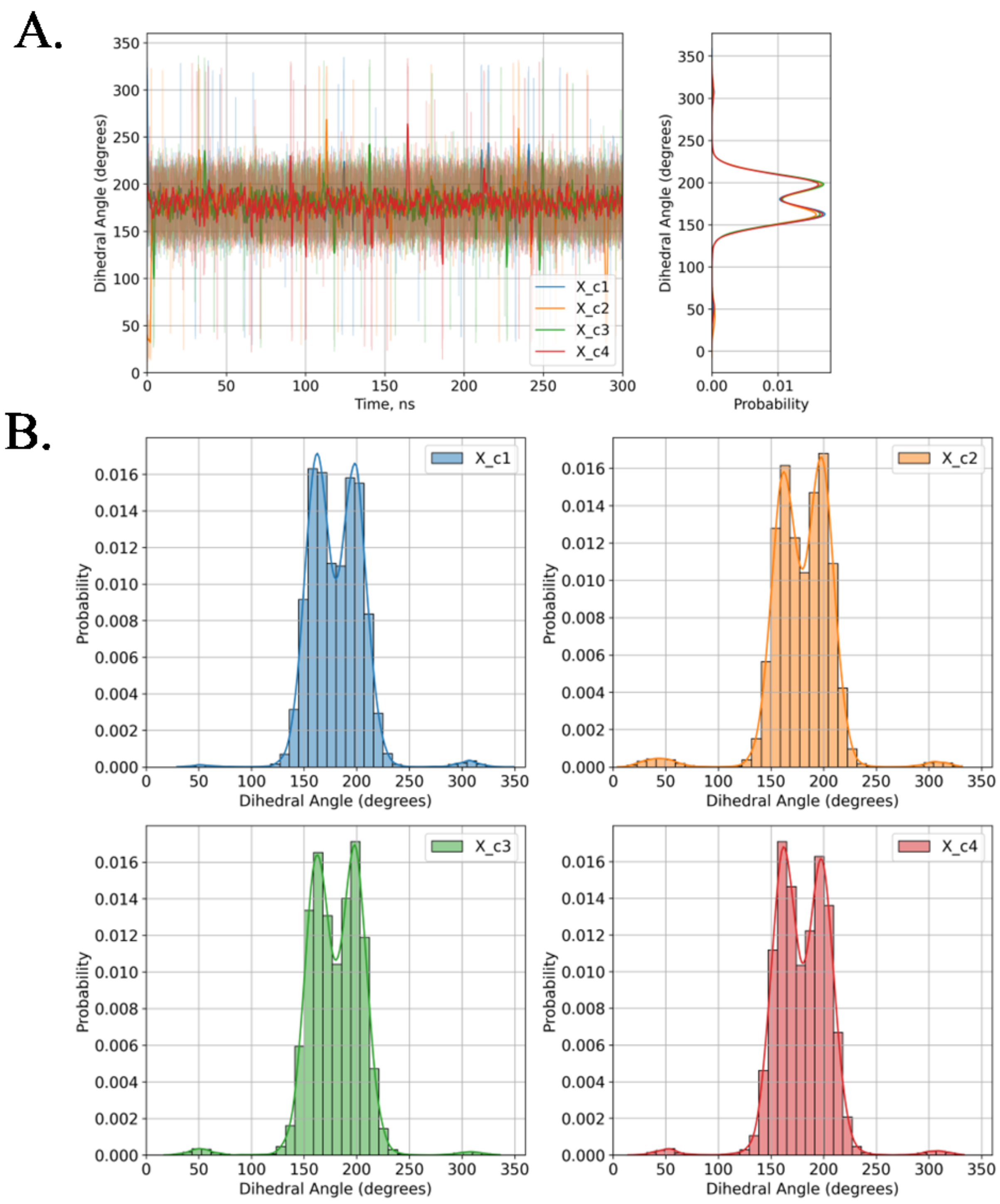

Figure S24. Evolution (A) and distribution (B) of dihedral angle d2 for four conformers of the **oxo-Ade$^{BZT}$** nucleoside: conformer 1 (blue), conformer 2 (orange), conformer 3 (green), conformer 4 (red).

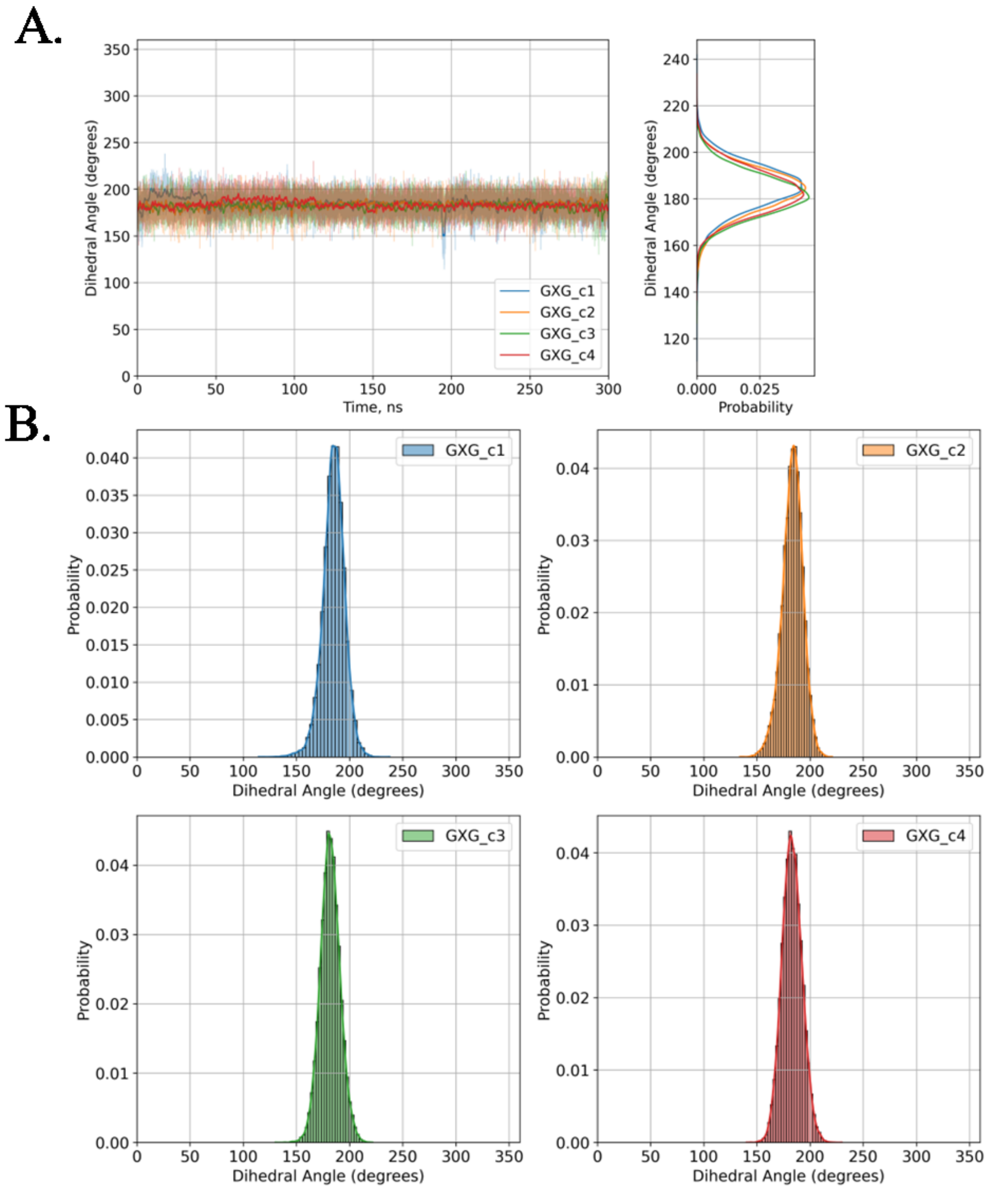

Figure S25. Evolution (A) and distribution (B) of dihedral angle d2 for four conformers of the **oxo-Ade**$^{BZT}$ nucleotide within ODN with the central triplet G**oxo-Ade**$^{BZT}$G: conformer 1 (blue), conformer 2 (orange), conformer 3 (green), conformer 4 (red).

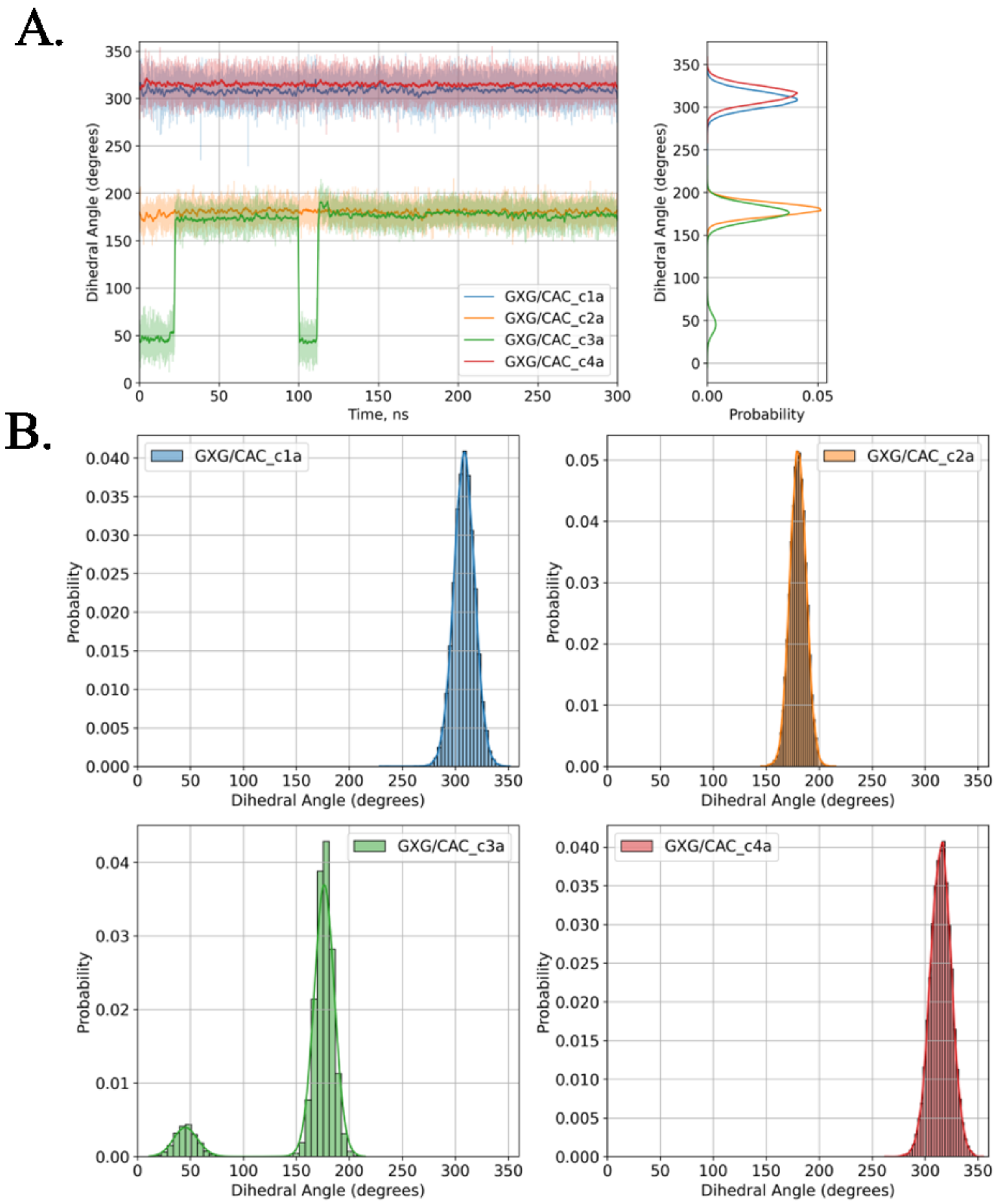

Figure S26. Evolution (A) and distribution (B) of dihedral angle d2 for four conformers of the **oxo-Ade**$^{BZT}$ nucleotide within the duplex with the central base pair triplet G**oxo-Ade**$^{BZT}$G/CA$^{anti}$C, where the 2'-deoxyadenosine nucleotide is in the anti conformation: conformer 1 (blue), conformer 2 (orange), conformer 3 (green), conformer 4 (red).

A.

B.

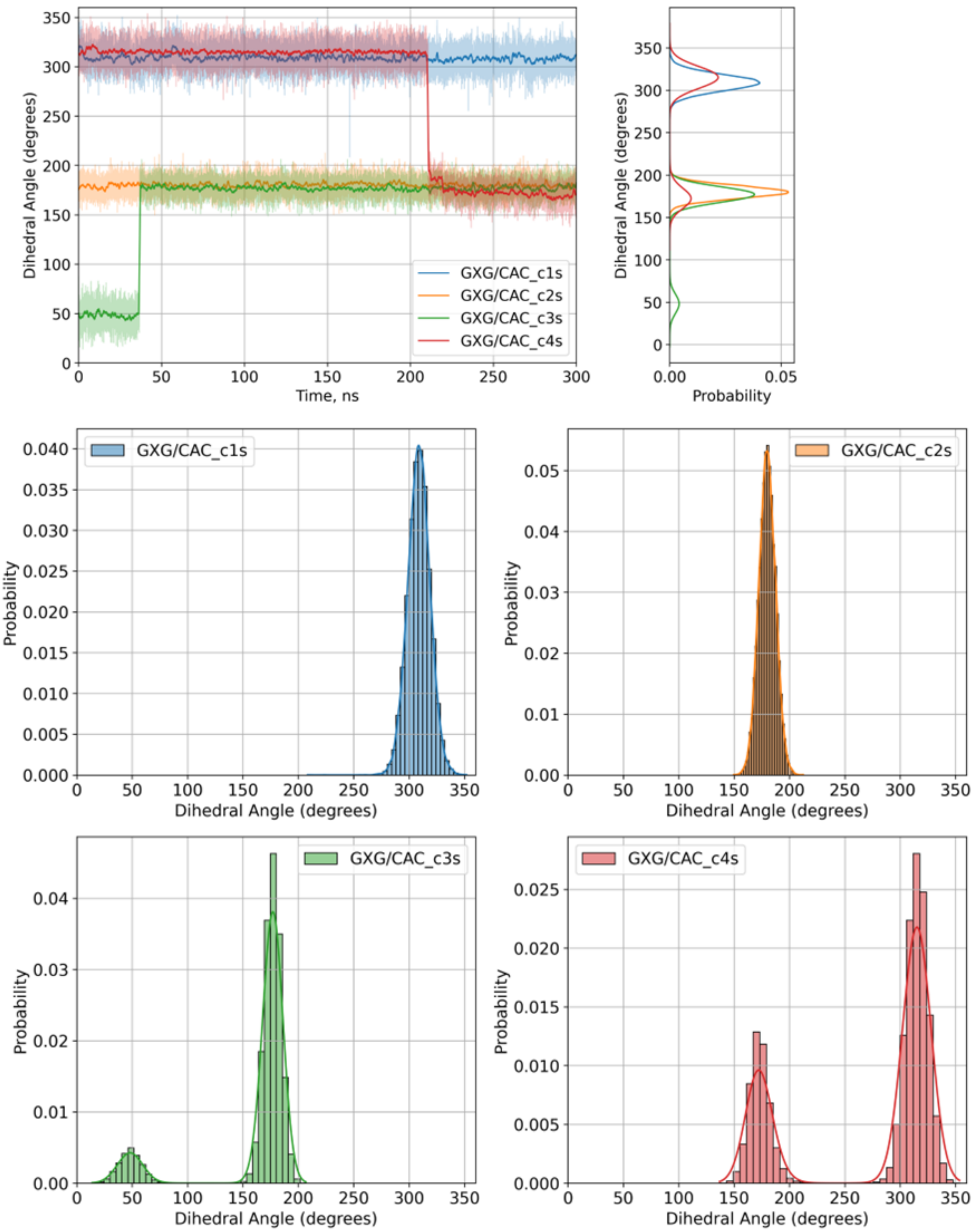

Figure S27. Evolution (A) and distribution (B) of dihedral angle d2 for four conformers of the **oxo-Ade**$^{BZT}$ nucleotide within the duplex with the central base pair triplet G**oxo-Ade**$^{BZT}$G/CA$^{syn}$C, where the 2'-deoxyadenosine nucleotide is in the syn conformation: conformer 1 (blue), conformer 2 (orange), conformer 3 (green), conformer 4 (red).

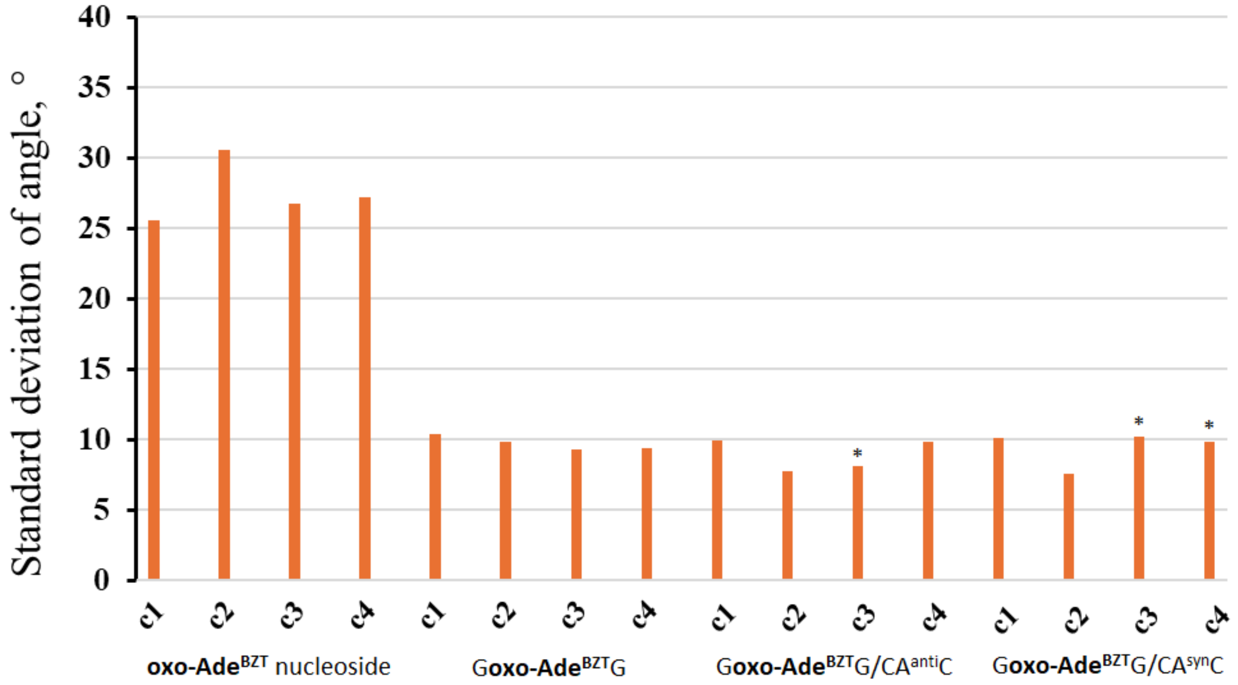

Figure S28. Standard deviation of d2 within the **oxo-Ade$^{BZT}$** nucleoside, ODN and duplexes for all four conformers. * since two stable states with different angles (Figure S27) were observed during the MD simulation, the position with the longest occupation time in the MD simulation was taken for the data; both values are presented in Table S8.

**Dihedral angle d3**

For all four conformers, the **oxo-Ade$^{BZT}$** nucleoside exhibits the high mobility along the N-glycosidic bond, represented by d3. The two peaks are observed with maxima of 100° and 280° for conformers 1 and 2 and 280° and 100° for conformers 3 and 4 (Figure S29). The distance between the maxima depending on the conformer is on average 180° for all four conformers. For ODN with the central triplet G**oxo-Ade$^{BZT}$**G, the mobility of d3 is significantly reduced compared to the **oxo-Ade$^{BZT}$** nucleoside and represented by one peak for all four conformers with a maximum of 125° for conformers 1 and 2, and 280° for conformers 3 and 4 (Figure S30). For the duplex with the central base pair triplet G**oxo-Ade$^{BZT}$**G/CA$^{anti}$C, where the 2'-deoxyadenosine nucleotide is in the anti conformation, the mobility of d3 is significantly reduced compared to the **oxo-Ade$^{BZT}$** nucleoside and represented by one peak for all four conformers with a maximum of 115° for conformer 1, 120° for conformer 2, 270° for conformer 3 and 235° for conformer 4 (Figure S31). For the duplex with the central base pair triplet G**oxo-Ade$^{BZT}$**G/CA$^{syn}$C, where the 2'-deoxyadenosine nucleotide is in the syn conformation, the mobility of d3 is represented by one peak for all four conformers with the same maximum as for the duplex with the central base pair triplet G**oxo-Ade$^{BZT}$**G/CA$^{anti}$C (Figure S32). For G**oxo-Ade$^{BZT}$**G/CA$^{anti}$C conformer 3 and G**oxo-Ade$^{BZT}$**G/CA$^{syn}$C conformers 3 and 4, an increase in the mobility of d3 is observed, which is most likely due to a 100-120° rotation along the single bond represented by d2 (Figure S33).

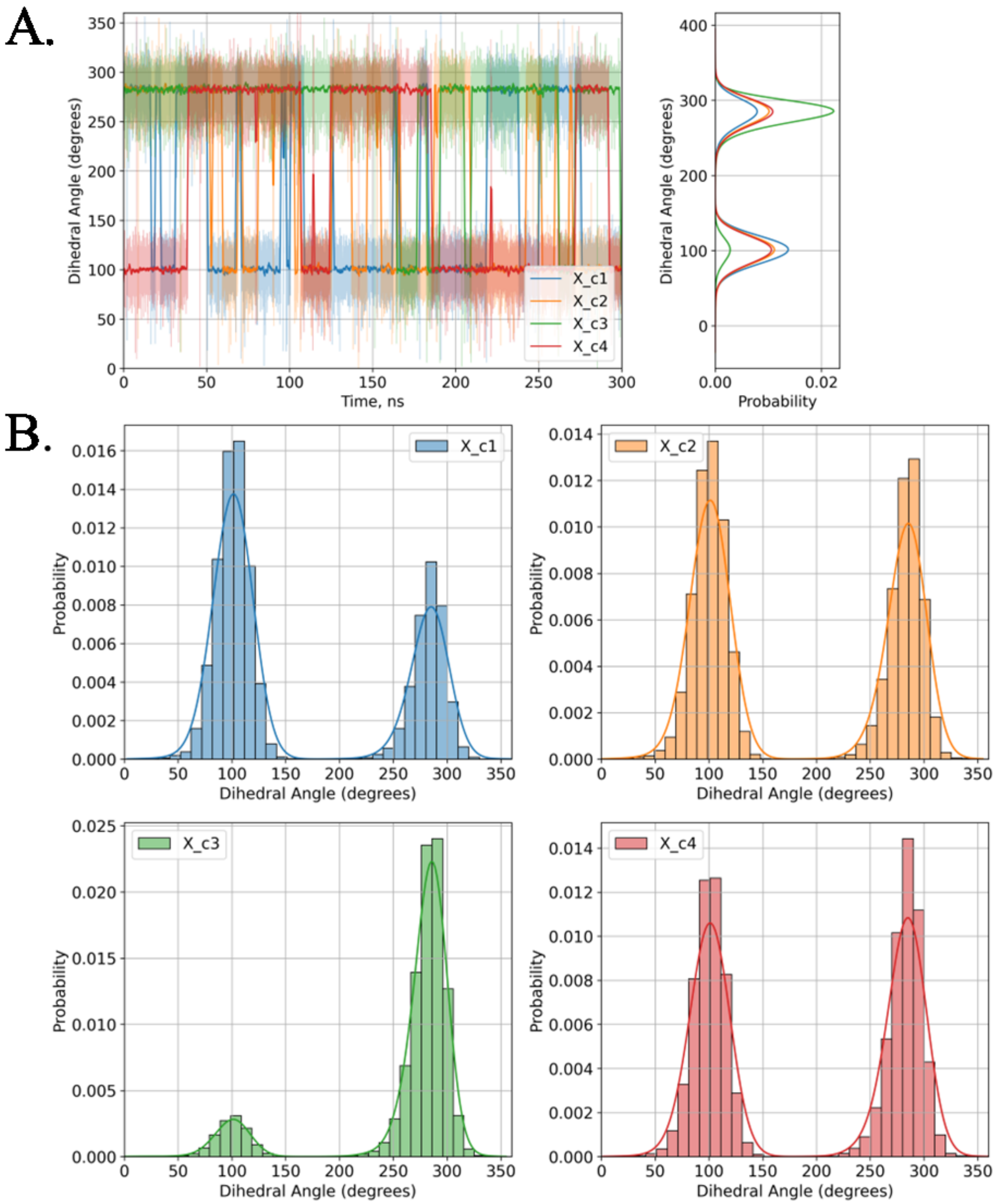

Figure S29. Evolution (A) and distribution (B) of dihedral angle d3 for four conformers of the **oxo-Ade**$^{BZT}$ nucleoside: conformer 1 (blue), conformer 2 (orange), conformer 3 (green), conformer 4 (red).

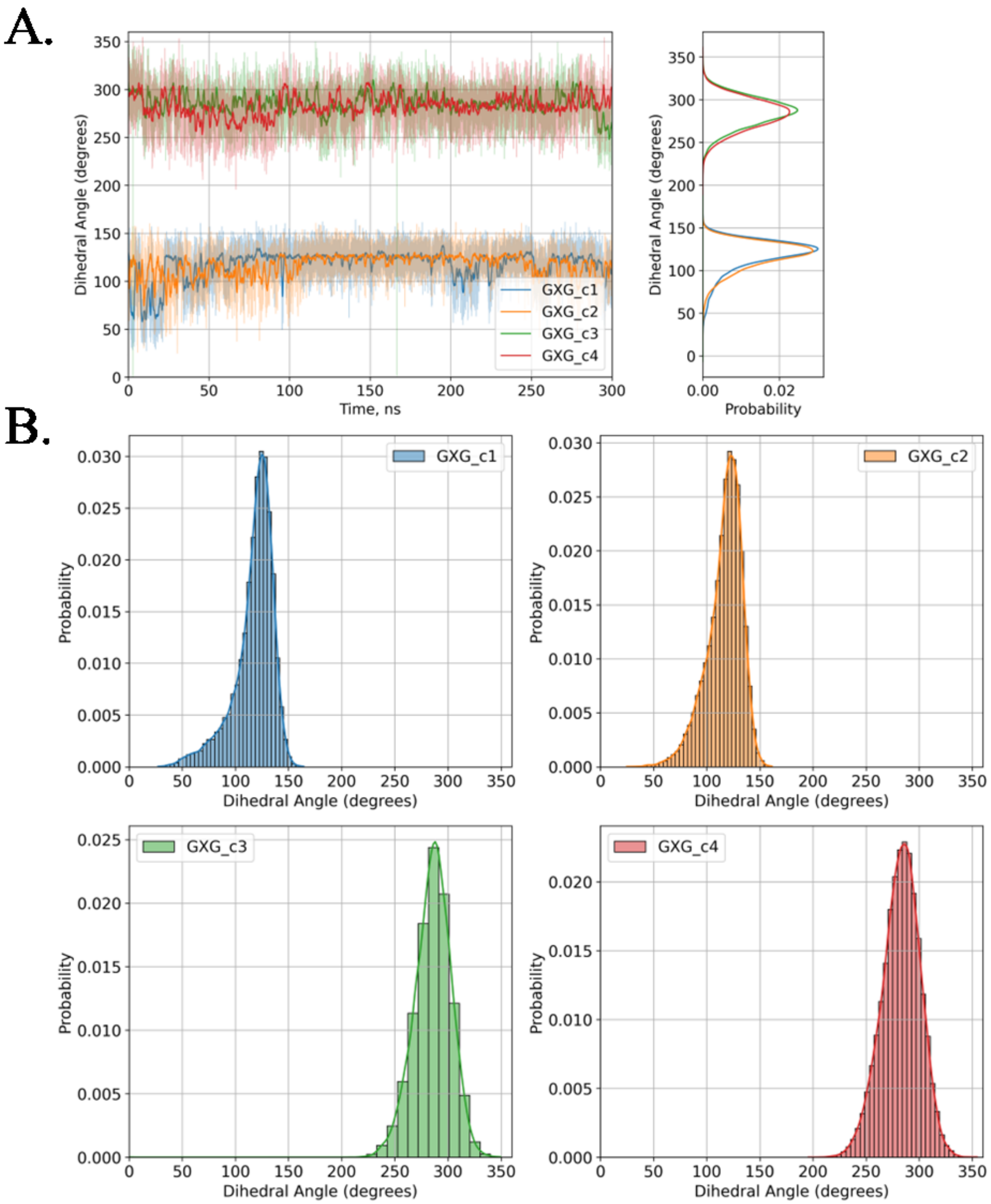

Figure S30. Evolution (A) and distribution (B) of dihedral angle d3 for four conformers of the **oxo-Ade**$^{BZT}$ nucleotide within ODN with the central triplet G**oxo-Ade**$^{BZT}$G: conformer 1 (blue), conformer 2 (orange), conformer 3 (green), conformer 4 (red).

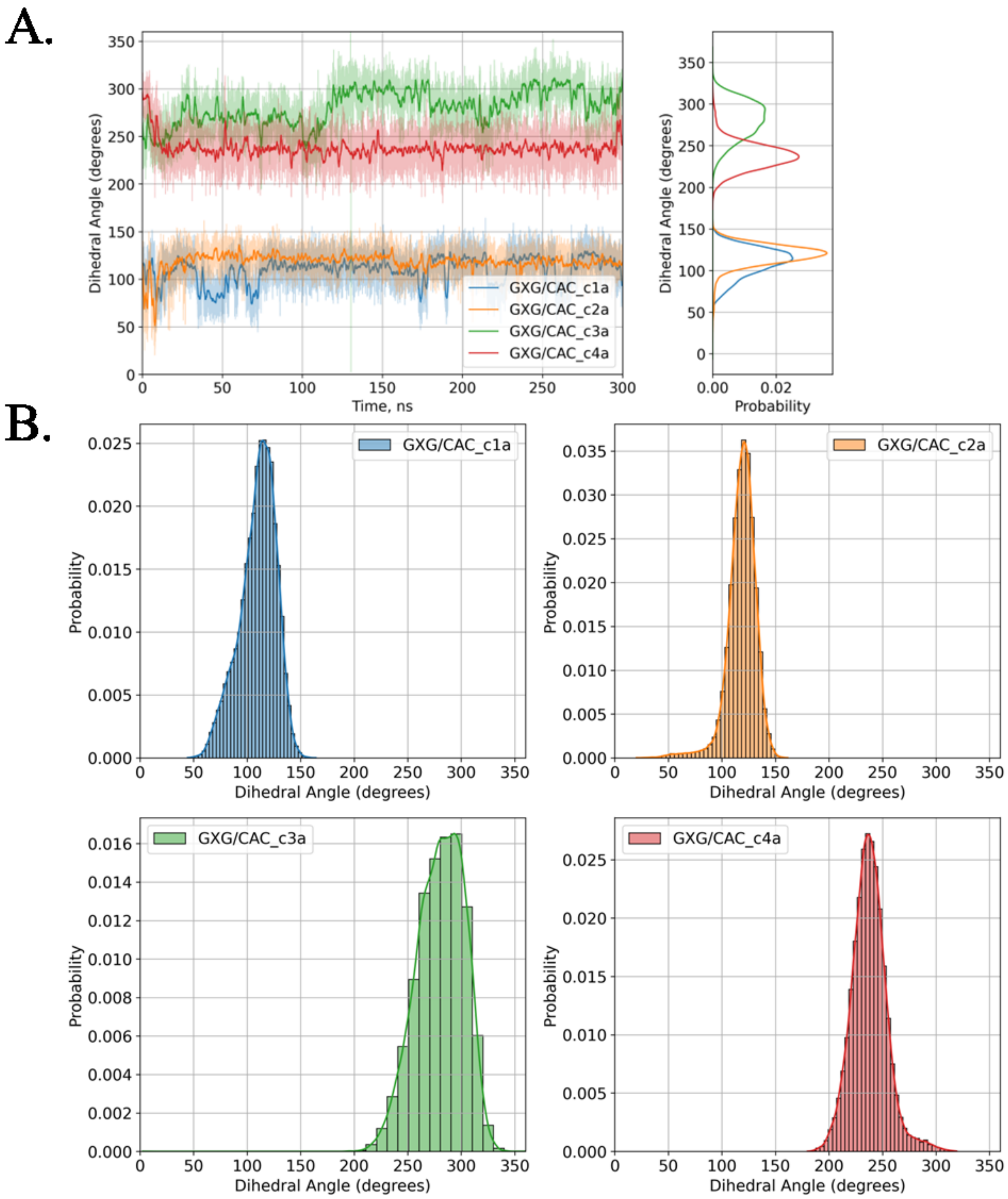

Figure S31. Evolution (A) and distribution (B) of dihedral angle d3 for four conformers of the **oxo-Ade**$^{BZT}$ nucleotide within the duplex with the central base pair triplet G**oxo-Ade**$^{BZT}$G/CA$^{anti}$C, where the 2'-deoxyadenosine nucleotide is in the anti conformation: conformer 1 (blue), conformer 2 (orange), conformer 3 (green), conformer 4 (red).

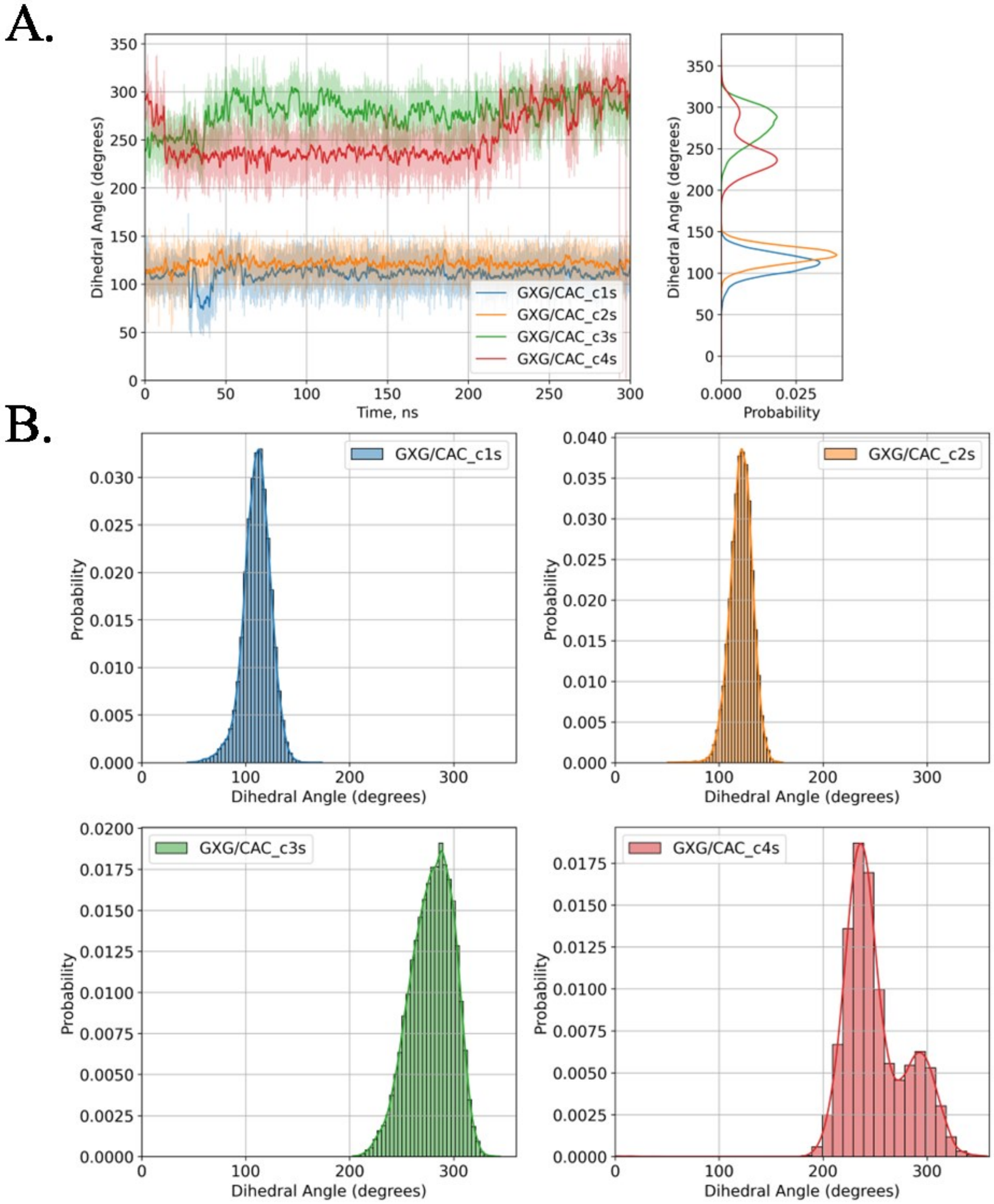

Figure S32. Evolution (A) and distribution (B) of dihedral angle d3 for four conformers of the **oxo-Ade**$^{BZT}$ nucleotide within the duplex with the central base pair triplet G**oxo-Ade**$^{BZT}$G/CA$^{syn}$C, where the 2'-deoxyadenosine nucleotide is in the syn conformation: conformer 1 (blue), conformer 2 (orange), conformer 3 (green), conformer 4 (red).

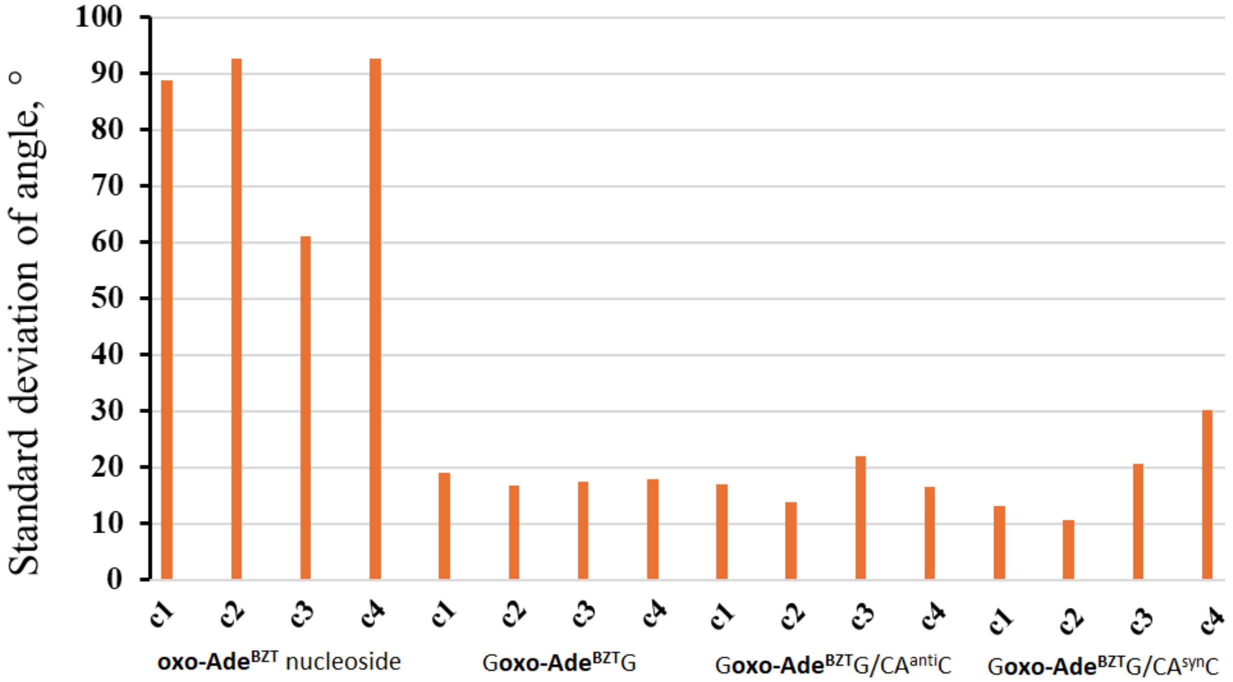

Figure S33. Standard deviation of d3 within the **oxo-Ade**$^{BZT}$ nucleoside, ODN and duplexes for all four conformers.

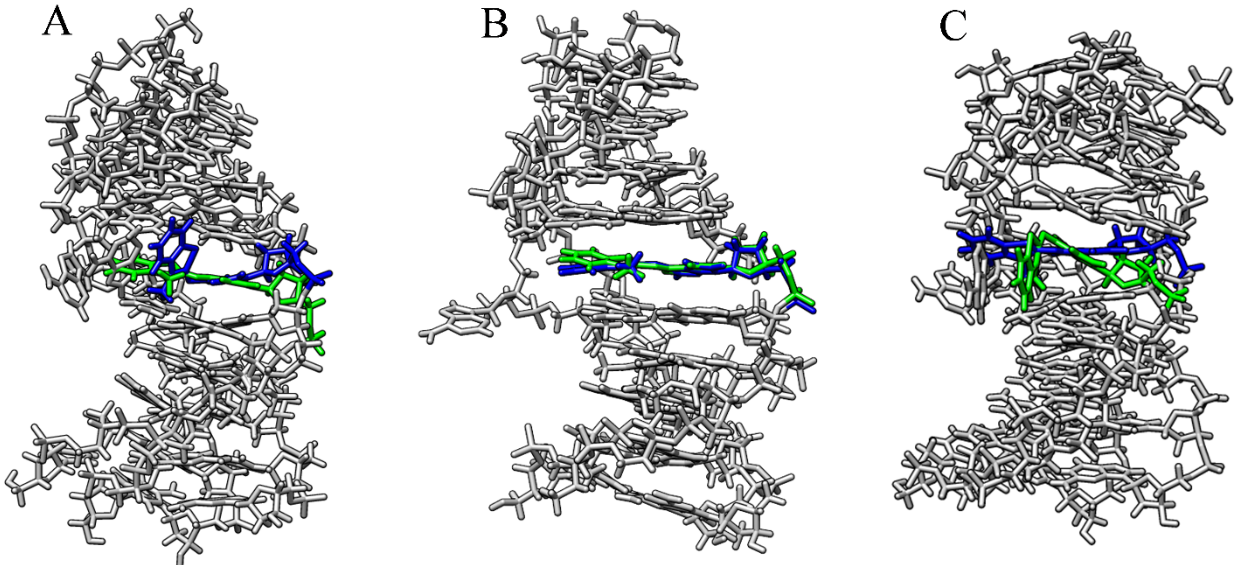

Figure S34. Superposition of the most represented structures and the structures with the largest deviations in the dihedral angles d1, d2, and d3: G**oxo-Ade**$^{BZT}$G/CA$^{anti}$C conformer 3 (A), G**oxo-Ade**$^{BZT}$G/CA$^{syn}$C conformer 3 (B), G**oxo-Ade**$^{BZT}$G/CA$^{syn}$C conformer 4 (C). The green and blue structures exhibit the highest and lowest RMSD for the dihedral angle relative to the total MD simulation time. The modification is in the specified states for more than 15 ns.

Table S8. Standard deviation of dihedral angles (in degrees) for all four conformers

| | | d1 | d2 | d3 |
|---|---|---|---|---|
| **oxo-Ade**$^{BZT}$ nucleoside | conformer 1 | 17.2 | 25.6 | 88.8 |
| | conformer 2 | 21.5 | 30.6 | 92.6 |
| | conformer 3 | 17.2 | 26.7 | 61.1 |
| | conformer 4 | 17.5 | 27.2 | 92.7 |
| G**oxo-Ade**$^{BZT}$G | conformer 1 | 10.1 | 10.4 | 19.0 |
| | conformer 2 | 9.5 | 9.8 | 16.7 |
| | conformer 3 | 10.8 | 9.3 | 17.3 |
| | conformer 4 | 11.1 | 9.4 | 17.9 |
| G**oxo-Ade**$^{BZT}$G/CA$^{anti}$C | conformer 1 | 7.8 | 10.0 | 17.0 |
| | conformer 2 | 7.7 | 7.7 | 13.7 |
| | conformer 3 | 11.7 | 8.2 (10.4)* | 22.0 |
| | conformer 4 | 10.6 | 9.9 | 16.5 |
| G**oxo-Ade**$^{BZT}$G/CA$^{syn}$C | conformer 1 | 7.9 | 10.1 | 13.1 |
| | conformer 2 | 7.7 | 7.6 | 10.6 |
| | conformer 3 | 9.5 | 7.5 (10.2)* | 20.6 |
| | conformer 4 | 10.7 (11.8)* | 9.8 (9.6)* | 30.2 |

* Two rotated states are observed during the MD simulation, the state with the longest occupation time in the MD simulation is used for correct calculations.

Table S9. Averaged standard deviation of dihedral angles (in degrees) for all four conformers

| | d1 | d2 | d3 |
|---|---|---|---|
| **oxo-Ade**$^{BZT}$ nucleoside | 18.4 ± 2.1 | 27.5 ± 2.2 | 83.8 ± 15.2 |
| G**oxo-Ade**$^{BZT}$G | 10.4 ± 0.7 | 9.7 ± 0.5 | 17.7 ± 1.0 |
| G**oxo-Ade**$^{BZT}$G/CAC | 9.2 ± 1.6 | 9.2 ± 1.2 | 18.0 ± 8.8 |

The large error value of d3 for the duplex for G**oxo-Ade**$^{BZT}$G/CA$^{anti}$C conformer 3 and G**oxo-Ade**$^{BZT}$G/CA$^{syn}$C conformers 3 and 4 is associated with its rotation toward the most energetically favorable state relative to the bond represented by d2 (Table S9). This suggests that these conformers are very unlikely present in solution. At the same time, for ODN and duplexes, a significant decrease in the mobility around the N-glycosidic bond is observed due to stacking interactions of the appended heterocyclic system with flanking nucleobases in the same and opposite strands.

### 10. NMR spectra

5',3'-*O*-acetyl-2'-deoxy-7,8-dihydro-8-oxoadenosine **2**

$^{1}$H spectrum

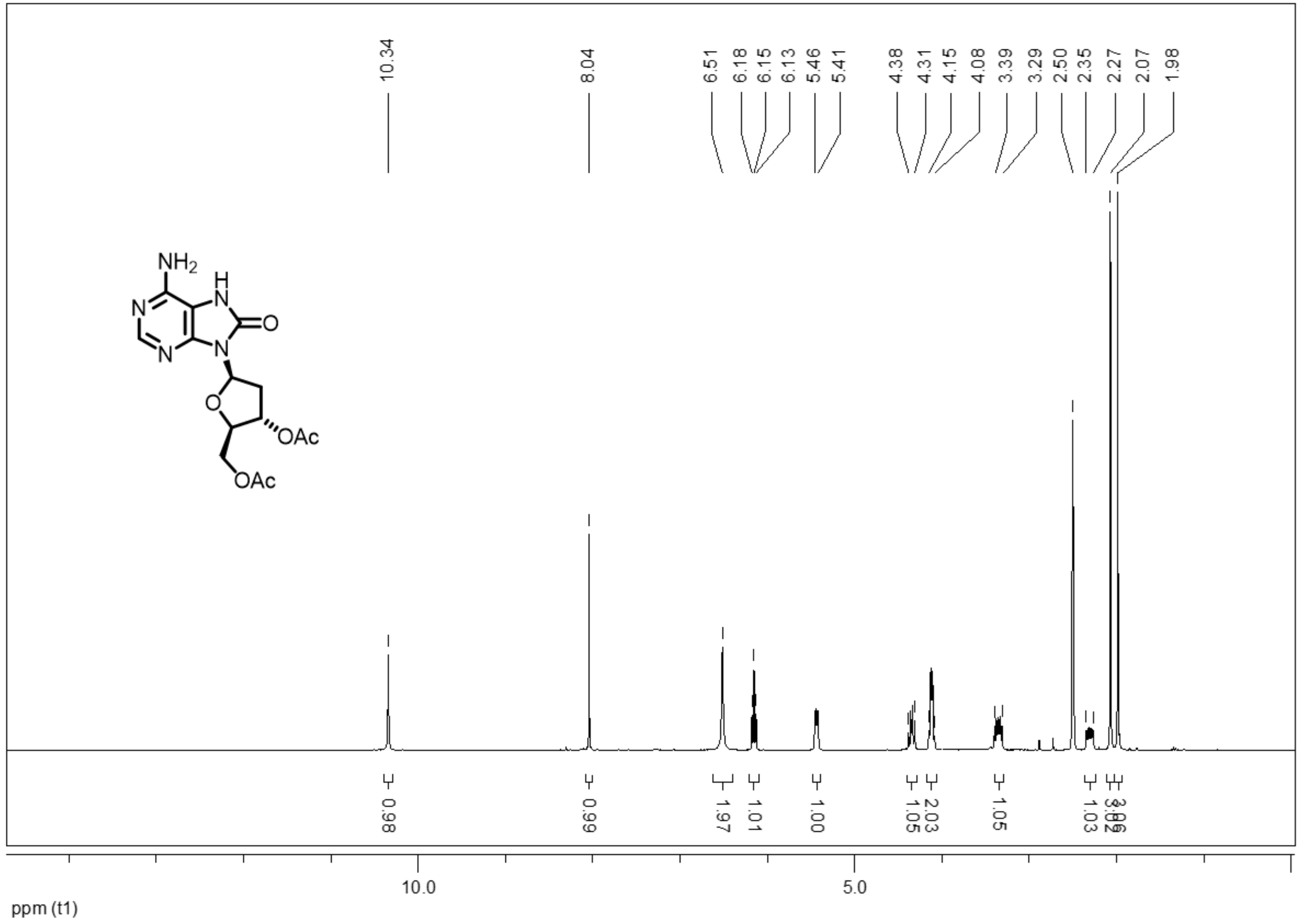

$^{13}C$ spectrum

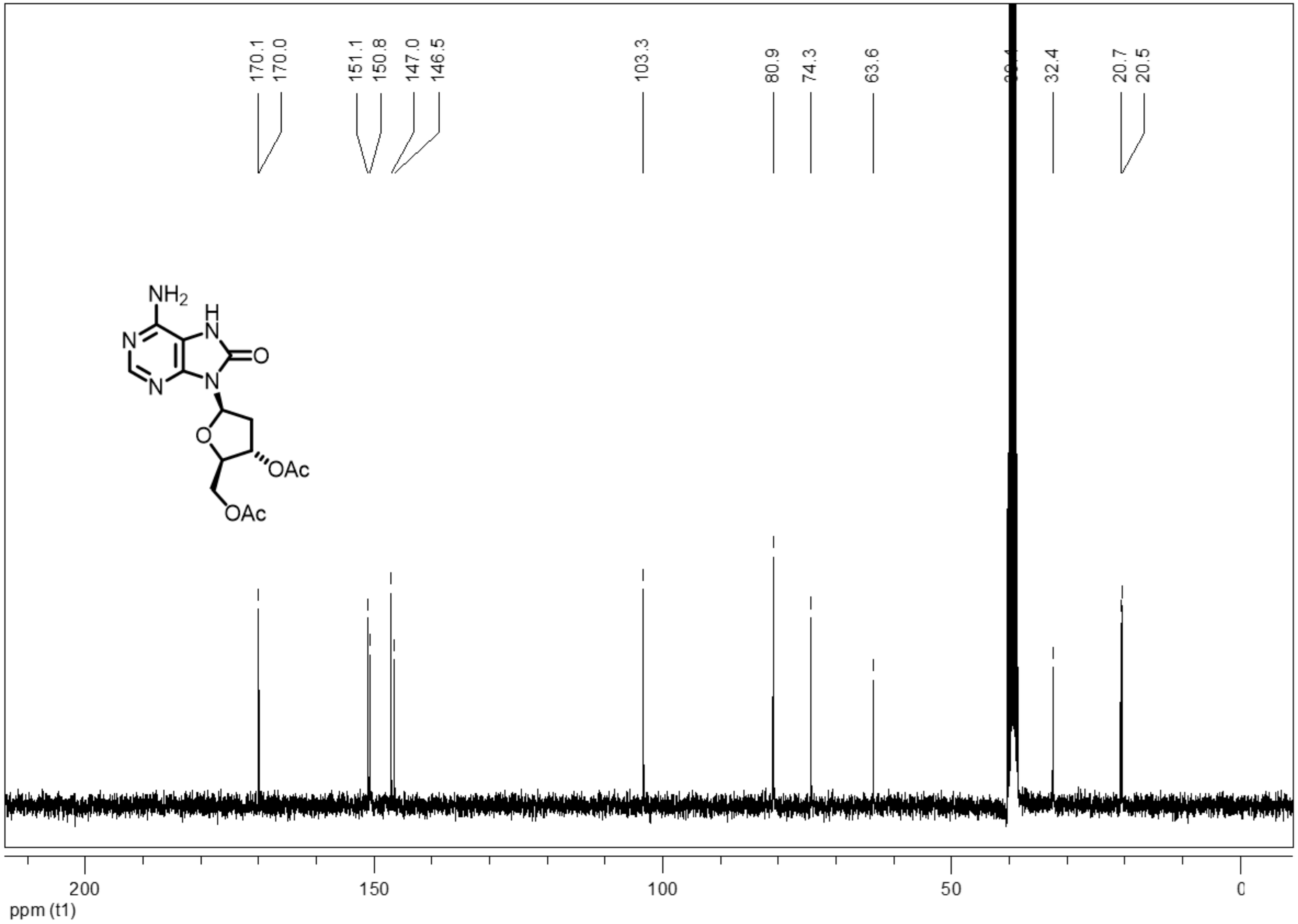

5',3'-*O*-acetyl-2'-deoxy-6-(3-methylbenzo[d]thiazol-2(3H)-ylidene)-7,8-dihydro-8-oxoadenosine **3**

$^{1}$H spectrum

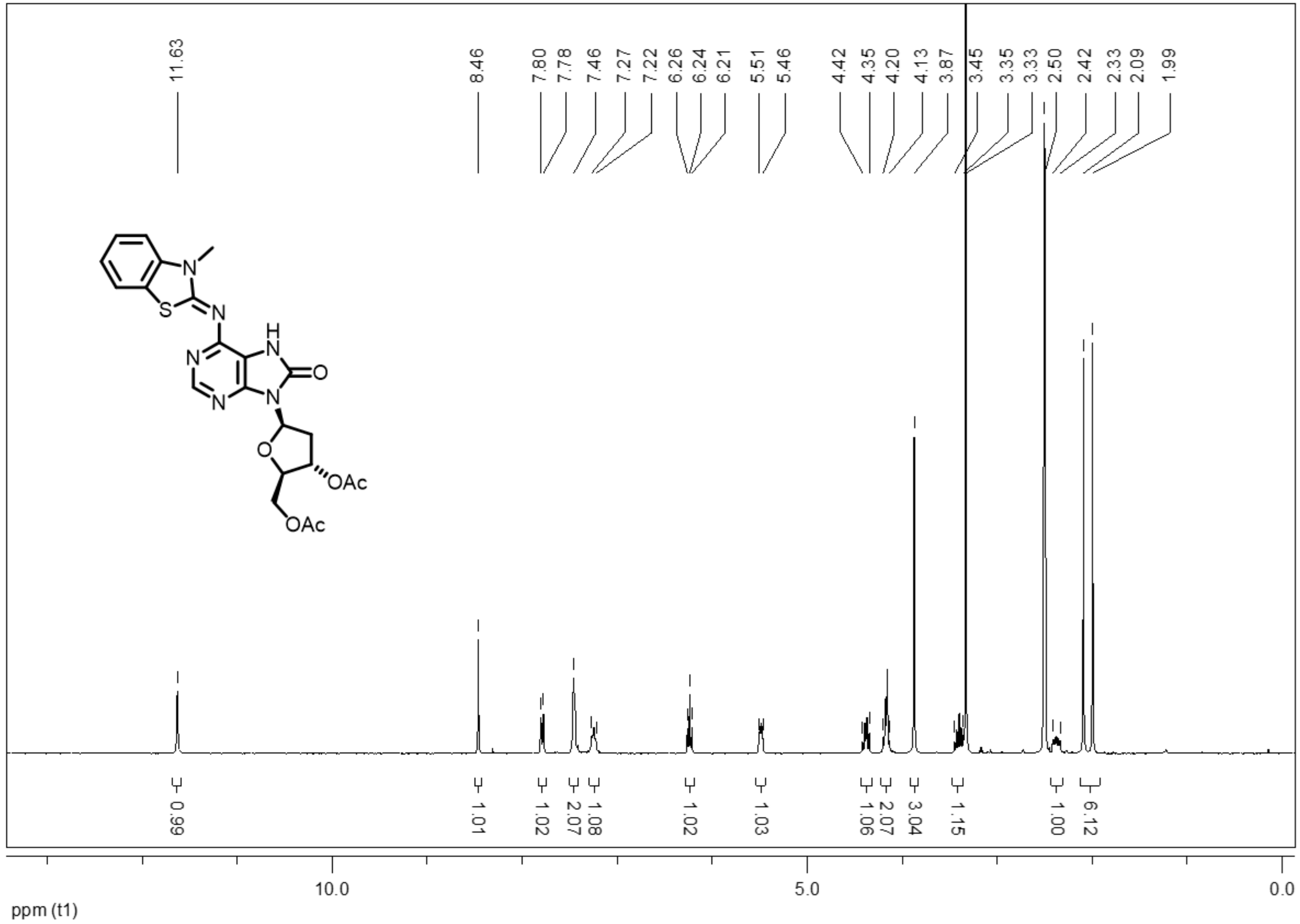

$^{13}C$ spectrum

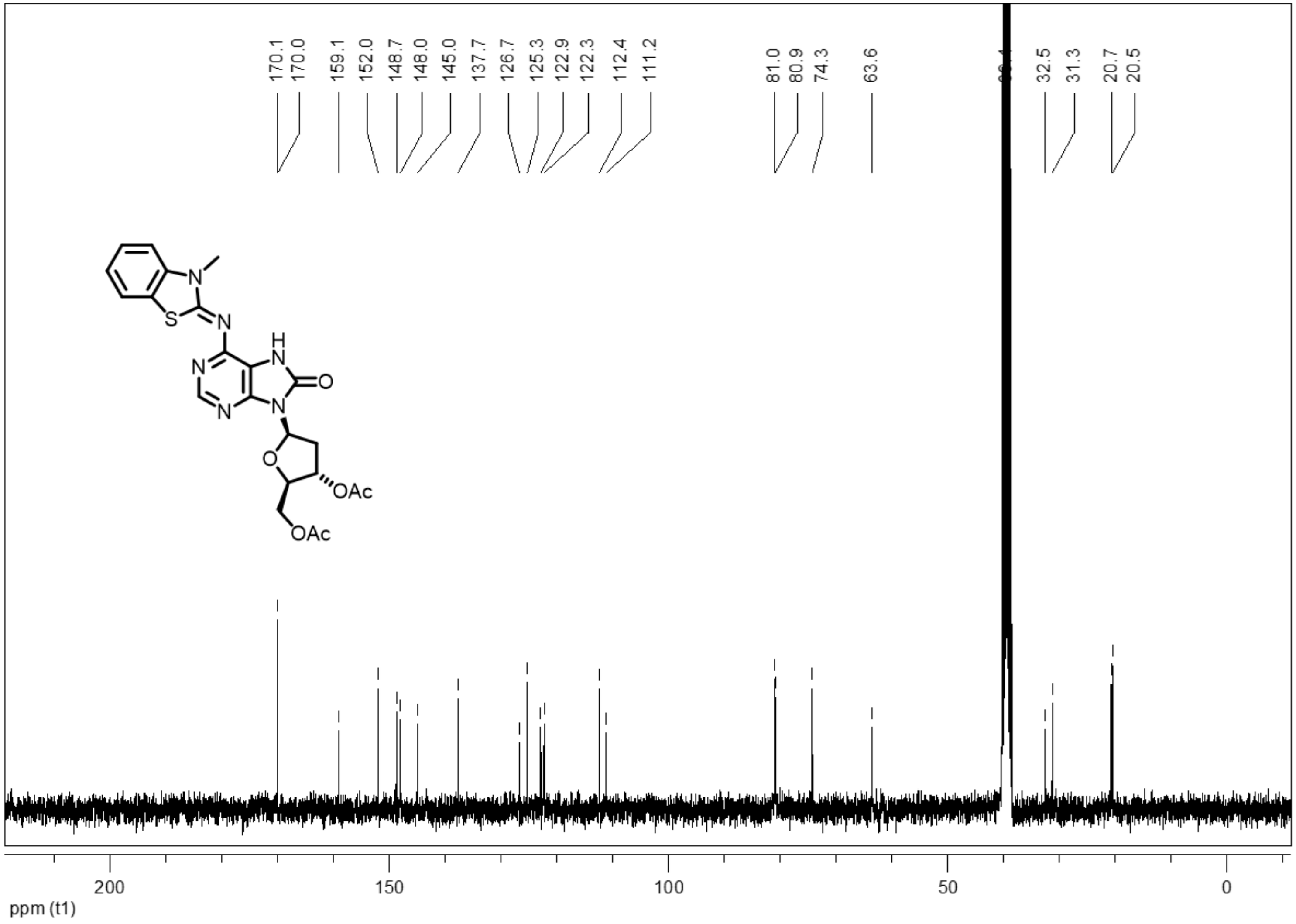

2'-deoxy-7,8-dihydro-8-oxo-6-(3-methylbenzo[d]thiazol-2(3H)-ylidene)adenosine **4**

$^{1}$H spectrum

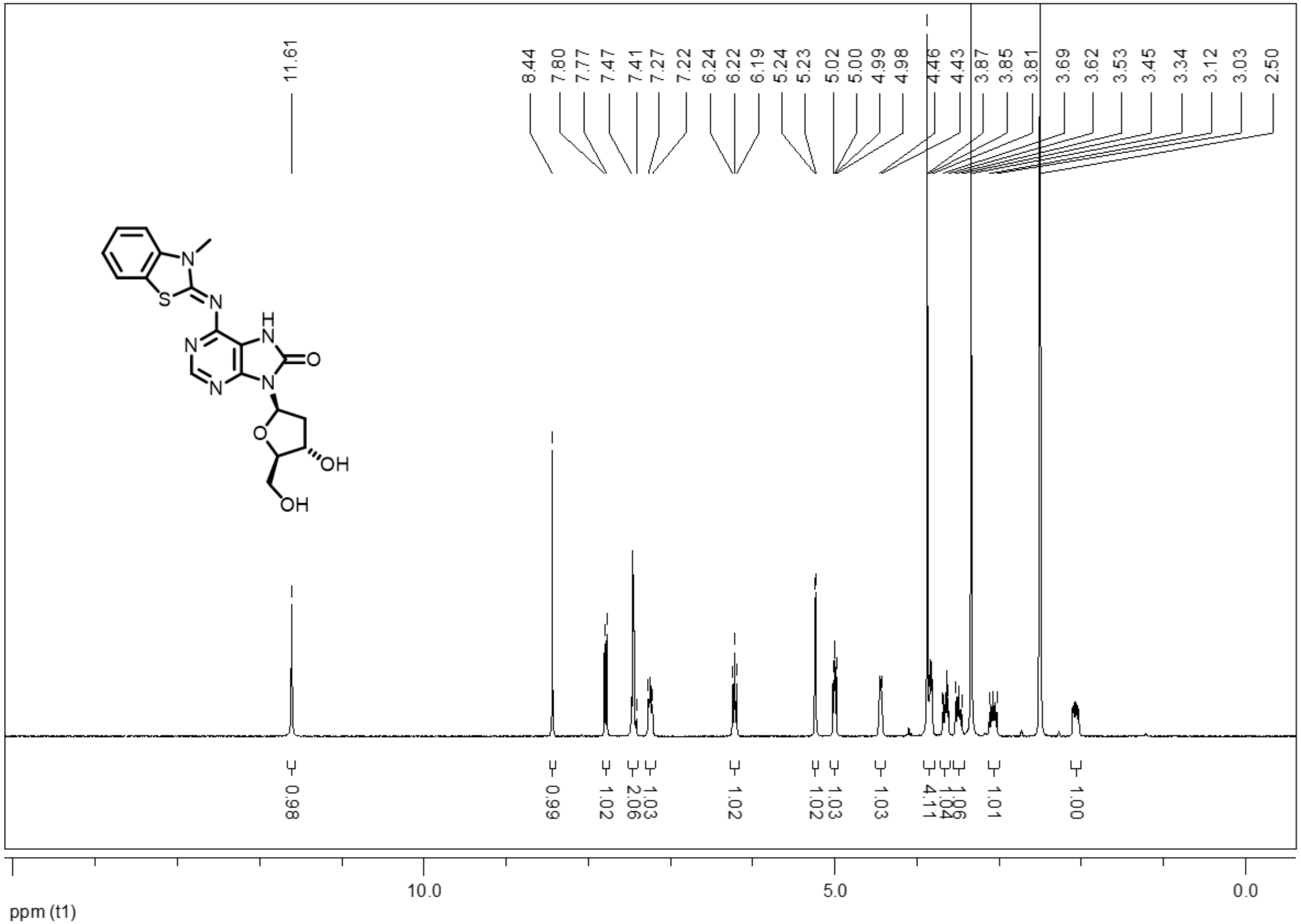

$^{13}$C spectrum

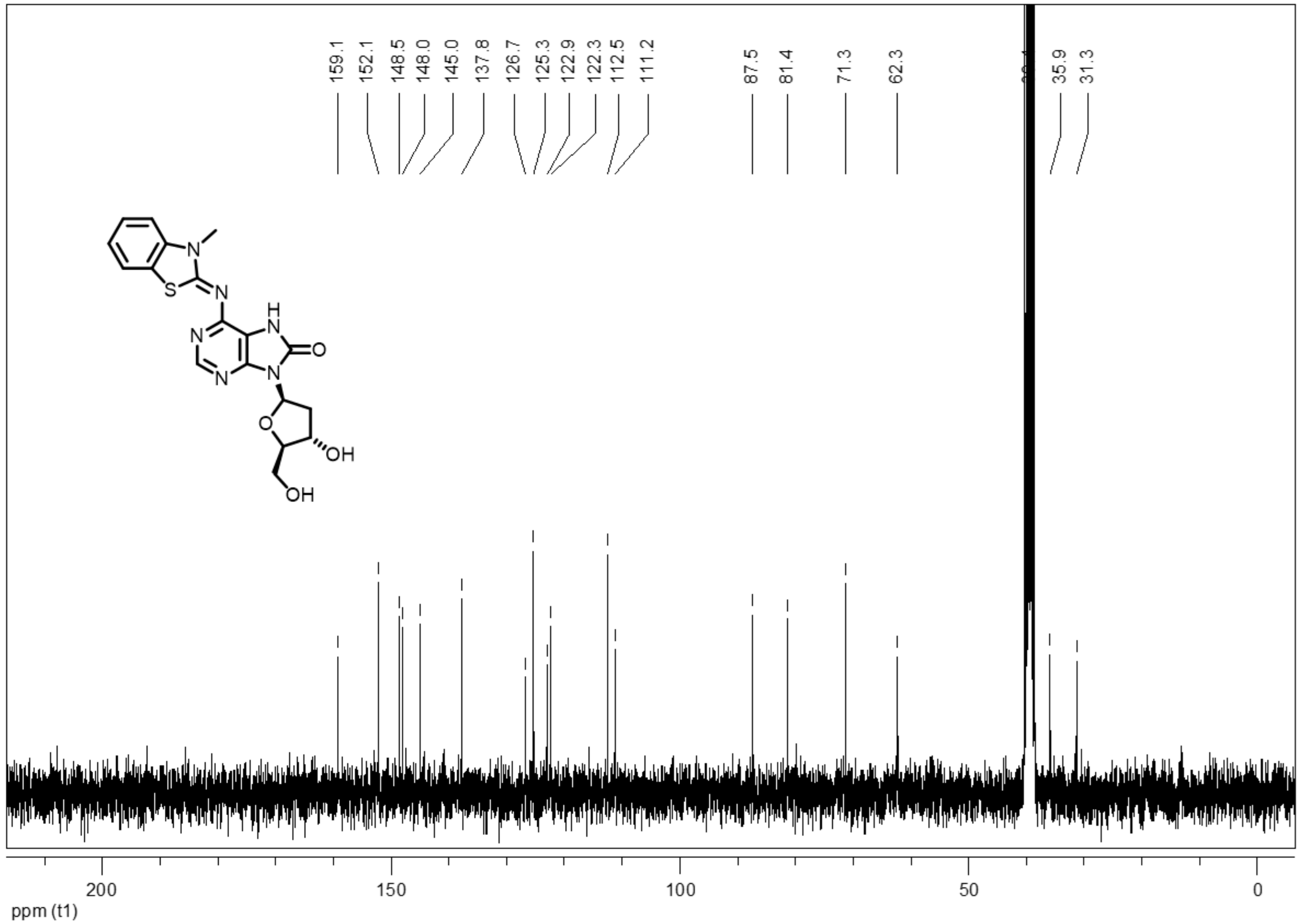

159.1
152.1
148.5
148.0
145.0
137.8
126.7
125.3
122.9
122.3
112.5
111.2
87.5
81.4
71.3
62.3
35.9
31.3
N
S
N
H
N
N
O
N
N
O
OH
OH
200
150
100
50
0
ppm (t1)

5'-*O*-(4,4'-Dimethoxytrityl)-2'-deoxy-7,8-dihydro-8-oxo-6-(3-methylbenzo[d]thiazol-2(3H)-ylidene)adenosine **5**

$^{1}$H spectrum

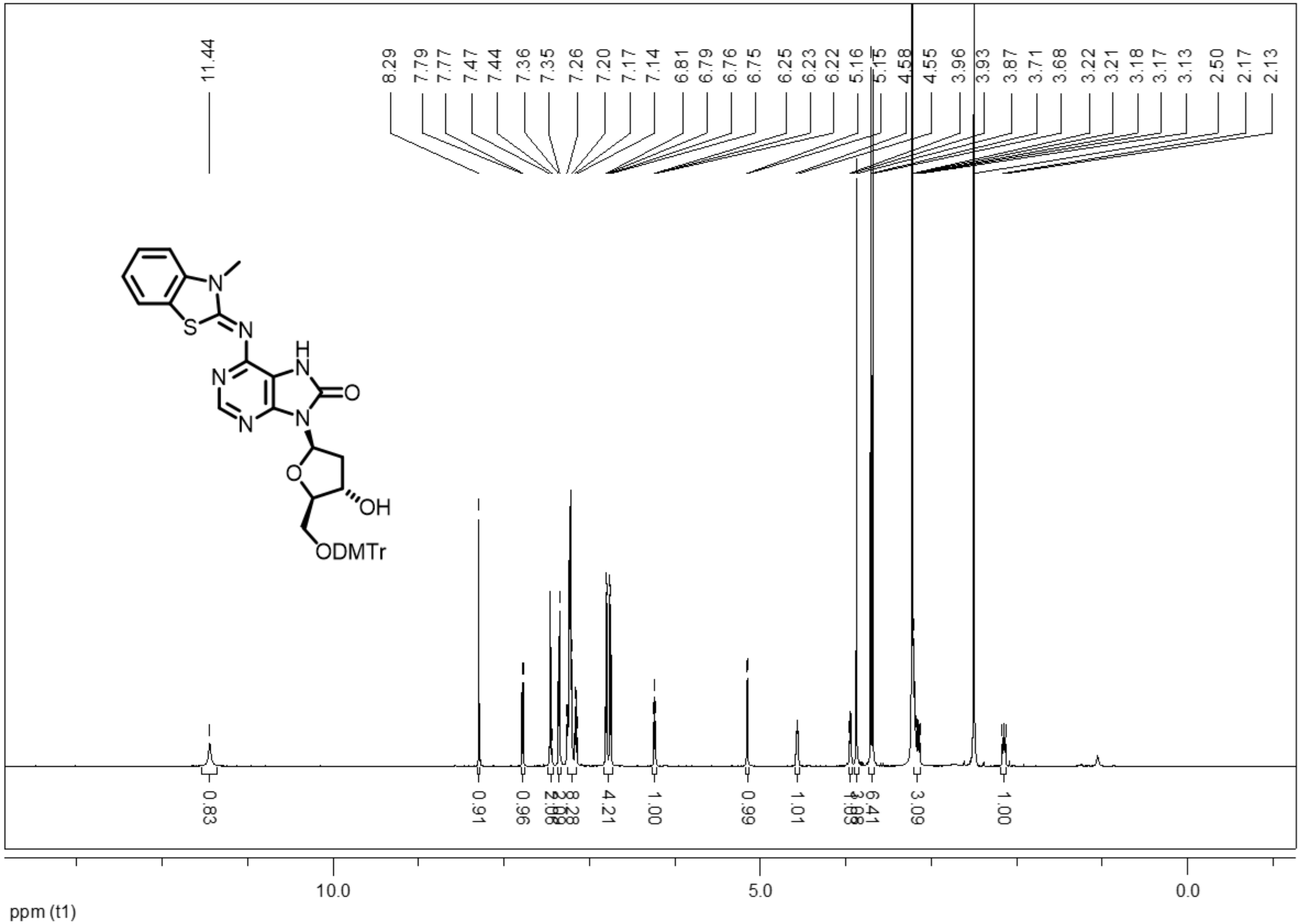

$^{13}$C spectrum

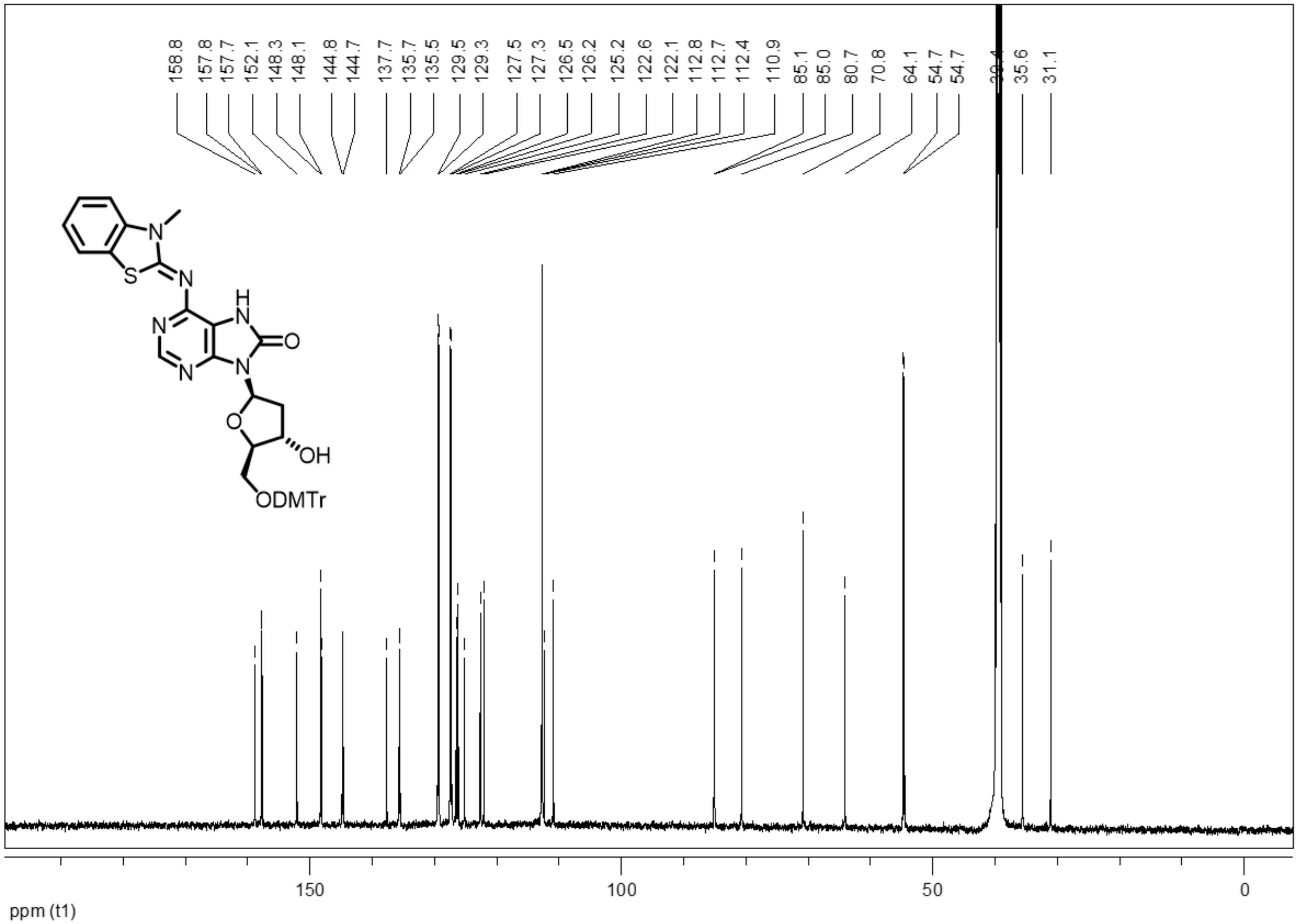
158.8
157.8
157.7
152.1
148.3
148.1
144.8
144.7
137.7
135.7
135.5
129.5
129.3
127.5
127.3
126.5
126.2
125.2
122.6
122.1
112.8
112.7
112.4
110.9
85.1
85.0
80.7
70.8
64.1
54.7
54.7
35.6
31.1
N
S
N
H
N
N
O
N
N
O
OH
ODMTr
150
100
50
0
ppm (t1)

5'-*O*-(4,4'-Dimethoxytrityl)-3'-*O*-((2-cyanoethyl)-*N*,*N*-diisopropylphosphoramidite)-2'-deoxy-7,8-dihydro-8-oxo-6-(3-methylbenzo[d]thiazol-2(3H)-ylidene)adenosine **6**

$^{1}$H spectrum

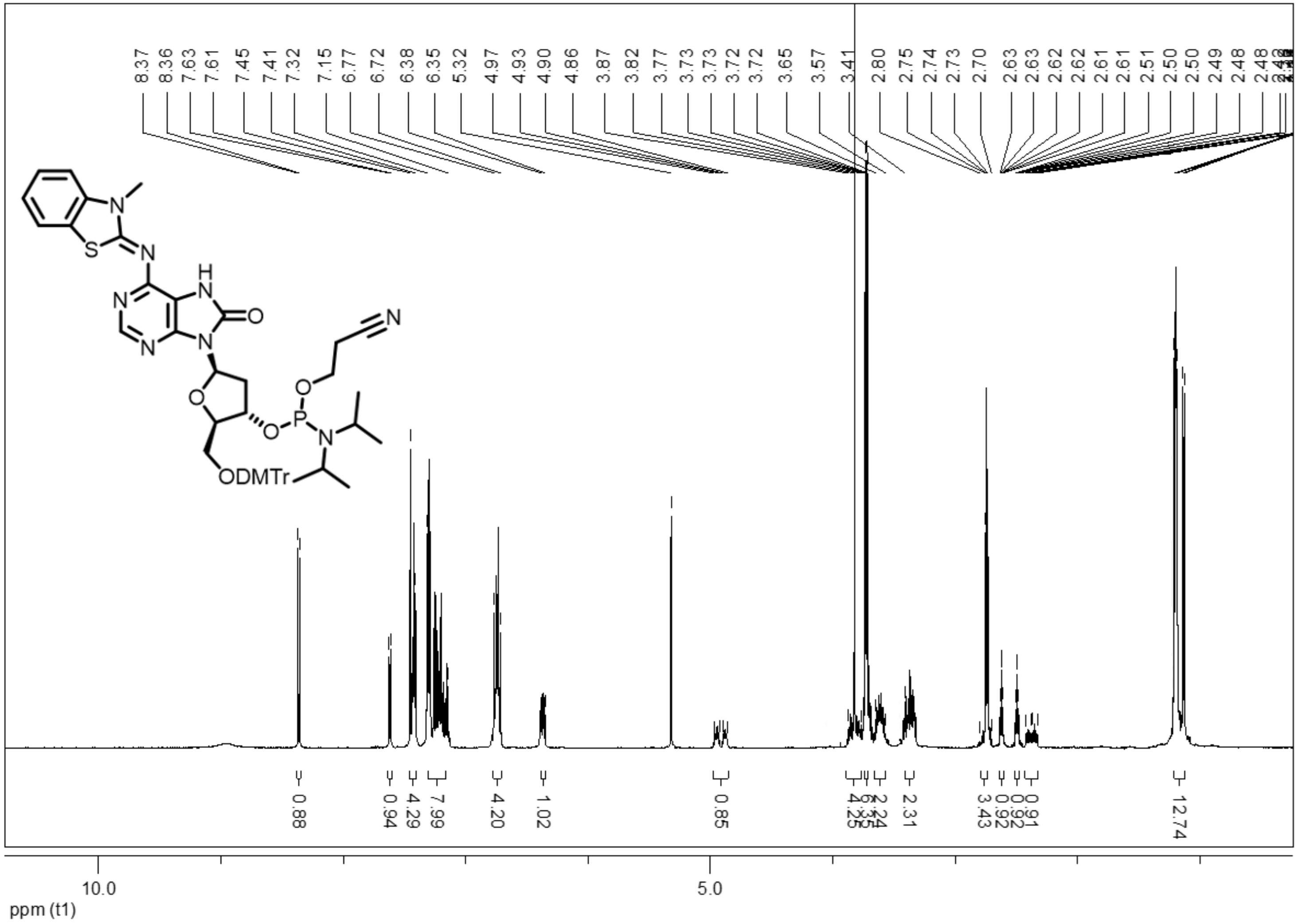

$^{31}$P spectrum

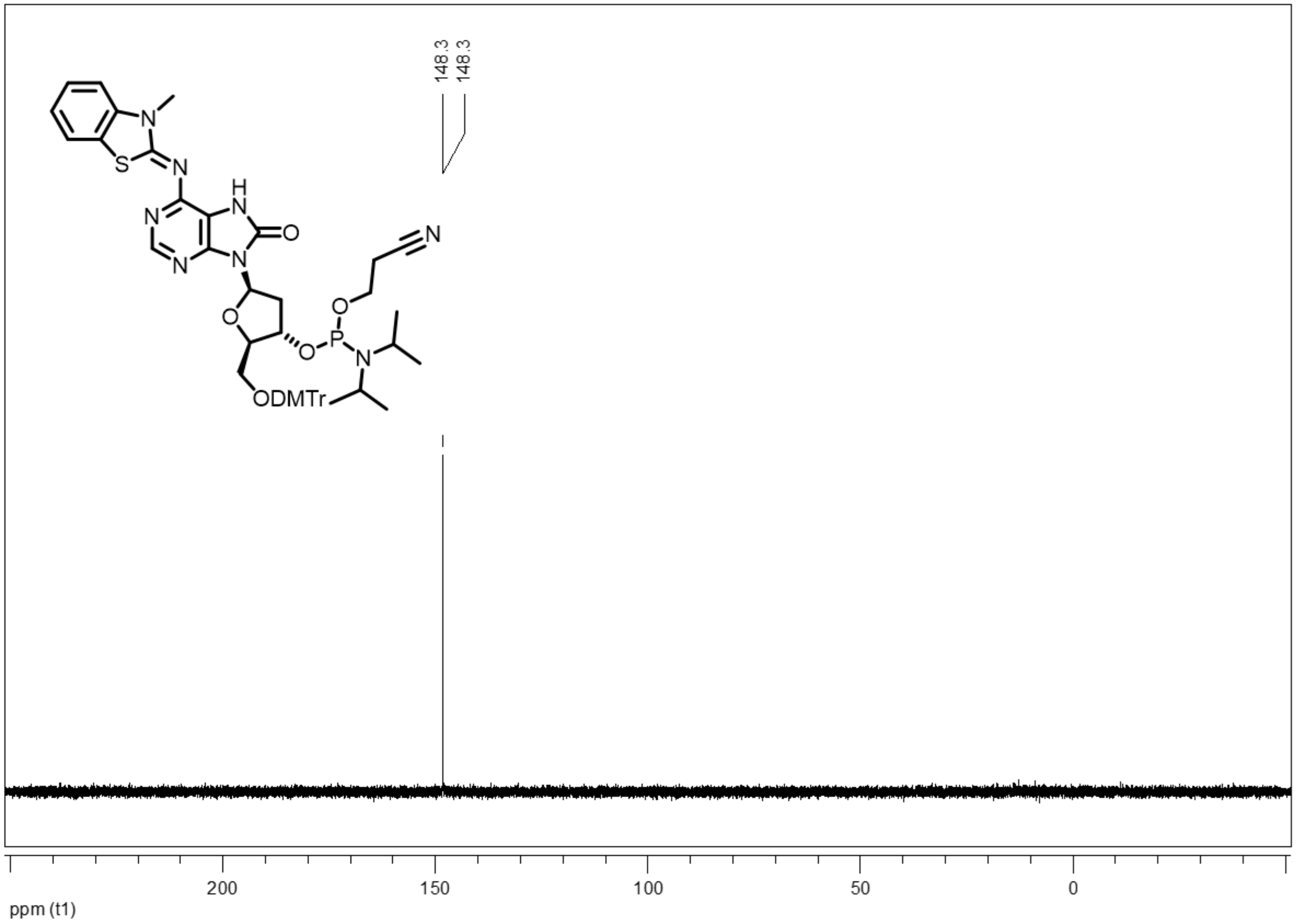
148.3
148.3
N
S
N
H
N
N
O
N
N
O
N
O
P
O
N
ODMTr
200
150
100
50
0
ppm (t1)
